# A model for the gradual evolution of dioecy and heterogametic sex determination

**DOI:** 10.1101/2023.03.24.534076

**Authors:** Thomas Lesaffre, John R. Pannell, Charles Mullon

## Abstract

In plants with separate sexes, the development of an individual as male or female is controlled by a dominant allele at a sex-determining locus - the fundamental basis of XY and ZW systems. The many independent transitions from hermaphroditism to dioecy that have taken place in flowering plants must therefore often have entailed the emergence of such a locus. One proposition is that this evolution occurs in two steps, with the initial invasion of a male-sterility mutation at one locus followed by mutations causing female sterility at a second closely linked locus. Here, we show how dioecy with heterogametic sex determination can also emerge in a gradual adaptive process, involving the co-evolution of resource allocation to different sexual functions jointly with its genetic architecture. Our model reveals that whether an XY or a ZW system evolves depends on the mating system of the ancestral hermaphrodites as well as the trade-off they face between allocation to male and female functions. In particular, the evolution of dioecy in response to selection to avoid selfing and inbreeding depression favours the emergence of XY systems, which characterise the vast majority of dioecious flowering plants. Selection favouring female specialisation also favours XY over ZW sex determination. Taken together, our results throw new light on the possible origins of dioecy from hermaphroditism by revealing a hitherto unrecognised link between the ecology and economics of sex allocation and the genetic basis of sex determination.

## Introduction

Many plants have evolved separate sexes (or dioecy) from hermaphroditism (Charlesworth, 1985; Renner, 2014; Henry et al., 2018). Explanations for these repeated transitions have come from two types of models. First, sex allocation theory uses optimality arguments (*sensu* Parker and Maynard Smith 1990) to identify conditions under which natural selection favours individuals allocating all their resources to one sexual function over those allocating to both (Charnov et al., 1976; Charnov, 1982). Whether this happens depends on how resource allocation affects fitness, which may be influenced by a number of ecological and physiological factors (Givnish, 1982; Lloyd, 1982; Renner and Ricklefs, 1995; Freeman et al., 1997; Pannell and Jordan, 2022; Masaka and Takada, 2023). The joint effect of these factors is usefully summarised in male and female ‘gain curves’, which relate fitness gained through each sexual function to the amount of resources allocated to it (Charnov et al., 1976). Accelerating gain curves indicate increasing returns on investment and therefore an advantage to sexual specialisation (Charnov et al., 1976; Charnov, 1982). Second, population genetic arguments have revealed that dioecy may also evolve as a mechanism to avoid self-fertilisation and inbreeding depression (Charlesworth and Charlesworth, 1978a,b, 1981). When inbreeding depression is strong, a population of partially self-fertilising hermaphrodites can be invaded by a mutation causing partial or full male-sterility (i.e., by females) because male-sterility reduces selfing (Charlesworth and Charlesworth, 1978a,b). In turn, the establishment of females favours the invasion of female-sterility mutations, converting hermaphrodites into males.

Ultimately, whether dioecy evolves under favourable conditions will depend on the genetic architecture of sex (Charlesworth, 1999; Beukeboom and Perrin, 2014). Because sex is often determined at a polymorphic locus at which one sex is heterozygous (e.g., XY or ZW, in the case of male or female heterogamety, respectively) and the other is homozygous (XX or ZZ females and males, Ming et al., 2011; Bachtrog et al., 2014; Beukeboom and Perrin, 2014), the transition from hermaphroditism to dioecy must in many cases have involved the emergence of such a locus. While optimality models rely on the ‘phenotypic gambit’ and are mute to genetic details (Charnov et al., 1976; Charnov, 1982), population genetic models are explicit about the genetic architecture of sex determination. These models indicate that sex-determining loci may evolve through the invasion of an initial male-sterility mutation at a first locus, followed by one or more mutations causing female sterility at a second, closely linked locus (the ‘two-gene model’, Charlesworth and Charlesworth, 1978a,b, 1981; Charlesworth, 1991; Charlesworth et al., 2005; Beukeboom and Perrin, 2014; Olito and Connallon, 2019). Depending on whether that initial male sterility allele is assumed to be fully recessive or fully dominant, this scenario predicts either XY or ZW sex determination, respectively.

Here, using a model drawing on optimality and population genetic arguments jointly, we demonstrate that the fecundity effects of mutations and their dominance relationships need not be assumed, but can co-evolve in ways that depend on the selfing rate, the level of inbreeding depression expressed by selfed progeny, and the ecology of mating. First assuming that sex allocation is determined at a single locus, we show that the co-evolution of allelic effects and dominance readily leads to the gradual emergence of a heterogametic sex-determining locus. We demonstrate that whether selection favours XY or ZW sex determination depends on both the shape of gain curves and on the mating system in the hermaphroditic ancestor, with partial selfing and inbreeding depression promoting the evolution of XY over ZW systems. We then assume that individual sex allocation is initially determined by alleles at many loci scattered across the genome. We show that selection for dioecy favours the concentration of the genetic architecture of sex allocation, leading to the emergence of a single sex-determining locus with complete dominance, i.e., to either XY or ZW sex determination, as in the first model.

## Model and analyses

### Model

The ecological scenario we envisage follows closely from Charlesworth and Charlesworth (1978b, 1981). We consider a large population in which diploid individuals allocate a proportion *x* of their reproductive resources to their female and 1 − *x* to their male function, leading to a trade-off between both functions. Strategy *x* results in female and male fecundities 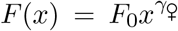 and 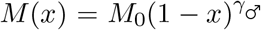, respectively, where *F*_0_ and *M*_0_ correspond to the maximum achievable fecundity, and exponents *γ*_♀_ and *γ*_♂_ control the shape of each gain curve and thus the nature of the trade-off between male and female functions (Figure 1A; many of our results are derived for functions *F* (*x*) and *M* (*x*) that are more general than these power functions; see Appendix). Following pollen and ovule production, individuals first self-fertilise a fraction *α*(*x*) of their ovules (‘prior selfing’; Lloyd, 1975), and then outcross the remaining 1 − *α*(*x*) via random mating. We assume that self-fertilisation (selfing hereafter) does not affect siring success through male function, but may decrease with allocation *x* to female function (i.e. *α*^*′*^(*x*) ⩽ 0; where necessary, we specifically assume that *α*(*x*) = *α*_0_(1 − *βx*), where 0 ⩽ *α*_0_ < 1 denotes the maximum achievable selfing rate and 0 ⩽ *β* ⩽ 1 controls the degree to which *α*(*x*) depends on female allocation, as in Charlesworth and Charlesworth, 1978b, 1981). Outcrossed offspring develop into viable seeds with probability 1, whereas selfed offspring develop into viable seeds with probability 1 − *δ*, where *δ* measures the magnitude of inbreeding depression. Finally, adults die and a new generation is formed from viable seeds (Figure 1B; see Appendix A for more details).

**Figure 1:**
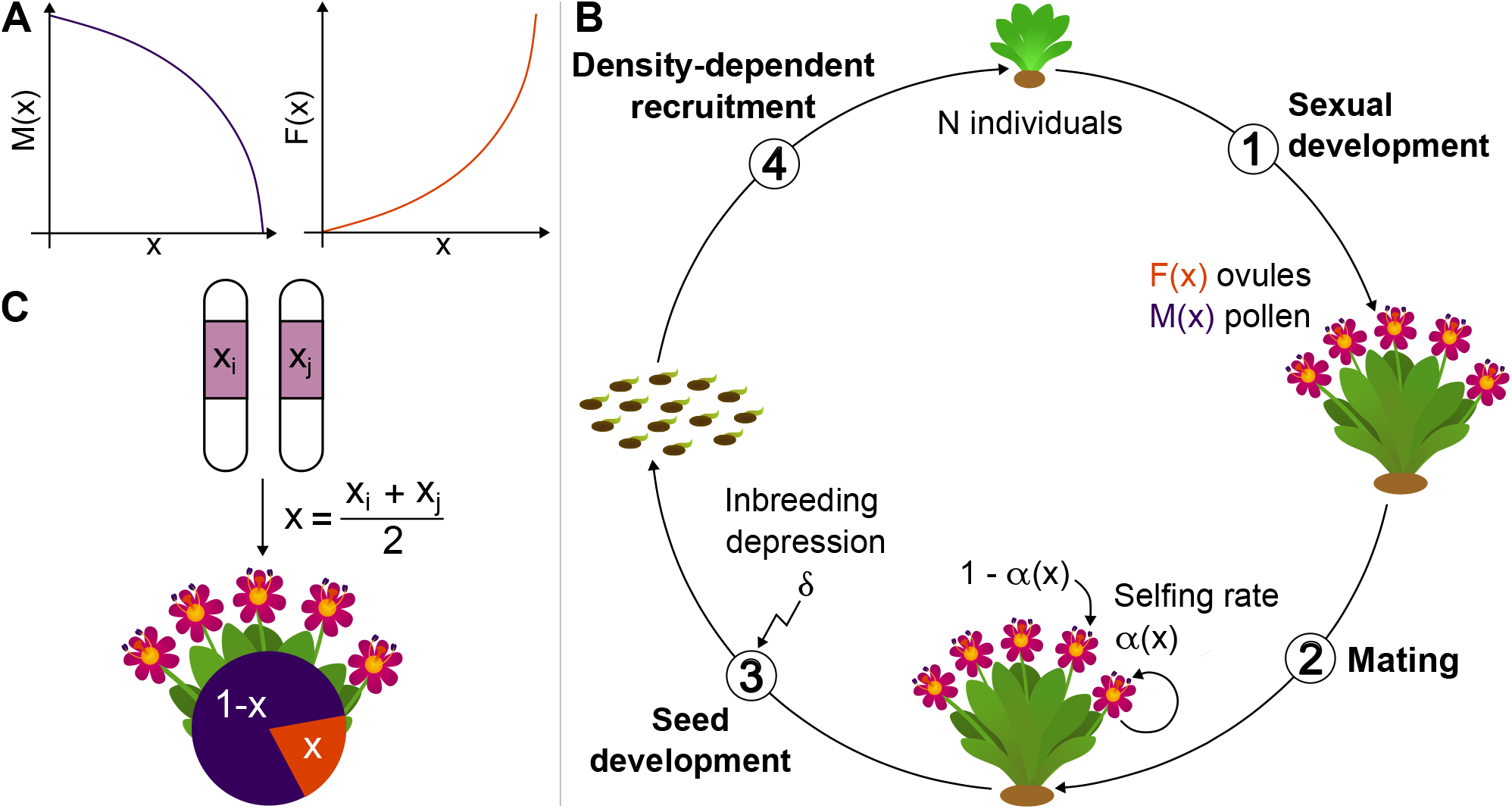
Life cycle and genetic architecture of sex allocation. **A** Male (*M* (*x*), dark purple) and female (*F* (*x*), orange) gain curves as functions of the fraction *x* of resources allocated to female function. In this example, the male gain curve is saturating, reflecting diminishing fitness returns through male function, whereas the female gain curve is accelerating, reflecting increasing fitness returns through female function. **B** Life cycle assumed in the model. See main text for details. **C** Genetic architecture of sex allocation in our baseline model. The sex allocation strategy *x* expressed by an individual is determined by its genotype at a quantitative trait locus where alleles are additive.

Previous theory demonstrates that dioecy is evolutionarily stable when gain curves are accelerating (*γ*_♀_ *>* 1 and *γ*_♂_ *>* 1, Charnov et al., 1976; Charnov, 1982) or when inbreeding depression is sufficiently strong (Charlesworth and Charlesworth, 1981); under these conditions, a hermaphrodite in a population of males and females will have lower than average fitness, and a population of hermaphrodites will be invadable by unisexuals. Existing theory further shows that dioecy may evolve from hermaphroditism through sequential invasions of fully dominant or fully recessive mutations causing complete (Charlesworth and Charlesworth, 1978a, 1981) or partial sterility (Charlesworth and Charlesworth, 1978b), most likely first of male and then female function.

Rather than fixing the nature and dominance of mutations a priori, we assume here that sex allocation *x* is influenced by a quantitative trait locus subject to recurrent mutations of small effects leading to gradual evolution (i.e., mutations create new alleles whose value deviates from the original allele by a small amount, the ‘continuum-of-alleles’ model; Fig. 1C; Kimura, 1965, p. 883 in Walsh and Lynch, 2018). This locus could be a regulatory element that influences the development of female and male traits, or one or more fully-linked genes that are independent targets of partial female and male sterility mutations (where in both cases there is a physiological trade-off among female and male functions; e.g., Charlesworth and Charlesworth, 1978a,b, 1981). Genetic effects on the phenotype *x* are initially assumed to be additive, meaning that the two alleles carried by an individual at the quantitative trait locus contribute equally to its expressed phenotype (note that although genetic effects on the phenotype are additive, they may translate to non-additive effects on fitness, as described by gain curves). To investigate the emergence of sex-determining systems, we later allow for the evolution of the genetic architecture of sex allocation *x* (i.e., we allow non-additive genetic effects on the phenotype *x* to evolve), first by considering the evolution of dominance at the quantitative trait locus, and then by extending our model to a case where sex allocation is influenced by multiple loci.

### Evolutionary analyses

We analyse our model using a combination of mathematical and simulation approaches. Our mathematical analyses assume that the population is large (so that drift can be ignored) and that mutations are rare and have small effects on the phenotype, such that evolutionary dynamics can be studied using the adaptive dynamics framework (Metz et al., 1996; Geritz et al., 1998; Dercole and Rinaldi, 2008; Otto and Day, 2011; Avila and Mullon, 2023). In this framework, the evolution of an initially homogeneous population proceeds in two phases. First, the population evolves under directional selection, whereby positively selected mutations sweep to fixation so that the population transits from being largely monomorphic for one trait value to being monomorphic for another. Through this process, the population may eventually attain a ‘convergence stable strategy’ *x*^*^, which is an attractor of evolutionary dynamics where directional selection vanishes (Eshel, 1983; Christiansen, 1991; Geritz et al., 1998). Once the population reaches this state, it experiences either stabilising selection (and therefore remains monomorphic) or negative frequency-dependent disruptive selection and becomes polymorphic in a process referred to as ‘evolutionary branching’ (Geritz et al., 1998; Pásztor et al., 2016). In this case, negative frequency-dependent selection maintains two alleles that initially encode weakly differentiated phenotypes around *x*^*^ (see Appendix 1 in Geritz et al., 1998 for a proof), that then become increasingly divergent as the two alleles accumulate further mutations. The distribution of allelic effects in the population thus gradually evolves from unimodal to bimodal as a result of negative frequency-dependent disruptive selection (‘disruptive selection’ hereafter) and small-effect mutations (Geritz and Kisdi, 2000).

In addition to our mathematical analyses, we run individual-based simulations to relax several of our assumptions, notably that the population is large and that mutations are rare. We also use these simulations to extend our model to a case where sex allocation is influenced by many loci. We provide more details on these later.

## Results

### Gradual evolution of sexual systems under complete outcrossing

We first assume that the population is fully outcrossing (*α*_0_ = 0) and focus on the effects of selection for sexual specialisation, as in classical sex allocation theory (Charnov et al., 1976; Charnov, 1982). We show in Appendix B.1 that the population either converges and remains monomorphic for the intermediate strategy *x*^*^ = *γ*_♀_*/*(*γ*_♀_ + *γ*_♂_), with all individuals being hermaphrodites, or experiences disruptive selection, resulting in the gradual differentiation of two types of alleles: one that causes its carrier to allocate more resources to female function, and the other more resources to male function. Which of these two outcomes occurs depends on the shape of gain curves, with disruptive selection requiring at least one of them to be sufficiently accelerating (specifically that 2*γ*_♀_*γ*_♂_ *> γ*_♀_ + *γ*_♂_; Fig. 2A). When both gain curves are accelerating (*γ*_♀_ *>* 1 and *γ*_♂_ *>* 1), disruptive selection leads to the co-existence of two alleles: one for a pure male (*x* = 0) and another for a pure female (*x* = 1) strategy. When only one curve is accelerating, one allele encodes a unisexual strategy (female or male), while the other encodes a hermaphroditic strategy, albeit biased towards the opposite sex (Appendix B.2 for analysis). Figs. 2B-E show how these different possible evolutionary dynamics unfold in individual-based simulations (detailed in Appendix B.3).

**Figure 2:**
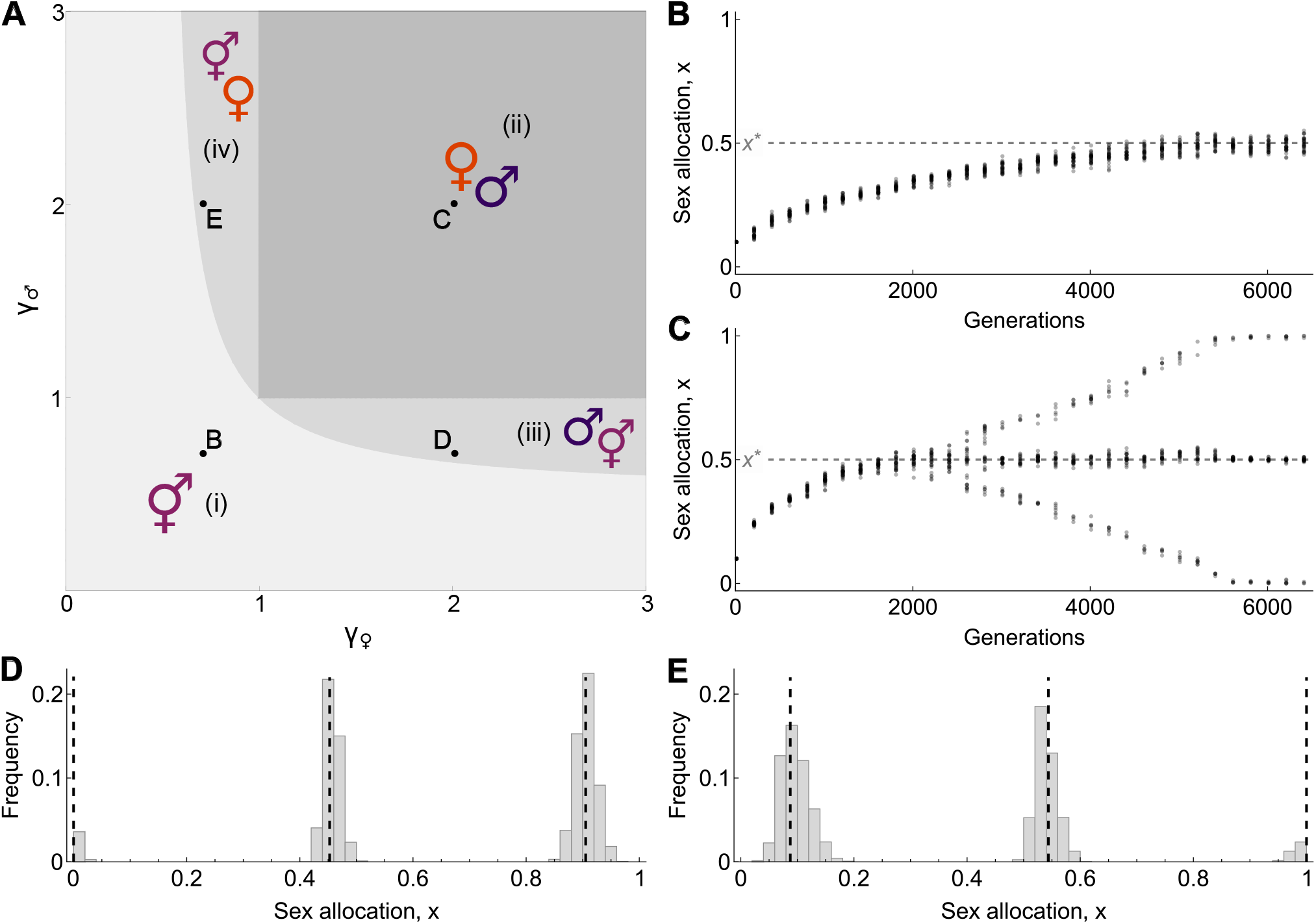
The gradual evolution of sex allocation and sexual systems under complete outcrossing. **A** The four outcomes of evolution according to *γ*_♀_ and *γ*_♂_ (Appendix B.1 for analysis): (i) hermaphroditism (light grey); (ii) dioecy (dark grey); (iii) androdioecy (medium light gray) and (iv) gynodioecy (medium light gray), where pure males and females coexist with hermaphrodites, respectively. **B-E** Results from individual based simulations showing the four possible outcomes outlined in Panel A. Simulations follow the evolution of a population of *N* = 10^4^ individuals, with a per-locus mutation rate of *µ* = 5 *×* 10^− 3^, and where a mutation creates a new allele whose genetic effect consists of its original value to which is added a small value randomly sampled from a Normal distribution with mean 0 and standard deviation *σ* = 10^− 2^ (Appendix B.3 for details on simulations). **B** The phenotypes expressed by 30 randomly sampled <u>i</u>ndividuals every 200 generations under conditions predicted to lead to hermaphroditism (with 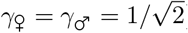). The population converges to express the equilibrium strategy *x*^*^ = *γ*_♀_*/*(*γ*_♀_ + *γ*_♂_) = 1*/*2, indicated by the light grey dashed line. **C** Same as B under conditions predicted to favour dioecy (with *γ*_♀_ = *γ*_♂_ = 2). Disruptive selection leads to the coexistence of pure male (*x* = 0) and female (*x* = 1) alleles. At equilibrium, the population is composed of males (with genotype {0,0}), females (with genotype {1,1}), and hermaphrodites (with genotype {1,0}). **D** Dis<u>t</u>ribution of phenotypes at equilibrium in a simulation where androdioecy evolves (with *γ*_♀_ = 2 and 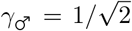). Dashed vertical lines indicate the equilibrium strategies the analytical model predicts, which are calculate<u>d</u> according to the method described in Appendix B.2. **E** Same as D where gynodioecy evolves (with 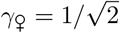 and *γ*_♂_ = 2).

The above results align with classical optimality models in that they delineate the same conditions for the evolutionary stability or instability of hermaphroditism (Charnov et al., 1976; Charnov, 1982, see Appendix B.4 for more details in this connection). Our analyses additionally demonstrate that an initially hermaphroditic population may evolve gradually towards the sexual systems predicted by classical optimality models (which are static models), via the gradual divergence of two initially similar alleles maintained as a polymorphism. This may appear at odds with Charlesworth and Charlesworth (1978b), who concluded that alleles with small effects on male and female fecundity rarely establish a genetic polymorphism (e.g. their inequality on p. 142 with *s* = 0 for complete outcrossing). Their conclusion, however, rests on the notion that the effect of a mutation on male fecundity is independent of its effect on female fecundity (statistically speaking). In contrast, the effects of a mutation on male and female functions are linked by a trade-off described by the gain curves in our model. We show that where these curves are such that selection is disruptive, the conditions derived by Charlesworth and Charlesworth (1978b) for the maintenance of polymorphism are satisfied (see Appendix B.5). From this point of view, our above results are thus entirely consistent with theirs.

### Emergence of XY and ZW sex determination through dominance evolution

Because we assumed so far that alleles have additive effects on sex allocation, disruptive selection in our model leads to the coexistence of not two but three types of individuals: two homozygotes that express female- and male-biased sex allocation strategies, respectively, and a heterozygote with an intermediate hermaphroditic strategy (Figure 2C-E), so that dioecy is incomplete. To examine how complete dioecy might ultimately evolve, we next model the co-evolution of sex allocation with dominance at the underlying locus. We first investigate this co-evolution using computer simulations, and then analyse a mathematical model to better understand the mechanisms governing it. In the simulations, we assume that the evolving locus is composed of two elements: a sex allocation gene, where alleles code for different sex allocation strategies; and a linked promoter that determines the level of expression of the sex allocation allele (Figure 3A). Variation at the promoter leads to variation in allelic expression through *cis* effects, which in turn determine the dominance relationships among sex allocation alleles (Van Dooren, 1999). We let the sex allocation gene and its promoter each undergo recurrent mutations of small effect (i.e. each follow the continuum-of-alleles model), so that dominance and sex allocation co-evolve (see Appendix C.1 for details on these simulations).

**Figure 3:**
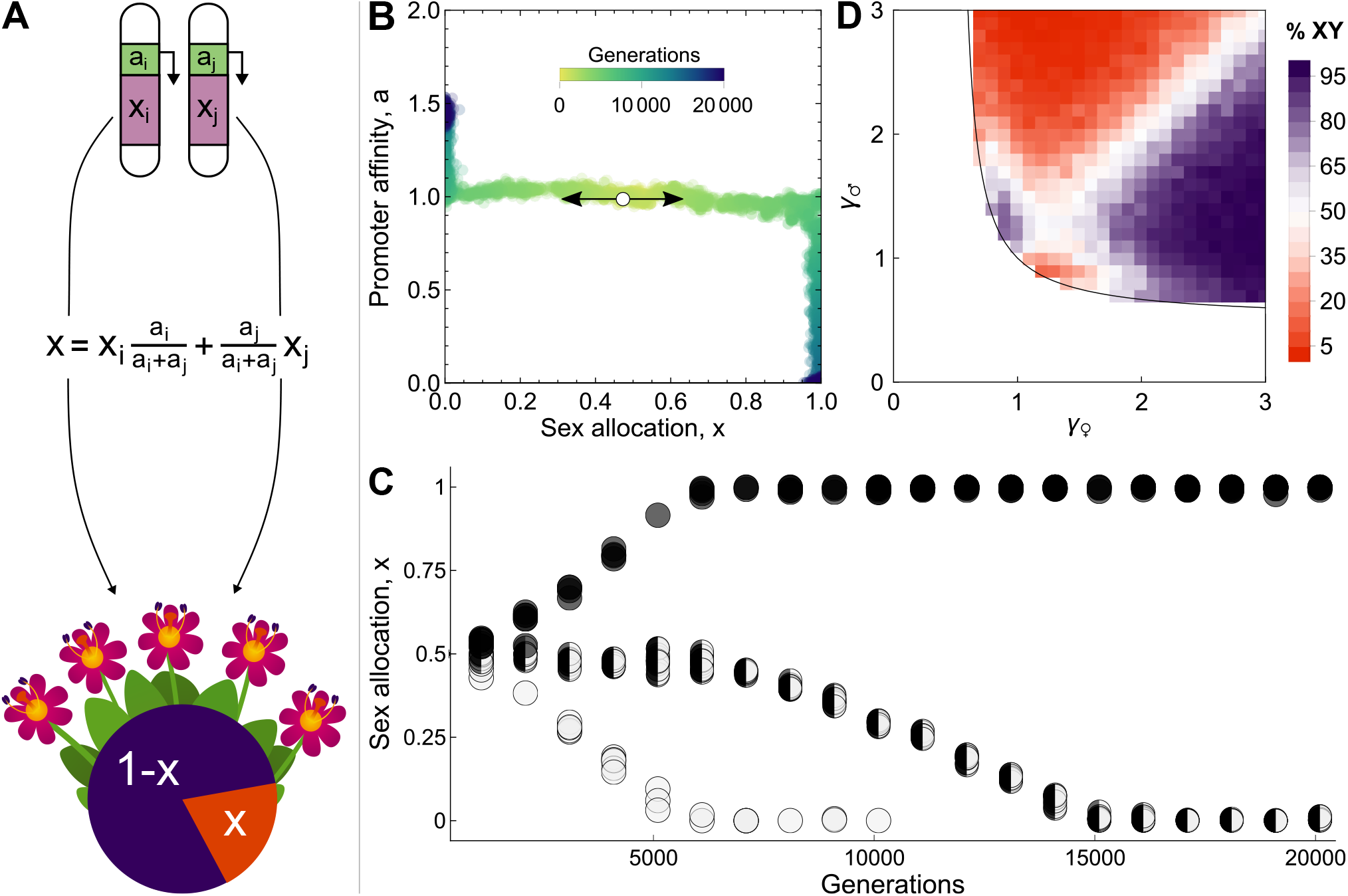
The co-evolution of sex allocation and gene-regulation. **A** Genetic architecture of sex allocation. The sex allocation locus is composed of a sex allocation gene and its promoter. Transcription factors must bind to the promoter for the sex allocation gene to be expressed, which they do at a rate that depends on the promoter’s affinity, *a*. Consequently, sex allocation alleles are expressed in proportion to their promoter’s affinity, and promoter affinities encode the dominance relationship between sex allocation alleles. In this example, alleles *x*_*i*_ and *x*_*j*_ are associated with promoters with affinities *a*_*i*_ and *a*_*j*_, so that they contribute in proportions *a*_*i*_*/*(*a*_*i*_ + *a*_*j*_) and *a*_*j*_*/*(*a*_*i*_ + *a*_*j*_) to the expressed sex allocation strategy *x*. **B** Phase diagram of sex allocation and promoter affinity when the two co-evolve in a simulation under conditions predicted to lead to dioecy (*γ*_♀_ = *γ*_♂_ = 2). Each dot depicts an allele, characterised by the sex allocation strategy it encodes and its promoter’s affinity. Colour indicates time since the start of the simulation (in generations), with darker colours indicating later times. The population is initially monomorphic with *x*_0_ = 0.5 and *a*_0_ = 1 (white circle). Here, the male allele becomes associated with an increasingly high affinity promoter while the female allele becomes associated with an increasingly low affinity one, leading to complete dominance of the male allele and the emergence of XY sex determination. (Parameters: *N* = 10^4^, Appendix C.1 for simulation details). **C** Phenotypes expressed by individuals as a function of time for the same simulation as figure B. Each circle depicts an individual. Fully black and white circles depict homozygotes for female- and male-biased alleles, respectively, whereas half black and white circles depict heterozygotes (defined as individuals bearing two alleles that are more different than the average difference between two alleles within the same individual). As sex allocation alleles diverge and dominance evolves, heterozygotes gradually become more male-biased, and eventually replace male homozygotes, thereby achieving dioecy with XY sex determination. **D** Proportion of XY systems evolving out of 200 simulations with *N* = 10^3^, for values of *γ*_♀_ and *γ*_♂_ spanning the parameter range in which selection on sex allocation is disruptive. XY and ZW systems are equally likely to emerge when *γ*_♀_ = *γ*_♂_, whereas XY systems are more prevalent where *γ*_♀_ *> γ*_♂_ and ZW systems where *γ*_♀_ *< γ*_♂_ when gain curves are sufficiently accelerating. When gain curves are close to being linear, the correspondence between gain curve shape and the proportion of XY evolving is reversed. Parameters used in all simulations: mutation probability *µ* = 5 *×* 10^− 3^, standard deviation in mutational effect *σ* = 10^− 2^.

We first run simulations under conditions predicted to lead to pure male and female alleles (so when *γ*_♀_ *>* 1 and *γ*_♂_ *>* 1). In these simulations, complete dominance of one sex allocation allele always evolves, so the population ultimately comprises only males and females, such that dioecy is complete (Figure 3B-C). Remarkably, whether the male or the female allele becomes dominant depends strongly on male and female gain curves (i.e., on *γ*_♀_ and *γ*_♂_, Figure 3D). Provided that neither curve is close to being linear, the male allele is more likely to become dominant when fitness increases more steeply via female function (i.e., when *γ*_♀_ *> γ*_♂_), leading to the emergence of an XY system. Conversely, when fitness returns increase more steeply via male function (i.e., when *γ*_♂_ *> γ*_♀_), the female allele most often becomes dominant, leading to a ZW system. When both gain curves are close to being linear (i.e., when *γ*_♂_ and *γ*_♀_ are close to one), this correspondence is reversed so that XY is favoured when *γ*_♂_ *> γ*_♀_, and ZW when *γ*_♀_ *> γ*_♂_. We also simulated scenarios predicted to lead to gyno- and androdioecy, where pure females and pure males coexist with hermaphrodites, respectively (i.e., with either *γ*_♀_ *>* 1 or *γ*_♂_ *>* 1), and obtained qualitatively similar results: the allele for the unisexual strategy most often becomes dominant (Figure 3D), so that the population typically ends up being composed of either heterozygote (XY) males and homozygote (XX) hermaphrodites (when *γ*_♀_ *>* 1), or heterozygote (ZW) females and homozygote (ZZ) hermaphrodites (when *γ*_♂_ *>* 1).

### Competition through male and female functions determines whether XY or ZW evolves

To better understand the nature of selection on dominance, we analyse mathematically a version of our model in which dominance is treated as a quantitative trait. In this version, two sex allocation alleles are maintained as a polymorphism by disruptive selection, *x*_♀_ and *x*_♂_, where one allele encodes a more female strategy than the other (*x*_♀_ *> x*_♂_, hereafter referred to as ‘female’ and ‘male’ alleles). In *x*_♂_*/x*_♀_ heterozygotes, the female allele is expressed proportionally to a dominance coefficient *h*, so that the sex allocation strategy of a *x*_♂_*/x*_♀_ heterozygote is given by *h x*_♀_ +(1 − *h*) *x*_♂_. We assume that the value of *h* is determined by a quantitative trait locus subject to recurrent mutations of small effect and unlinked to the sex allocation locus. This allows us to investigate the nature of selection on other mechanisms that may modify dominance (e.g. *trans* effects, Billiard et al., 2021; see Appendix C.2 for details on this model and its analysis).

The selection gradient on *h*, which gives the direction and strength of selection acting on mutations modifying dominance in a population expressing *h*, reveals that there exists a threshold *h*^*^ below which selection favours ever lower values of *h* (i.e. *h* → 0 when *h < h*^*^) and above which selection favours ever higher values of *h* (i.e., *h* → 1 when *h > h*\*). Complete dominance of either the male or female allele therefore also always evolves here, resulting in the emergence of XY or ZW sex determination, respectively (Fig. 4A). Computing *h*^*^ explicitly is difficult, but its position relative to 1*/*2 can be inferred from the sign of the selection gradient at *h* = 1*/*2, with a positive gradient indicating that *h*^*^ *>* 1*/*2 (such that XY is favoured), and a negative gradient indicating that *h*^*^ < 1*/*2 (such that ZW is favoured). In fact, we observe an almost perfect correspondence between this analysis and the outcome of our earlier individual-based simulations (compare Fig. 3D with Fig. 4B). This shows that whether selection promotes XY or ZW sex determination is independent of the particular mechanisms responsible for variation in dominance (whether through *cis* or *trans* effects), but rather comes down to the shape of the gain curves in the absence of selfing.

**Figure 4:**
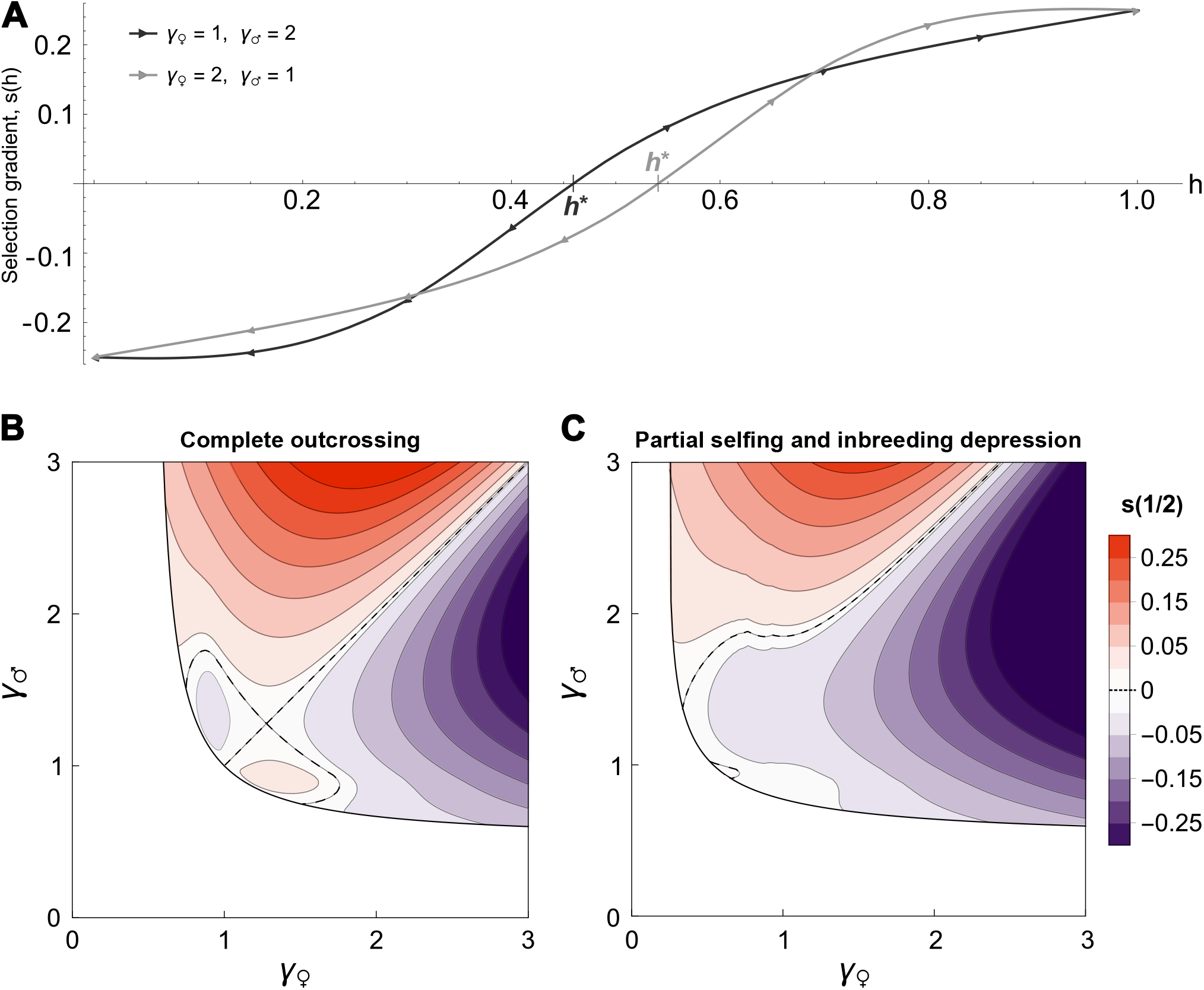
The nature of selection on dominance at a sex-determining locus. **A** Selection gradient *s*(*h*) acting on the dominance *h* of the female allele *x*_♀_ = 1 over the male allele *x*_♂_ = 0 for two cases leading to dioecy (Appendix C.2.4 for how to compute this gradient). The selection gradient *s*(*h*) is negative when *h* is smaller than the threshold *h*^*^ such that *s*(*h*^*^) = 0, and positive when *h* is greater than *h*^*^. Therefore, selection always eventually leads to either *h* = 0 (leading to an XY system) or *h* = 1 (leading to a ZW system). Additionally, the larger *h*^*^ is, the more readily an XY system should evolve and conversely, the smaller *h*^*^, the more likely a ZW system evolves. For the examples shown here, we expect to see an XY in the case depicted in gray and a ZW in black. **B** Selection gradient on dominance at additivity, *s*(1*/*2), in the complete outcrossing case (*α*_0_ = 0). Dashed lines indicate points where the gradient is zero so that XY and ZW sex determination are equally likely to evolve. Orange shades are for *s*(1*/*2) *>* 0, which indicates that *h*^*^ < 1*/*2 and thus that ZW sex determination is favoured, whereas purple shades are for *s*(1*/*2) < 0, which entails that *h*^*^ *>* 1*/*2 and XY sex determination is favoured. Variations in the sign and intensity of the selection gradient match almost perfectly with the proportion of XY systems evolving in simulations (Fig. 3D). **C** Same as B but with partial selfing and strong inbreeding depression (*α*_0_ = 0.75, *δ* = 0.75, *β* = 1). The parameter space in which an XY system is favoured becomes much larger than the one in which a ZW system is favoured, indicating that selfing promotes XY over ZW sex determination.

Decomposing the selection gradient on dominance reveals that selection on sex-determining systems and its relationship with gain curves can be understood as follows (Appendix C.2.4 for details). Selection on dominance *h* acts only in *x*_♂_*/x*_♀_ heterozygotes, which are hermaphrodites. Such a heterozygote can become more female (or more male) through an increase (or a decrease) in *h*. However, whatever the change in dominance, this heterozygote will always be less fit than female homozygotes through female function, and less fit than male homozygotes through male function. In fact, homozygotes are typically so competitive because of their fecundity advantage, that it is best for a heterozygote to allocate more to the sex in which this advantage is weakest. This scenario favours heterozygote individuals that are more female when *γ*_♂_ *> γ*_♀_ and more male when *γ*_♂_ *< γ*_♀_, leading to the evolution of ZW and XY systems, respectively. When both gain curves are close to linear (*γ*_♀_ and *γ*_♂_ close to one), the advantage of homozygotes over herterozygotes is reduced, and it is then best for a heterozygote to allocate to the sexual function that leads to the greater increase in fecundity, i.e., to become more female when *γ*_♀_ *> γ*_♂_ and more male when *γ*_♀_ *< γ*_♂_.

### Partial selfing and inbreeding depression favour XY sex determination

Our analysis so far has assumed that hermaphrodites are completely outcrossing. However, it is well-established that partial selfing and inbreeding depression can play an important role in the evolution of dioecy and other polymorphic sexual systems such as gyno- and androdioecy (Charlesworth and Charlesworth, 1978a,b, 1981). To examine how these factors influence the gradual evolution of sexual systems and sex determination, we now analyse our model for *α*_0_ > 0 and *β* > 0 (see Appendix D for details).

To investigate the influence of selfing on the gradual emergence of polymorphism, we first fix dominance at the sex allocation locus (see Appendices D.1-D.2). Previous analyses have found that the invasion of a partial male-sterility mutation in a population of hermaphrodites is either facilitated or hindered by partial selfing, depending on whether inbreeding depression is high or low, respectively (Charlesworth and Charlesworth, 1978b). Consistent with these observations, we find that selfing favours disruptive selection, and thus the emergence of polymorphism in sex allocation, when inbreeding depression is high (*δ >* 1*/*2), whereas it inhibits polymorphism when inbreeding depression is low (*δ <* 1*/*2, see Fig. S12 in the Appendix). By decomposing the disruptive selection coefficient (eq. D17 in Appendix D), we further reveal that this effect of selfing stems from the interplay between its twofold transmission advantage and the deleterious effects of inbreeding depression (Fisher, 1941), which influences fitness gained through female function. In particular, when *δ >* 1*/*2 (i.e., when a selfed individual is less than half as fit as an outcrossed individual), an individual transmits on average more copies of its genes to the next generation by outcrossing than by self-fertilising its seeds. In this case, increased allocation into female function leads to multiplicative fitness benefits, as it allows individuals to produce not only more seeds but also seeds that transmit on average more copies of their genes due to increased outcrossing (since *β >* 0). Such multiplicative benefits favour polymorphism, and allow the emergence of dioecy even when both gain curves are saturating (i.e. where *γ*_♀_ < 1 and *γ*_♂_ < 1; Fig. 4C).

These analyses reinforce the conceptual link between the evolution of dioecy in response to selection for inbreeding avoidance, as exposed by population genetic models (Charlesworth and Charlesworth, 1978a,b), and its evolution in response to selection for sexual specialisation, as modelled in terms of the shape of gain curves in sex allocation theory (Charnov et al., 1976; Charnov, 1982). While we have used the term ‘gain curves’ to refer to the relationship between resource allocation and fecundity in our model (i.e., the number of gametes produced, similar to previous authors; e.g., Charlesworth and Charlesworth, 1981), gain curves can more generally be thought of as functions relating resource allocation in one sex to *fitness* gained through that sex (i.e., the total number of gene copies transmitted through seeds or pollen, Charnov, 1982). The shape of fecundity and fitness gain curves are the same in our baseline model under complete outcrossing, but they may differ depending on factors influencing competition.

In particular, when the selfing rate is coupled with sex allocation, partial selfing alters the shape of the female fitness gain curve, causing it to accelerate when inbreeding depression is greater than 1*/*2 (see also Appendix F for an example in which the coupling of sex allocation with seed dispersal modulates the shape of the female fitness gain curve).

To study the effect of partial selfing on the evolution of sex determination, we next investigate selection on dominance when disruptive selection favours polymorphic sexual systems (as in section “Competition through male and female functions determines whether XY or ZW evolves”; Appendix D.3 for details on these analyses). We show that partial selfing favours the evolution of XY over ZW sex determination, especially when inbreeding depression is high (Fig. 4C). This is because selfing increases competition for reproduction through female relative to male function, and inbreeding depression reduces the reproductive value of offspring produced via the female function (i.e., it reduces the relative influence of self-fertilised offspring on the long-term demography of the population; Charlesworth, 1980; Caswell, 2001; Rousset, 2004). The combination of these two effects means that, in a population where male and female alleles segregate, an intermediate, hermaphroditic heterozygote is better off allocating more resources to its male function, as this reduces the competition from homozygotes and boosts the reproductive value of its offspring. Together, these conditions favours the evolution of dominance of the male over the female allele, and therefore the emergence of XY sex determination.

### Disruptive selection promotes the concentration of the genetic basis of sex and leads to smooth transitions among sexual systems

For practical reasons, we have assumed thus far that sex allocation is the outcome of allelic expression at a single locus. However, sex allocation in hermaphroditic populations may often be a quantitative trait influenced by many loci (Meagher, 1999; Ashman, 2003; Mazer et al., 2007). This possibility raises the question of how dioecy might evolve in a hermaphroditic population in which variation in sex allocation has a polygenic basis. Previous modelling has shown that disruptive selection promotes the concentration of the genetic basis of traits from many to few or only one locus, as this results in greater heritability of the differentiated phenotypes (van Doorn and Dieckmann, 2006; Kopp and Hermisson, 2006). To study how this might occur in the evolution of dioecy, we extend our simulations to a scenario where sex allocation is initially determined by *L* freely-recombining sex allocation loci. In addition, we introduce a modifier locus at which alleles that determine the contribution of each locus to the phenotype can segregate (for instance, one allele may code for an equal contribution of each of the *L* sex allocation loci, while another may cause one of the *L* loci to determine most of the variation in sex allocation, Kopp and Hermisson, 2006; Appendix E for details). Alleles at the modifier locus are subject to small-effect mutations, so that the relative contribution of loci to sex allocation evolves jointly with allelic effects and dominance at the loci concerned, all in a gradual manner.

To see the effects of disruptive selection on the genetic basis of sex, we assume that each of the *L* sex allocation loci initially contributes equally to the trait and that conditions are such that selection initially favours hermaphroditism (e.g., we assume that the gain curves saturate). Simulations show that, in this case, the contributions of the different loci to the phenotype, though variable, remain similar (Fig. 5A, shaded area). The population, meanwhile, shows a unimodal trait distribution centred around the optimal value *x*^*^ (Fig. 5B, shaded area). Suppose then that, at some given generation, conditions change such that selection on sex allocation now favours dioecy (e.g., gain curves now accelerate). When this occurs, we observe the progressive silencing of all but one locus, whose relative contribution to the trait keeps increasing until it explains all variation in sex allocation (Fig. 5A, non-shaded area). This concentration of sex allocation to a single locus allows for the concomitant evolution of separate sexes in the population (Fig. 5B, non-shaded area). In contrast to our earlier one-locus simulations, the progressive concentration of the genetic basis of sex means that, here, the transition from hermaphroditism to dioecy occurs via the gradual divergence of essentially two observable phenotypes: males and females (Fig. 5B, instead of passing through a trimorphic state with three sex-allocation phenotypes, as shown in Fig. 3C).

**Figure 5:**
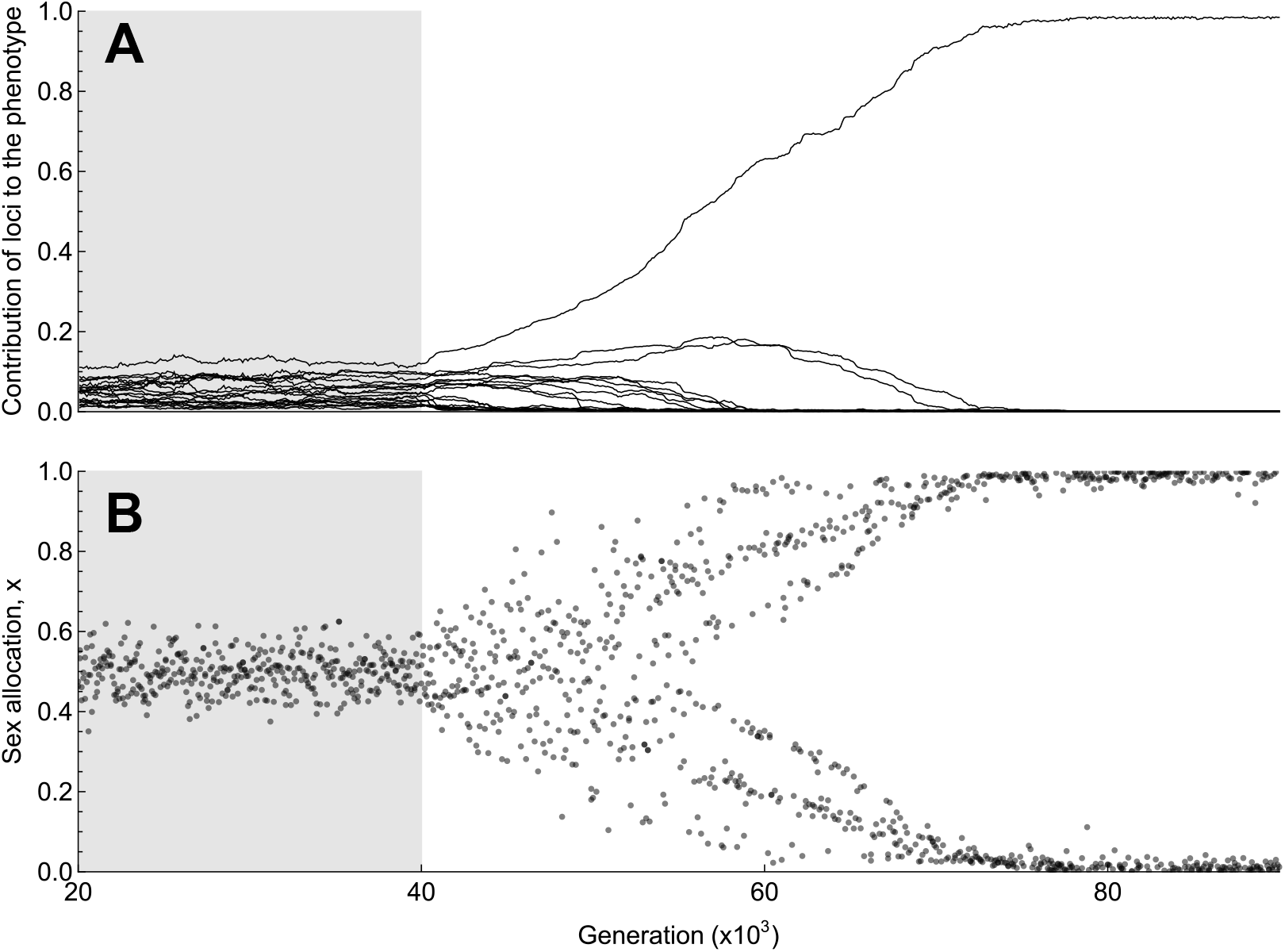
Concentration of the genetic architecture of sex allocation in response to selection for dioecy. Results of a simulation with *L* = 20 loci initially contributing equally to the sex allocation strategy expressed by individuals (Appendix E for details on these simulations). Selection favours hermaphroditism for the first 40,000 generations (i.e. gain curves are saturating, with *γ*_♀_ = *γ*_♂_ = 1*/*2; grey background in the plots). An ecological change then occurs, causing selection to favour dioecy for the rest of the simulations (i.e. gain curves become accelerating, with *γ*_♀_ = *γ*_♂_ = 2; white background in the plot). **A** Relative contributions of the 20 quantitative trait loci to the phenotype as a function of time (after a burn-in period of 20,000 generations).When hermaphroditism is favoured (before the ecological change), loci contributions vary due to drift but remain roughly equal (on average 0.05). In contrast, when dioecy is favoured, selection drives the evolutionary dynamics of loci contributions, leading all but one locus to become silenced (i.e. to not contribute to the sex allocation phenotype), with the remaining locus acting as the sex-determining locus. **B** Sex allocation strategies expressed in the population as a function of time. While selection favours hermaphroditism, the population remains unimodally distributed around *x*^*^ = *γ*_♀_*/*(*γ*_♀_ + *γ*_♂_) with little phenotypic variance. When dioecy becomes favoured, the phenotypic variance increases and the distribution gradually shifts from unimodal to bimodal, ultimately achieving dioecy. Parameters used in simulations: population size *N* = 5 *×* 10^3^; mutation probability *µ* = 10^− 2^; standard deviation in mutational effect *σ* = 5 *×* 10^−2^.

## Discussion

Dioecy is thought to have often evolved from hermaphroditism in a step-wise process that involves gynodioecy as an intermediate step, with XY or ZW sex determination emerging if the initial mutation causing male sterility was fully recessive or fully dominant, respectively (the ‘gynodioecy’ pathway, Charlesworth and Charlesworth, 1978a, 1981). An alternative scenario for transitions to dioecy invokes the gradual divergence in sex allocation of hermaphrodites in response to selection for sexual specialisation (the ‘monoecy-paradioecy’ pathway, Lloyd, 1980). This scenario has been much discussed (Lloyd, 1980; Renner and Ricklefs, 1995; Cronk, 2022; Pannell and Jordan, 2022), but there has so far been little theoretical investigation of how it might unfold and lead to XY or ZW sex determination (Charlesworth and Charlesworth, 1978b). Here, we conducted a formal analysis of the gradual evolution of dioecy from hermaphroditism, and showed that heterogametic sex determination can be the outcome of a gradual adaptive process involving the joint evolution of sex allocation with its genetic architecture.

Our results demonstrate that selection can act on dominance at a sex-determining locus, thereby providing an adaptive hypothesis for why some species transitioning to dioecy acquire XY while others acquire ZW sex determination. Namely, we found that selection can influence whether XY or ZW sex determination evolves, and that which of these two systems is more likely to emerge depends on the mating system of ancestral hermaphrodites as well as on the trade-off between male and female allocation and its fitness implications (as described by fitness gain curves). Under complete outcrossing, the conditions favouring XY or ZW sex determination are symmetrical, with selection favouring dominance of the allele for the sex where the benefits of sexual specialisation are the weakest (Fig. 4B). However, this symmetry is broken when dioecy evolves in populations of partially self-fertilising hermaphrodites, in which case the emergence of XY sex determination is more likely, especially when inbreeding depression is high and selfing is frequent (Fig. 4C). Given that most dioecious plants documented so far have XY systems (about 85%, Ming et al., 2011), our results yield a new argument in support of the suggestion that dioecy might often evolve from hermaphroditism as a device to avoid inbreeding (Charlesworth and Charlesworth, 1978a,b).

In addition to evolving in response to selection for inbreeding avoidance, dioecy may also evolve when the ecological context favours sexual specialisation (Charnov et al., 1976; Freeman et al., 1997). In this case, our model predicts that whether an XY or a ZW system evolves should depend on how ecology influences the relative shapes of the male and female fitness gain curves, with XY favoured over ZW when the female gain curve is more accelerating than the male one. To the extent that dioecy might have evolved in response to selection for sexual specialisation, our results thus suggest that the observed excess of XY systems in dioecious plants could also be contributed to by cases in which the female gain curve is more accelerating than the male one. There are still very few empirical estimates of the shape of these curves, but it is generally thought that they are more likely to be saturating than accelerating, e.g., due to local mate and resource competition under limited pollen or seed dispersal, respectively (Hamilton, 1967; Taylor and Bulmer, 1980; Charnov, 1982; Brunet, 1992; Charlesworth, 1999; Pannell and Jordan, 2022). Indeed, the high prevalence of hermaphroditism in flowering plants may well be attributable to largely saturating gain curves (Käfer et al., 2017). Nevertheless, ecological mechanisms that may cause gain curves to accelerate have been proposed in the literature (Bawa, 1980; Givnish, 1982; Freeman et al., 1997; Charlesworth, 1999; Pannell and Jordan, 2022). In plants with fleshy fruits, for instance, individuals producing larger crops of fruits (i.e., allocating more heavily to female function) may achieve more efficient seed dispersal due to increased attractiveness to animal dispersers. This coupling of sex allocation with seed dispersal can generate multiplicative benefits to specialising into female function through reduced kin competition among seeds of more female individuals, thereby causing the female gain curve to accelerate (Givnish, 1982; Vamosi et al., 2007; Biernaskie, 2010, see also Appendix F for a mathematical formalisation of this argument). Similarly, seeds of plants producing larger seed crops may benefit from a lower predation risk due to ‘predator satiation’ (Janzen, 1971; Lloyd, 1982), which could also lead the female gain curve to accelerate through a coupling between seed survival and seed production. Furthermore, we showed in this paper that the coupling of the selfing rate with sex allocation can be seen as a mechanism causing the female gain curve to accelerate under conditions of high inbreeding depression, highlighting the connection between selection for inbreeding avoidance and the shape of gain curves. The above arguments point to the possibility of an accelerating female gain curve. In contrast, hardly any argument has been developed for why the male gain curve might accelerate for ecological reasons (Lloyd, 1982; Zhang, 2006). The possibility that female gain curves might accelerate more often than male gain curves thus deserves greater attention. Certainly, however, there is a great need for further empirical estimates of the shape of both male and female fitness gain curves.

Irrespective of why selection promotes dioecy, our model also throws light on the emergence of single-locus sex determination from an initially polygenic basis of sex allocation in hermaphrodites. Specifically, our multilocus simulations reveal that selection for dioecy favours the concentration of genetic variation in sex allocation at a single (sex-determining) locus. Empirically, the way and the speed at which this concentration materialises will depend on the genomic processes involved, which may include, for example, rearrangements of the regulatory network, recombination suppression, or gene duplication (Bachtrog et al., 2014; Henry et al., 2018), and on the amount of available standing variation. There is ample evidence for polygenic variation for sex allocation in many hermaphroditic taxa (as recently shown in e.g. *Mercurialis annua*, Cossard et al., 2021, or *Schiedea salicaria*, Campbell et al., 2022; for reviews see Meagher, 1999, Table 1 in Ashman, 2003 and in Mazer et al., 2007), but the specific loci involved are not yet known for any species. In dioecious plants, meanwhile, the specific genes involved in sex determination have only been described in a handful of species, with sex-determining loci consisting of either one master switch (e.g., in persimmon, poplar and willow, Akagi et al., 2014; Müller et al., 2020) or two fully-linked genes at which sterility mutations segregate as expected under the ‘two-gene model’ (e.g., in asparagus and kiwifruit, Akagi et al., 2019; Harkess et al., 2020; see also Westergaard, 1958 for phenotypic evidence consistent with this type of architecture). Either of these genetic architectures is compatible with the outcome of our model, which sees dioecy ultimately achieved through a single dominant Mendelian element. Together, our results suggest that valuable insights could be gained from studying the role played in the genetic control of sex allocation in hermaphrodites by genes involved in sex determination in closely related dioecious taxa.

The gradual scenario we describe might be especially relevant to some of the frequent transitions from monoecy to dioecy that appear to have taken place in flowering plants, with individuals gradually diverging in the number of their male and female flowers (Renner and Ricklefs, 1995; Cronk, 2022). In fact, taxa comprising closely related dioecious and monoecious species might be particularly well-suited to investigate the evolutionary dynamics outlined in our model, as sex allocation is more easily quantified in monoecious plants (where male and female flowers can be counted) than in species with bisexual flowers. In animals, our model might also be useful to understand the evolution of separate sexes from hermaphroditism in taxa such as polychaete annelids (e.g., in the genus *Ophryotrocha*; Picchi and Lorenzi, 2018) and flatworms (e.g., in the genus *Schistosoma*; Ramm, 2016), or the evolution of ‘split sex-ratios’ in ants and other social Hymenoptera, where colonies produce either male or female sexuals leading to a form of colony-level dioecy (Meunier et al., 2008; Kuemmerli and Keller, 2009; Lagunas-Robles et al., 2021).

Theoretical investigations of the economics of sex allocation and the evolution of sex determination have so far relied largely on two different modelling approaches - optimality models on the one hand and population genetics on the other hand. In this paper, we have established a link between these two types of models. In doing so, we have restated several key predictions made by each of them, including general conditions for the maintenance of dioecy, gynodioecy, androdioecy and dioecy (Charnov et al., 1976; Charnov, 1982), as well as the importance of inbreeding avoidance as an evolutionary force favouring dioecy (Charlesworth and Charlesworth, 1978a,b, 1981). Building on these foundations, we have thrown new light onto how selection might shape the genetic architecture of sex determination, and, especially, how this architecture could depend on the ecology of mating.

## Acknowledgements

The authors thank Michel Chapuisat for an interesting discussion on split sex ratios, Aline Muyle for suggesting useful references and Gabriel Marais for comments on a previous version of the manuscript. The authors thank the Swiss National Science Foundation (SNF grants 310030 185196 to JRP and PCEFP3181243 to CM) for funding.

## Authors contributions

TL, JRP and CM conceptualised the study. TL performed the analysis under the supervision of CM. TL wrote the initial draft, and all three authors contributed to the final version. JRP and CM acquired the funding for the project.

## Appendix A

### Model

Our model considers a large population of diploid hermaphrodites with the following life-cycle. (i) *Sexual development:* First, individuals allocate resources to female and male functions in proportions *x* and 1 − *x*, respectively. An individual that invests *x* into female function produces *F* (*x*) ovules and *M* (*x*) pollen grains in large numbers (so we can ignore demographic stochasticity). We use the term ‘gametes’ to generically refer to ovules and pollen hereafter for simplicity, although these are haploid *gametophytes* rather than gametes. (ii) *Mating:* Following gamete production, individuals self-fertilise a fraction *α*(*x*) of their ovules prior to any opportunity for outcrossing (i.e., we assume *prior* selfing; Lloyd, 1975). The remaining fraction 1 − *α*(*x*) is outcrossed via random mating. Self-fertilisation (selfing hereafter) is assumed to require a negligible amount of pollen, so that the selfing rate *α*(*x*) does not affect siring success through male function. Following Charlesworth and Charlesworth (1978b, 1981), we assume that the selfing rate *α*(*x*) is a decreasing function of allocation *x* to female function and that fully female individuals (i.e. that express *x* = 1) cannot self-fertilise, specifically we assume

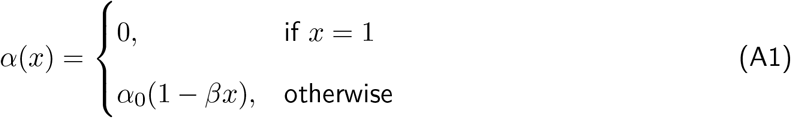

where *α*_0_ is the maximum achievable selfing rate and 0 ⩽ *β* ⩽ 1 controls the degree to which *α*(*x*) depends on female allocation. (iii) *Seed development:* Outcrossed zygotes develop into viable seeds with probability 1, whereas selfed zygotes suffer from inbreeding depression, so that they develop into viable seeds with probability 1−*δ*, where *δ* measures the magnitude of inbreeding depression (Charlesworth and Charlesworth, 1987). (iv) *Density regulation:* All adults die and are replaced by *N* uniformly sampled zygotes. The population size *N* is assumed large so that genetic drift can be ignored (we later relax this assumption).

We are interested in the evolution of the sex allocation strategy 0 ⩽ *x* ⩽ 1, and in particular whether such evolution can lead to polymorphic sexual systems. We assume that an individual investing no resources in either female (*x* = 0) or male (*x* = 1) function produces no gametes of that sex, i.e.

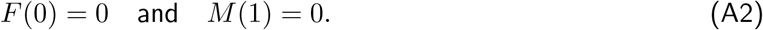

A dioecious population would thus be characterised by the coexistence of males, which express *x* = 0, and females, which express *x* = 1. For consistency, any increased investment into a sex function results in an increased number of gametes of that sex:

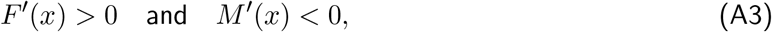

where the prime ^*′*^ denotes differentiation. Throughout the appendix, we consider general functions *F* (*x*) and *M* (*x*), also referred to as gain curves, but where relevant we use the more specific power functions,

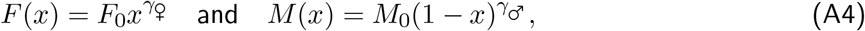

which form the basis of our results presented in the main text. In these functions, *F*_0_ and *M*_0_ are the maximum female and male fecundities, with *N* ≪ *F*_0_ ≪ *M*_0_ (so that there is no pollen limitation and no demographic stochasticity). The parameters *γ*_♀_ > 0 and *γ*_♂_ > 0, meanwhile, control the shape of the gain curves (Fig. S1). When *γ*_*u*_ = 1, extra investment in sex *u* ∈ {_♀_, _♂_} results in a linear increase in fecundity through that sex. By contrast, when *γ*_*u*_ > 1 that increase is accelerating, and saturating when *γ*_*u*_ < 1.

## Appendix B Gradual evolution of sexual systems under complete outcrossing

In this appendix, we study the evolution of sex allocation under complete outcrossing (so when *α*_0_ = 0 in eq. A1). We derive the results presented in the main text section “Gradual evolution of sexual systems under complete outcrossing” and Fig. 2.

### B.1 Evolutionary dynamics of sex allocation

To model the evolution of the sex allocation strategy *x*, we assume that this trait is genetically encoded by alleles with additive effects at a quantitative trait locus. We label alleles at this locus by their quantitative phenotypic effects, e.g., a carrier of alleles *x*_1_ ∈ [0, 1] and *x*_2_ ∈ [0, 1] expresses a sex allocation strategy *x* = (*x*_1_ + *x*_2_)*/*2. Evolution occurs through selection on mutations that arise at a constant rate, which is assumed to be small, and that have weak, unbiased phenotypic effects. In this case, evolutionary dynamics can be inferred from an invasion analysis that is detailed below.

### B.1.1 Invasion fitness

We first characterise the invasion fitness *W* (*x*_mut_, *x*) of a rare genetic mutation *x*_mut_, arising as a single copy in a resident population monomorphic for *x* (i.e., the geometric growth rate of the mutation when it is rare). Since mating is random, the *x*_mut_ allele can only be found in heterozygous form when rare. As a result, *W* (*x*_mut_, *x*) is given by the expected number of *x*_mut_*/x* heterozygotes produced by a *x*_mut_*/x* heterozygote over one full iteration of the life-cycle (Geritz and Kisdi, 2000; Metz and Leimar, 2011).

To express *W* (*x*_mut_, *x*), let *ω*_♀_(*x*_mut_, *x*) and *ω*_♂_(*x*_mut_, *x*) be the number of *x*_mut_*/x* heterozygotes produced by a *x*_mut_*/x* heterozygote through female and male gametes, respectively, before density-regulation (before step (iv) of the life-cycle in Appendix A). With population size *N* large enough for self-fertilisation to be negligible under random mating, these can be expressed as

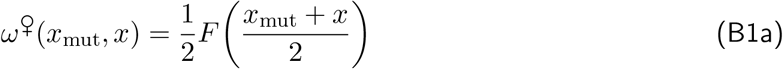

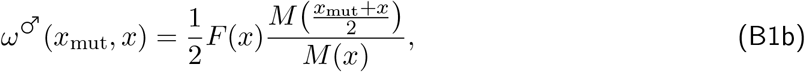

where, in both, the factor 1/2 accounts for Mendelian segregation, and (*x*_mut_ + *x*)*/*2 is the allocation to female function by a *x*_mut_*/x* heterozygote. Since we assume no pollen limitation, the number of heterozygotes produced through female function, *ω*_♀_(*x*_mut_, *x*) (eq. B1a), is simply half the number of ovules produced. By contrast, the number of heterozygotes sired through male function *ω*_♂_(*x*_mut_, *x*) (eq. B1b) depends on the number of resident ovules *F* (*x*) fertilised by mutant pollen through competition with resident pollen (with success given by the mutant’s relative contribution to the pollen pool, i.e. by *M* ([*x*_mut_ + *x*]*/*2)*/M* (*x*)). After density regulation, the number of heterozygotes is then given by

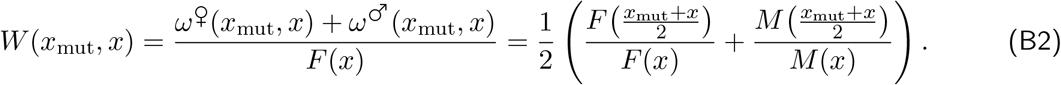

Equation (B2) corresponds to the diploid version of the haploid invasion fitness used in traditional sex allocation theory, which is sometimes called the Shaw-Mohler equation (Shaw and Mohler, 1953), see eq. 14.3 on p. 221 in Charnov (1982).

#### B.1.2 Evolution in two phases

When mutations are rare with small phenotypic effects, trait evolution can be decomposed into two phases (Metz et al., 1996; Geritz et al., 1998; Dercole and Rinaldi, 2008; Avila and Mullon, 2023). First, the population evolves gradually under directional selection while remaining largely monomorphic (i.e., there is little genetic variance in the population). The population may thus attain a “convergence stable strategy”, which is an attractor of directional selection. Once the population expresses such a strategy, it either experiences stabilising selection and remains monomorphic, or it experiences disruptive selection and becomes polymorphic in a process referred to as “evolutionary branching” (Geritz et al., 1998). Since we are interested in the evolution of polymorphic sexual systems, we are particularly interested in the conditions that lead to disruptive selection.

##### B.1.2.1 Directional selection

Directional selection is given by the selection gradient,

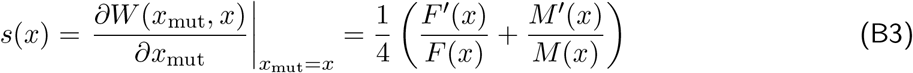

(where we used eq. B2), whose sign tells us about the direction favoured by selection in a resident population expressing *x*. Specifically, selection favours an increase in allocation to female function when *s*(*x*) > 0, and an increase in allocation to male function when *s*(*x*) < 0. A singular strategy *x*^*^ such that directional selection ceases to act is defined as

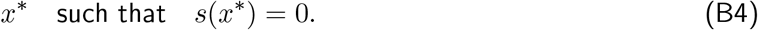

For our model (using eq. B3), such a singular strategy *x*^*^ is characterised by,

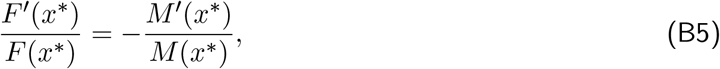

i.e., where the relative loss of fecundity through one sex is exactly compensated by the gain in the other. Assuming power gain curves (eq. A4), eq. (B5) leads to a single singular strategy

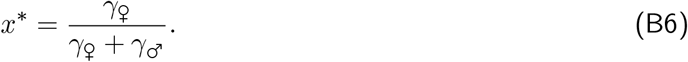

When gain curves have the same shape (*γ*_♀_ = *γ*_♂_), the singular strategy is equal allocation to male and female functions (*x*^*^ = 1*/*2). Otherwise, the singular strategy is biased towards the sex with the more accelerating gain curve (i.e. *x*^*^ > 1*/*2 when *γ*_♀_ *> γ*_♂_ or *x*^*^ < 1*/*2 when *γ*_♀_ *< γ*_♂_), as this leads to greater fitness returns through that sex.

The population converges to a singular strategy *x*^*^ through gradual evolution when

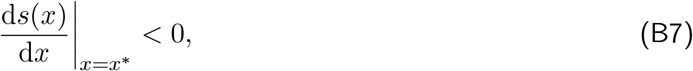

in which case *x*^*^ is said to be convergence stable. In our model (using eqs. B3 and B5), this is when

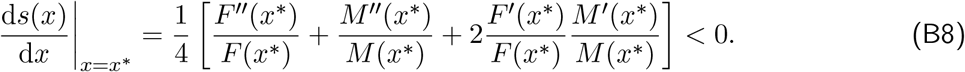

This condition holds true provided gain curves are not too accelerating relative to their increase close to the singular point (note from eq. A3 that the product *F* ^*′*^(*x*^*^)*M* ^*′*^(*x*^*^) < 0 is always negative). By construction of the model, condition eq. (B8) must be true for at least one singular point 0 *< x*^*^ < 1. This can be seen from eqs. (A2) and (A3), which imply that the selection gradient eq. (B3) becomes infinitely positive when *x* approaches 0 and infinitely negative when *x* approaches 1 (i.e. lim_*x*→0_ *s*(*x*) → ∞ and lim_*x*→1_ *s*(*x*) → −∞). Directional selection thus always pushes the population away from 0 and 1, which must then settle somewhere 0 *< x*^*^ < 1. In biological terms, selection necessarily favours investment to male function in a population of females and to female function in a population of males. In fact, assuming power gain curves (eq. A4), condition eq. (B8) (using eq. B6) becomes

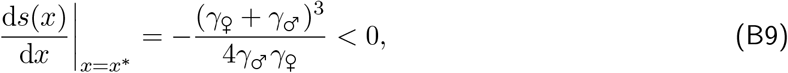

meaning that the unique singular strategy given by eq. (B6) is always convergence stable.

##### B.1.2.2 Disruptive selection

Once the population has converged to singular strategy *x*^*^ (i.e., satisfying eqs. B4 and B7), selection is disruptive and favours polymorphism when

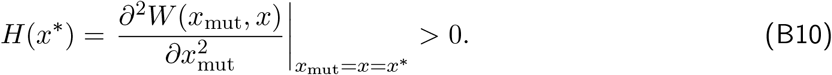

Otherwise, selection is stabilising so that the population remains unimodally distributed around *x*^*^. Plugging eq. (B2) into eq. (B10), we find that, for polymorphism to occur in our model, it is necessary that

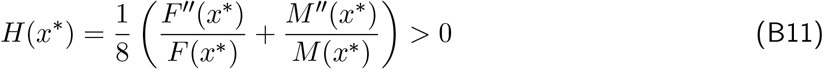

i.e., that at least one gain curve is sufficiently accelerating (while the other is not too saturating).

With power gain curves (eq. A4), condition (B11) becomes

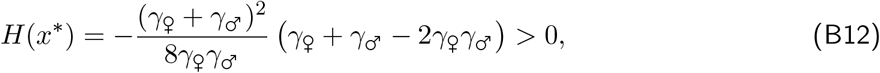

which is true when,

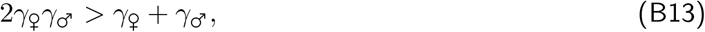

as shown in Figure S2 (darker grey region).

This condition (B13) reveals that when both gain curves increase more than linearly (*γ*_♀_ > 1 and *γ*_♂_ > 1), polymorphism emerges. Otherwise, we can rearrange eq. (B13) as

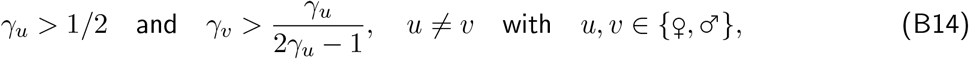

which shows, that provided the smaller gain curve exponent *γ*_*u*_ is greater than 1*/*2 (i.e., the gain curve is not too saturating), polymorphism is favoured when the other gain curve is sufficiently accelerating. As pointed out by a reviewer, Condition (B13) can also be rearranged to the harmonic mean of the gain curve exponents being greater than one, i.e.

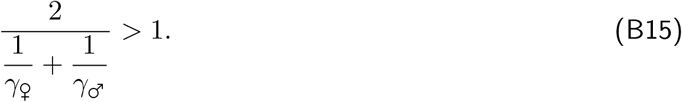

### B.2 Stable polymorphism in sex allocation alleles

The above analysis reveals whether polymorphism emerges, in which case two alleles, say *x*_1_ and *x*_2_, coding for different sex allocation strategies coexist in the population. To characterise these two alleles at evolutionary equilibrium, we need to consider the invasion fitness of a mutant allele *x*_mut_ in a population where *x*_1_ and *x*_2_ are both common and at equilibrium. Those values of *x*_1_ and *x*_2_ for which no other allele can invade then constitute an evolutionarily stable polymorphism (or coalition, Vincent and Brown, 2005; Metz, 2011).

#### B.2.1 Equilibrium resident population

We first characterise the equilibrium of a resident population in which alleles *x*_1_ and *x*_2_ are both common, i.e., the equilibrium number of *x*_1_*/x*_1_ and *x*_2_*/x*_2_ homozygotes and *x*_1_*/x*_2_ heterozygotes. To that end, let *n*_11,*t*_(*x*_1_, *x*_2_), *n*_22,*t*_(*x*_1_, *x*_2_) and *n*_12,*t*_(*x*_1_, *x*_2_), respectively, be the number of *x*_1_*/x*_1_ and *x*_2_*/x*_2_ homozygotes and *x*_1_*/x*_2_ heterozygotes at some generation *t*. Since population size *N* is constant, we always have

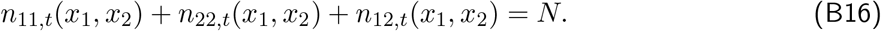

We can thus write

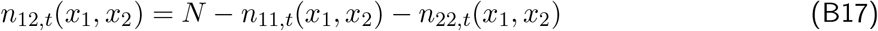

for the number of heterozygotes and focus on the dynamics of the homozygotes.

To specify *n*_11,*t*+1_(*x*_1_, *x*_2_) and *n*_22,*t*+1_(*x*_1_, *x*_2_), it is useful to define 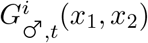 as the number of male gametes and 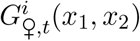 as the number of female gametes bearing allele *x*_*i*_ (for *i* ∈ {1, 2}) at generation *t*. These are given by

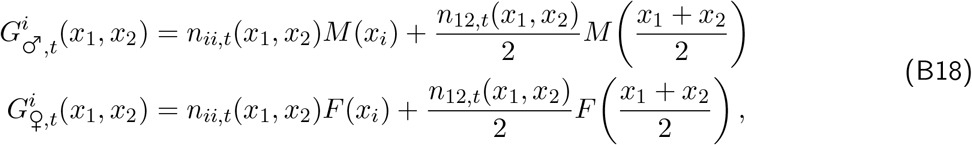

where the first term of each equation is the number of gametes produced by homozygotes and the second by heterozygotes. Further, let

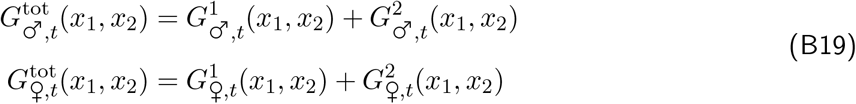

be the total number of male and female gametes produced at generation *t*, respectively.

Using the above notation, the number of *x*_1_*/x*_1_ and *x*_2_*/x*_2_ homozygotes at generation *t* + 1 can be written as the product of three factors,

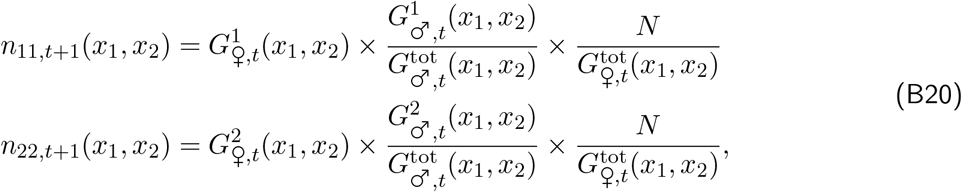

which can be understood as follows. The first factor corresponds to the number of ovules carrying allele *x*_*i*_ (*i* ∈ {1, 2}) produced at generation *t*, while the second factor is the probability that these ovules are fertilised by pollen carrying allele *x*_*i*_. The product of these two terms thus gives the number of zygotes with genotype *x*_*i*_*/x*_*i*_ produced at generation *t*. The third factor is the probability that a zygote is recruited, which is equal to the number of breeding spots available (*N*) divided the total number 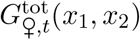 of zygotes competing for them.

The equilibrium number of *x*_1_*/x*_1_ and *x*_2_*/x*_2_ homozygotes, 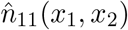 and 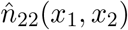, respectively, are then obtained by solving

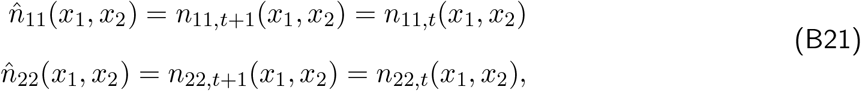

which in turn gives,

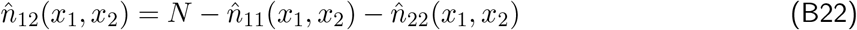

for the equilibrium number of heterozygotes. We do not solve these equilibria analytically as it is too complicated but use the above equations in our upcoming numerical analysis.

#### B.2.2 Invasion fitness and polymorphic equilibrium

Since the resident population now has two common alleles, *x*_1_ and *x*_2_, the rare mutant *x*_mut_ can be found in two heterozygous forms, *x*_1_*/x*_mut_ and *x*_2_*/x*_mut_. We refer to these forms as ‘classes’ and let *x*_1_*/x*_mut_ and *x*_2_*/x*_mut_ be class 1 and 2, respectively. The dynamics of the mutant can then be modelled by the matrix equation

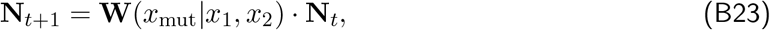

where **N**_*t*_ = {*N*_1,*t*_, *N*_2,*t*_} is a vector containing the number of mutants in classes 1 (*N*_1,*t*_) and 2 (*N*_2,*t*_) at some generation *t*, and **W**(*x*_mut_|*x*_1_, *x*_2_) is a 2 *×* 2 matrix whose (*i, j*)-entry *w*_*ij*_(*x*_mut_|*x*_1_, *x*_2_) is the expected number of successful mutants of class *i* produced by a mutant of class *j*.

To specify the fitness matrix **W**(*x*_mut_|*x*_1_, *x*_2_), let us denote for *i* ∈ {1, 2}

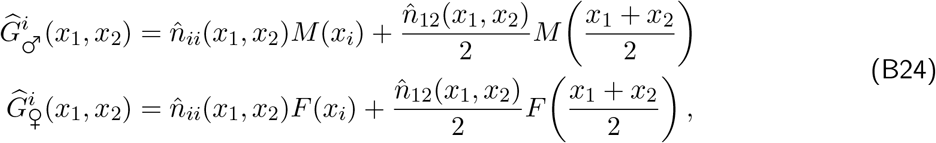

the number of pollen grains and ovules carrying allele *x*_*i*_ produced in the resident population at equilibrium (using eq. B18), and similarly let

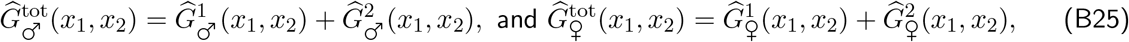

denote the total number of male and female gametes produced in the resident population at equilibrium.

Using the above notation, the (*i, j*)-entry *w*_*ij*_(*x*_mut_|*x*_1_, *x*_2_) of **W**(*x*_mut_|*x*_1_, *x*_2_) can be written as

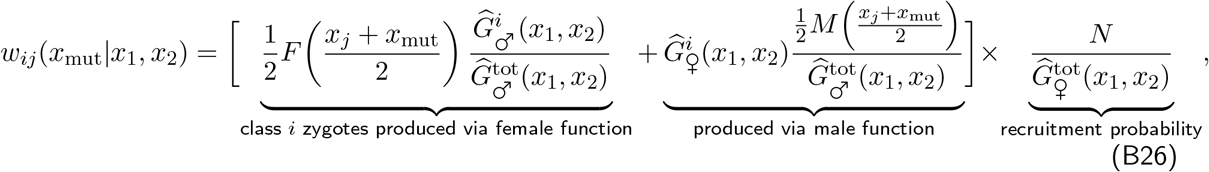

which can be understood as follows. The term between square brackets gives the number of zygotes of class *i* produced by a mutant of class *j*. It is composed of two summands. The first summand corresponds to class *i* zygotes produced through the female function. To produce such an offspring, a mutant must contribute an ovule carrying allele *x*_mut_, which corresponds to half of its total ovule production (*F* ([*x*_*j*_ + *x*_mut_]*/*2)*/*2), and receive pollen carrying allele *x*_*i*_, which occurs with a probability given by the proportion of *x*_*i*_ pollen in the total pollen pool 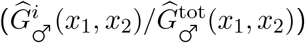. The second summand corresponds to class *i* zygotes produced through the male function, i.e., to the *x*_*i*_ ovules produced by residents 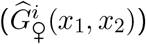, which are fertilised by pollen of the mutant carrying *x*_mut_. The number of successful offspring is then obtained by multiplying the number of zygotes produced by the probability that they will be recruited, which is given by the number of breeding spots available (*N*) divided by the total number of zygotes competing for them, 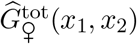.

The leading eigenvalue *ρ*(*x*_mut_|*x*_1_, *x*_2_) of the matrix **W**(*x*_mut_|*x*_1_, *x*_2_) is the invasion fitness of the mutant allele *x*_mut_, which can thus be used to characterise the evolutionary dynamics of the dimorphic population (Geritz et al., 1998). In particular, the derivatives of *ρ*(*x*_mut_|*x*_1_, *x*_2_) with respect to *x*_mut_ evaluated at *x*_mut_ = *x*_1_ and *x*_mut_ = *x*_2_, i.e.,

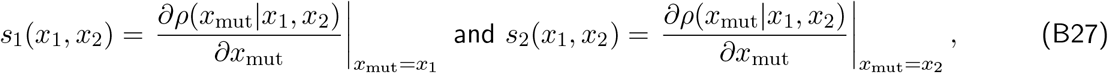

are the selection gradients acting on allelic values *x*_1_ and *x*_2_, respectively. These gradients indicate the direction of selection on *x*_1_ and *x*_2_ in a resident population where *x*_1_ and *x*_2_ are at equilibrium. For instance, in a resident population where *x*_1_ *> x*_2_ and *s*_1_(*x*_1_, *x*_2_) > 0 and *s*_2_(*x*_1_, *x*_2_) < 0, selection favours greater divergence between the sex allocation strategies of homozygotes (i.e., an increase in *x*_1_ and a decrease in *x*_2_). Conversely, selection favours lower divergence where *s*_1_(*x*_1_, *x*_2_) < 0 and *s*_2_(*x*_1_, *x*_2_) > 0 (i.e., a decrease in *x*_1_ and an increase in *x*_2_).

#### B.2.3 Numerical analysis

We study eq. (B27) numerically, assuming power gain curves (eq. A4). For different sets of parameters (i.e., different values of *γ*_♀_ and *γ*_♂_), we compute the selection gradients **s**(*x*_1_, *x*_2_) = (*s*_1_(*x*_1_, *x*_2_), *s*_2_(*x*_1_, *x*_2_)) for many pairs (*x*_1_, *x*_2_) across phenotypic space (0 ⩽ *x*_1_ ⩽ 1 and 0 ⩽ *x*_2_ ⩽ 1). The vector **s**(*x*_1_, *x*_2_) points in the direction favoured by selection in a population in which alleles *x*_1_ and *x*_2_ segregate. Repeating this operation for many (*x*_1_, *x*_2_) pairs across the phenotype space, we obtain a vector field that determines the evolutionary trajectories favoured by selection in phenotype space and where allelic values converge to. We denote by 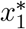 and 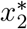 these equilibria.

Inspection of these vector fields for different values of *γ*_♀_ and *γ*_♂_ reveals four possible evolutionary outcomes (summarised in Fig. S3, see Fig. S4 for examples). **(i)** When condition (B13) is not satisfied (blue region in Fig. S3), selection favours a single intermediate allelic value *x*_1_ = *x*_2_ = *x*^*^ so that hermaphroditism is maintained, as expected (Fig. S4A). By contrast, when condition (B13) holds, three types of polymorphic equilibrium emerge: **(ii)** If the female gain curve is saturating and the male gain is sufficiently accelerating (1*/*2 *< γ*_♀_ < 1 and *γ*_♂_ *> γ*_♀_*/*(2*γ*_♀_ − 1), green region in Fig. S3), one allele encodes full allocation to the female function while the other encodes a male-biased hermaphroditic strategy (‘gynodioecy’, i.e., 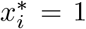 and 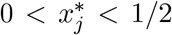 for *I* ≠*j*; Fig. S4B); **(iii)** Conversely, if the male gain curve is saturating and the female gain curve is sufficiently accelerating (1*/*2 *< γ*_♂_ < 1 and *γ*_♀_ *> γ*_♂_*/*(2*γ*_♂_ − 1); yellow region in Fig. S3), one allele encodes a female-biased hermaphroditic strategy while the other encodes a pure male strategy (‘androdioecy’, i.e., 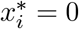 and 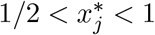 for *i* ≠ *j*; Fig. S4C); **(iv)** Finally, when both gain curves are accelerating (*γ*_♀_ > 1 and *γ*_♂_ > 1; red region in Fig. S3), two allele coexist, one that encodes a pure female and another a pure male strategy (‘dioecy’, i.e., 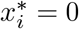 and 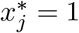 for *i* ≠ *j*; Fig. S4D).

To explore cases **(ii)** and **(iii)** further, we calculated the equilibrium reached by the population under the assumption that one of the alleles encodes a unisexual strategy, which substantially reduced computation time and allowed us to investigate a greater parameter range. For case **(ii)**, we first assume that allele *x*_1_ encodes full female allocation (*x*_1_ = 1), and compute the selection gradient acting on allele *x*_2_, *s*_2_(1, *x*_2_) to determine 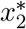 as follows

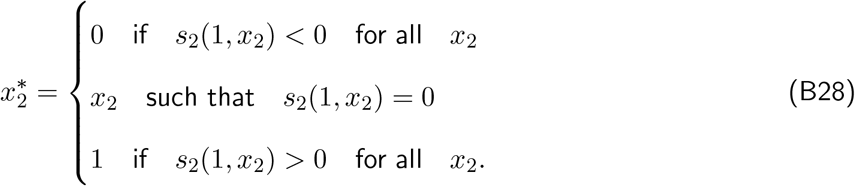

Given 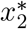, we in turn compute the selection gradient acting on allele *x*_1_, i.e. 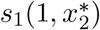. If 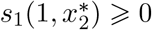, then 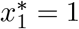 is favoured and 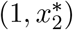 constitutes a stable equilibrium. Otherwise, 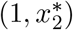 is unstable and selection will lead the population away from it. We repeat the same analysis for case **(iii)** by first setting *x*_1_ = 0, which allows us to determine whether an equilibrium involving males and hermaphrodites is stable. The results of this analysis are reported in Fig. S5A and S5B for cases **(ii)** and **(iii)**. These show that, in both cases, the hermaphroditic strategy is strongly biased towards the opposite sex to unisexuals, i.e., it is strongly male-biased in case **(ii)** and strongly female-biased in case **(iii)**.

### B.3 Individual-based simulations

To accompany the mathematical analysis presented above, we ran individual-based simulations of our model. These simulations assume that alleles are additive at the sex allocation locus, and their output is shown in Figure 2B-E in the main text (we extend these simulations to include dominance evolution in Appendix C.1).

The program was coded in C++11 (see files attached to submission for the code). It simulates a diploid population of constant size *N* (see captions in Fig. 2 for parameter values). Individuals *i* ∈ {1, …, *N* } are characterised by their diploid genotype (*x*_*i*1_, *x*_*i*2_) at the sex allocation locus. The population is initially fixed for an arbitrary sex allocation allele *x*_0_ ∈ (0, 1). We then let the population evolve for *t*_max_ = 50, 000 generations, and record genotypes every *t*_mes_ = 100 generations. For each generation, we begin by determining the sex allocation strategy expressed by individuals. For individual *i*, its strategy *x*_*i*_ is given by

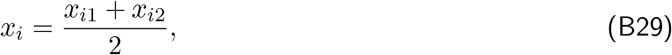

i.e., alleles at the sex allocation locus are additive. From this, we determine the male and female fecundities of each individual *M* (*x*_*i*_) and *F* (*x*_*i*_) using eq. A4 (see figure captions for parameter values). We form the next generation by sampling *N* fathers and *N* mothers with replacement from the population, with probabilities weighted by the individuals’ male and female fecundities, respectively. Parents then each transmit one of their sex allocation alleles to their offspring with equal probability to create new diploid individuals (Mendelian segregation). Each allele has a probability *µ* = 0.005 to mutate, in which case its mutated value is sampled in a Gaussian distribution centered on the parental value with standard deviation *σ* = 0.01, truncated such that values are kept within bounds (*x* ∈ [0, 1]). In other words, for a parental allele encoding strategy *x*_par_ undergoing mutation, the phenotypic value *x*_off_ encoded by the mutated allele is given by

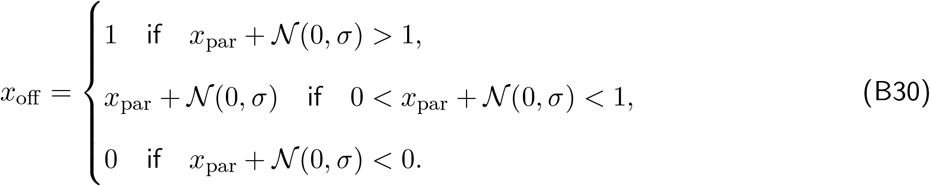

Note that using *σ* = 0.01 entails that essentially all mutations have small effects on the phenotype encoded by alleles. For instance, the probability that a mutation will cause a change with an absolute value greater that 0.05 is of order 10^−7^. Once the *N* offspring are produced, parents are replaced by the new generation (i.e., there is no generation overlap).

### B.4 Connection with fitness sets and fecundity trade-offs

Our results align with those from classical sex allocation theory (Charnov et al., 1976), which relies on the notion of a ‘fitness set’. The fitness set relates female to male fecundity (scaled by their respective maxima). Using the notation of our model, it is given by the function *φ* defined such that

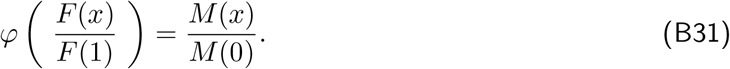

Classical theory studies the ability of fixed combinations of sexual phenotypes to invade one another depending on the shape of the fitness set. It demonstrates that hermaphroditism is maintained when the fitness set is convex (*φ*^*′′*^(*x*) < 0), which corresponds to a case where fecundity trade-offs are weak in both sexes, i.e., where a pure male starting to allocate resources to female gametes reaps a higher female fecundity benefit than the cost it incurs in terms of male fecundity (and vice-versa). In our model, this corresponds to both gain curves being saturating (*F* ^*′′*^(*x*) < 0 and *M* ^*′′*^(*x*) < 0; dashed line in Fig. S6). In contrast, dioecy is maintained if the fitness set is concave (*φ*^*′′*^(*x*) > 0). This corresponds to strong trade-offs in both sexes, i.e., the fecundity cost paid for deviating from a pure strategy is higher than the fecundity benefits gained in the other sex, which in our model occurs when both gain curves are accelerating (*F* ^*′′*^(*x*) > 0 and *M* ^*′′*^(*x*) > 0; solid line in Fig. S6; see Fig. 1 in Charnov et al., 1976). Finally, sexual systems involving a mixture of hermaphrodites and unisexual females or males can be maintained if the fitness set is convex-concave or concave-convex (i.e., when *φ*^*′′*^(*x*) changes sign between 0 and 1), respectively, corresponding to asymmetrical trade-offs (i.e., deviating from a pure strategy is beneficial in one sex, but not the other). In our model, this occurs when one gain curve is saturating and the other is accelerating (*F* ^*′′*^(*x*)*M* ^*′′*^(*x*) < 0; dotted and dot-dashed lines in Fig. S6; see Fig. 2 in Charnov et al., 1976).

Our results thus extend classical theory by showing that gradual evolution will lead populations to these endpoints. In addition, classical theory focuses on determining optimal sexual systems at a phenotypic level (the ‘phenotypic gambit’, Grafen, 1991), overlooking the genetic architecture underlying them. Our work thus further extends classical theory by revealing how gradual evolution can shape the genetic architecture of sexual systems in diploid populations (in Appendix C) and how this interacts with partial selfing and inbreeding depression (in Appendix D).

### B.5 Connection with population genetics models

We have demonstrated that a genetic polymorphism of sex allocation strategies can be established by mutants with small effects, which then diverge gradually under the action of disruptive selection. At first glance, these results may appear to be at odds with previous population genetics analyses that concluded that mutations with large effects on sex allocation are necessary for the establishment of a genetic polymorphism (Charlesworth and Charlesworth, 1978b). The analyses of Charlesworth and Charlesworth (1978b) include partial selfing and inbreeding depression (which we consider in Appendix D), but the apparent discrepancy between our results and those of Charlesworth and Charlesworth (1978b) can be addressed assuming complete outcrossing (partial selfing can be included in the arguments below, and this leads to the same conclusions, but it makes the mathematics much more tedious and harder to follow).

For the sake of clarity, let us first briefly go over the results obtained by Charlesworth and Charlesworth (1978b) under complete outcrossing. We therefore consider a large population of annual hermaphrodites with a female fecundity arbitrarily set to one and a male fecundity set to *b* ∈ ℝ. Each generation, we individuals develop and mature sexually, then mate randomly with one another and die after setting seed. The next generation is then formed by sampling juveniles from the seeds produced that year. We consider the fate of a partially male-sterile mutant, such that its female fecundity is 1 + *k* and its male fecundity is 1 − *K*. The invasion fitness of this mutant is given by,

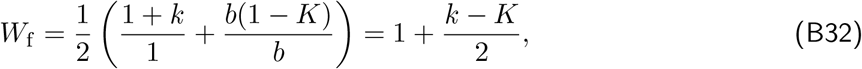

which corresponds to eq. (3) in Charlesworth and Charlesworth (1978b) with *s*_f_ = *s* = 0. If

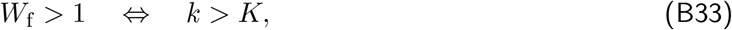

then the mutant may invade. To determine whether this establishes a genetic polymorphism instead of reaching fixation, we then consider the ability of the hermaphrodite to invade a population consisting only of female-biased individuals (*females* for short throughout this section). The invasion fitness of the hermaphrodite is given by

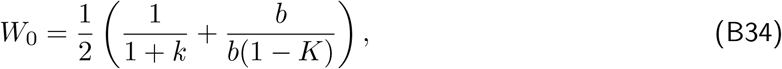

which shows that the hermaphrodite may invade when

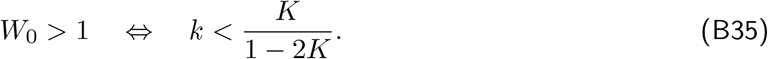

Combining eqs. (B33) and (B35) together, we obtain that for a genetic polymorphism to be established, *k* and *K* must be such that

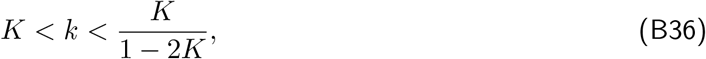

which corresponds to the condition given on p. 142 in Charlesworth and Charlesworth (1978b) with *s* = 0. This condition is plotted Fig. S7, which shows that the space of parameters *k* and *K* that lead to polymorphism gets smaller as *k* and *K* get small. If one assumes that a mutation is equally likely to cause any possible combination of *k* and *K*, i.e. if one assumes that when a mutation occurs, its effects on female and male fecundity are uniformly and independently distributed, one may therefore conclude that a genetic polymorphism is unlikely to be due to a small-effect mutation (Charlesworth and Charlesworth, 1978b).

In contrast, when a mutation occurs in our model, its influence on female and male fecundity are determined by its effect on sex allocation *x* and in turn by the shape of gain curves *F* (*x*) and *M* (*x*) (i.e., the distributions of *k* and *K* are neither uniform nor independent). To be more specific about this, let *x* be the sex allocation strategy of the resident hermaphrodite. Let *x*_f_ *> x* denote the sex allocation strategy of the female-biased mutant. Then, the parameters *k* and *K* as defined in Charlesworth and Charlesworth (1978b) for this model are given by

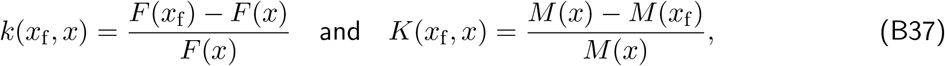

respectively.

Performing the same invasion analysis as in Charlesworth and Charlesworth (1978b) using eq. (B37) then recovers our exact same results concerning directional and disruptive selection (from Appendix B.1). Indeed, assume that the mutant has a weak phenotypic effect, i.e. *x*_f_ = *x* + *δx* where *δx* is small, so that we can write eq. (B37) as,

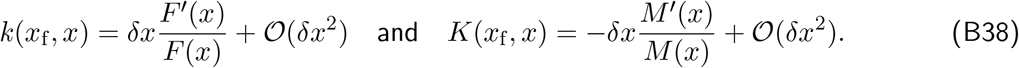

Plugging these expressions in eq. (B32) and solving for condition (B33), we obtain that the female-biased mutant can invade the initial hermaphrodite if

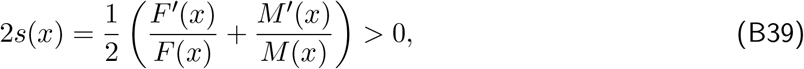

where *s*(*x*) is the selection gradient on sex allocation we derived earlier (eq. B3). Conversely, plugging eq. (B38) into eq. (B35), we obtain that a hermaphrodite expressing *x* can invade a population fixed for *x*_f_ when

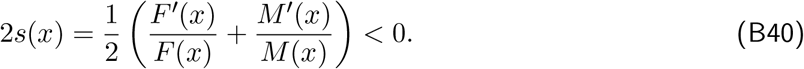

These results demonstrate that where *s*(*x*) ≠0, i.e. where the strategy expressed by individuals is away from the singular strategy *x*^*^ (eq. B6), any mutant with a weak effect arising in a hermaphroditic population will either be purged or sweep to fixation owing to directional selection. Through a sequence of substitutions, however, the population will eventually fix *x*^*^, at which point *s*(*x*^*^) = 0.

Once the population expresses the singular strategy *x*^*^ and directional selection ceases, disruptive selection is revealed by considering higher order effects in *δx*. Plugging eq. (B37) into eqs. (B32) and (B34) with *x* = *x*^*^ yields,

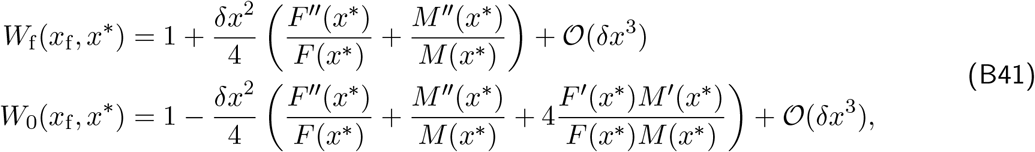

for the invasion fitness of the female and hermaphrodite types, respectively. From these two equations, it is straightforward to derive the conditions for a genetic polymorphism using the same method as in Charlesworth and Charlesworth (1978b), i.e. the conditions for *W*_f_ (*x*_f_, *x*^*^) > 1 and *W*_0_(*x*_f_, *x*^*^) > 1. We find these are

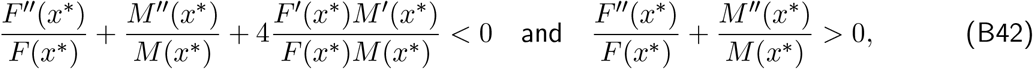

which are equivalent to the conditions we derived earlier for polymorphism using our adaptive dynamics approach (eqs. B8 and B11). This shows that where our model predicts polymorphism, the condition obtained by Charlesworth and Charlesworth (1978b) are satisfied (i.e. eq. B36 is satisfied). We further show in Appendix B.2 that once two alleles that initially encode weakly differentiated phenotypes around *x*^*^ have established a polymorphism, these two alleles then become increasingly divergent as they each accumulate further mutations.

## Appendix C Emergence of XY and ZW sex determination under complete outcrossing

Here, we investigate the joint evolution of sex allocation with its underlying genetic architecture under complete outcrossing. We derive the results presented in main text sections “Emergence of XY and ZW sex determination through dominance evolution” and “Competition through male and female functions determines whether XY or ZW evolves”.

### C.1 Simulating dominance evolution through gene expression evolution

First, we extend our model by allowing dominance and sex allocation to co-evolve in an individual-based simulation (used to generate simulation results in Fig. 3; see files attached to submission for the code). The basic structure of the simulation remains unchanged from Appendix B.3, but the sex allocation phenotype expressed by individuals is now assumed to be determined by their genotype at a locus made of two fully-linked elements, a sex allocation gene and its promoter. Alleles at the sex allocation locus are expressed in proportion to the affinity of their promoter for transcription factors (Van Dooren, 1999). Specifically, we assume that the phenotype *x*_*i*_ of individual *i* bearing genotype *g*_*i*_ = {(*a*_*i*1_,*x*_*i*1_), (*a*_*i*2_,*x*_*i*2_)} is given by

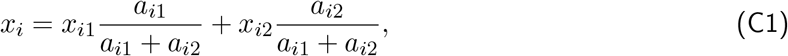

where *x*_*i*1_ and *x*_*i*2_ ∈ [0, 1] are alleles at the sex allocation gene, and *a*_*i*1_ and *a*_*i*2_ ∈ (0, +∞) are its alleles at the promoter (a positive lower bound is imposed on promoter affinity *a* to avoid divisions by zero). Each time an offspring is produced, its alleles at the sex allocation locus and at its promoter each have a probability *µ* = 0.005 to mutate. When an allele mutates, its new value is sampled in a Gaussian distribution centered on the parental value with standard deviation *σ* = 0.01 (truncated if necessary so that traits are kept within bounds). The population is set to be initially monomorphic for an arbitrary *x*_0_ ∈ (0, 1) at the sex allocation gene and *a*_0_ = 1 at the promoter. We then let the population evolve for *t*_max_ = 50, 000 generations, and record the phenotype and genotype of each individual every *t*_mes_ = 100 generations. Figures 3B-D in the main text were generated using this simulation program. They show that complete dominance of one allele over the other always evolves, leading to the emergence of XY and ZW sex determination.

### C.2 The co-evolution of dominance and sex allocation

To better understand the forces that lead to XY and ZW sex determination in our simulations, we analyse a more general model of the co-evolution of dominance and sex allocation in this section.

#### C.2.1 The model

We consider a model in which two sex allocation alleles, *x*_♀_ and *x*_♂_ (*x*_♀_ *> x*_♂_), segregate in the population, with gain curves *F* (*x*) and *M* (*x*) such that a singular strategy *x*^*^ exists, is convergence stable and invadable (so that polymorphism is favoured and maintained by disruptive selection). In heterozygotes, allele *x*_♀_ is expressed in proportion to a dominance coefficient *h*, which we assume to be a quantitative trait controlled by a modifier locus freely recombining with the sex allocation gene. This allows us to study how selection acts on dominance in general, beyond the promoter mechanism assumed in simulations. Similar to the sex allocation gene, we label alleles at the dominance modifier by their quantitative effect on dominance, i.e. an individual carrying alleles *h*_*i*_ and *h*_*j*_ ∈ [0, 1] at the modifier expresses a dominance coefficient

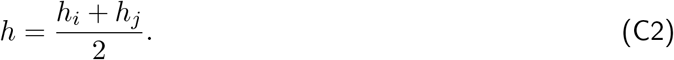

Let *x*_het_(*h*_*i*_, *h*_*j*_) denote the strategy expressed by a heterozygote at the sex allocation gene (*x*_♀_*/x*_♂_), given they carry genotype *h*_*i*_*/h*_*j*_ at the dominance modifier. According to our assumptions, this strategy is given by

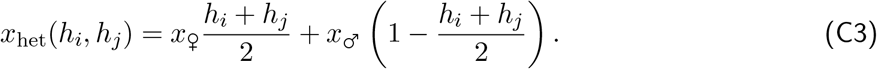

XY sex determination in a dioecious population then corresponds to the case where the population has evolved for *x*_♀_ = 1, *x*_♂_ = 0 and *h* = 0. Conversely, a dioecious population with ZW sex determination shows *x*_♀_ = 1, *x*_♂_ = 0 and *h* = 1.

To better understand the conditions that lead to XY or ZW sex determination, we study the coevolution of dominance and sex allocation, i.e. we analyse the three selection gradients *s*_h_(*x*_♀_, *x*_♂_, *h*), *s*_♀_(*x*_♀_, *x*_♂_, *h*) and *s*_♂_(*x*_♀_, *x*_♂_, *h*), acting on the dominance coefficient *h* and on the sex allocation strategies *x*_♀_ and *x*_♂_, respectively. We do so in three steps. Firstly, we characterise the equilibrium state of the resident population (i.e. the frequency of each type where *x*_♀_, *x*_♂_ and *h* are all fixed; section C.2.2). Secondly, we study the sex allocation strategies encoded by alleles *x*_♀_ and *x*_♂_ at evolutionary equilibrium, which we denote by 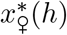 and 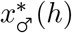, respectively, for a given dominance coefficient *h* (section C.2.3). Thirdly, we study selection on the dominance coefficient *h* in a population where alleles *x*_♀_ and *x*_♂_ encode equilibrium strategies 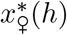 and 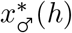 (section C.2.4).

#### C.2.2 Equilibrium state of the resident population

The resident population can be seen as a class-structured population with three classes, where each class corresponds to a genotype at the sex allocation gene (genotypes *x*_♀_*/x*_♀_, *x*_♀_*/x*_♂_ and *x*_♂_*/x*_♂_ are referred to as class 1, 2 and 3 hereafter, respectively). We denote by *n*_*i,t*_ the number of individuals in class *i* in the resident population at time *t*. Since we assume a constant population size, i.e.,

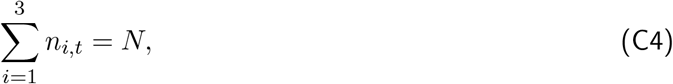

we can write the number of *x*_♂_*/x*_♂_ homozygotes as

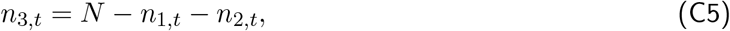

so that the dynamics of the resident population can be described by the dynamics of classes 1 and 2, i.e., of genotypes *x*_♀_*/x*_♀_ and *x*_♀_*/x*_♂_.

We denote by 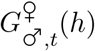 and 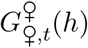 the number of male and female gametes carrying allele *x*_♀_ produced by the residents at time *t*, which are given, respectively, by

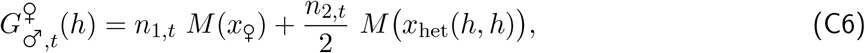

and

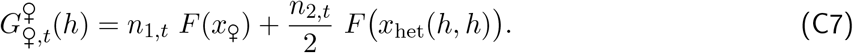

Similarly, we denote by 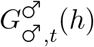 and 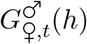 the number of male and female gametes carrying allele *x*_♂_ produced by the residents at time *t*, which are given by

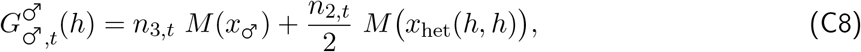

and

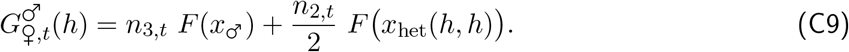

Furthermore, we denote as

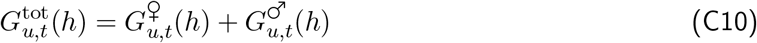

the total number of gametes of sex *u* ∈ {_♀_, _♂_} produced by residents at time *t*.

Using these expressions, the dynamics of the resident population are given by

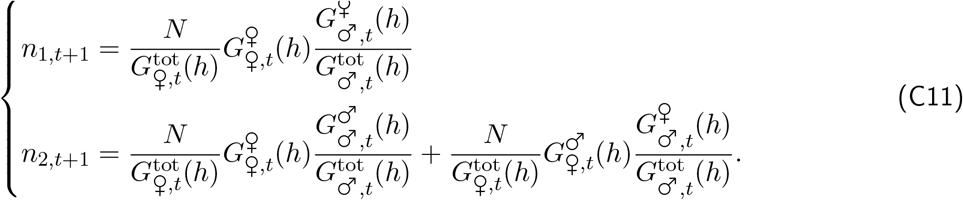

Eq. (C11) is an extension of eq. (B20) to the case of an arbitrary dominance relationship between alleles. The demographic equilibrium of the resident population is then determined by

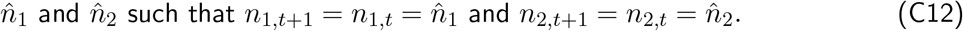

At this equilibrium, we let 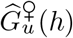 and 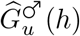 be the number of *x*_♀_ and *x*_♂_ gametes of sex *u* ∈ {_♀_, _♂_} produced by residents at equilibrium. Furthermore, we define

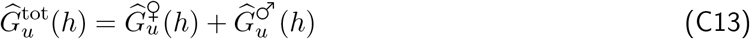

as the total number of gametes of sex *u* ∈ {_♀_, _♂_} produced by residents at equilibrium. The quantities 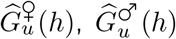 and 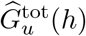 are the same as those defined in eqs. (C6) to (C10), expressed at the resident’s demographic equilibrium (i.e. with 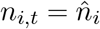 for all *i* ∈ {1, 2, 3}).

#### C.2.3 Selection on sex allocation alleles for a given dominance coefficient

We now examine the evolution of sex allocation alleles *x*_♀_ and *x*_♂_ for a given dominance coefficient *h* (rather than where *h* = 1*/*2 as in Appendix B.2). For a given pair of sex allocation strategies *x*_♀_ and *x*_♂_ for a given *h*, we first solve the recursions eq. (C11) numerically to obtain the equilibrium number of individuals of genotypes *x*_♀_*/x*_♀_, *x*_♀_*/x*_♂_ and *x*_♂_*/x*_♂_ in the resident population. We then introduce a rare mutant *x*_mut_ that influences sex allocation into this population. Depending on which sex allocation allele the mutant arises from (i.e. depending on whether *x*_mut_ derives from allele *x*_♀_ or *x*_♂_), the mutant allele has a different dominance relationship with alleles *x*_♀_ and *x*_♂_. To describe this, we denote by 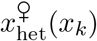 and 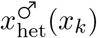 the sex allocation strategy expressed by *x*_mut_*/x*_*k*_ (where *k* ∈ {_♀_, _♂_}), heterozygotes when the mutant allele derives from allele *x*_♀_ and *x*_♂_, respectively. We assume that

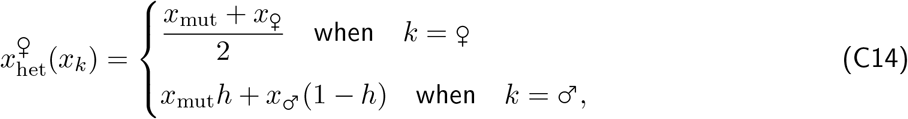

and

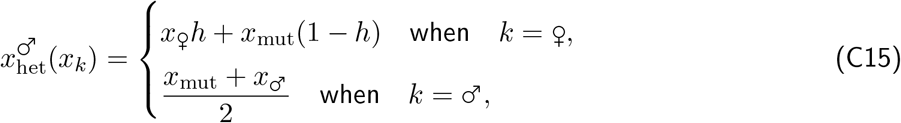

i.e. that the mutant allele retains the dominance status of its ancestor (so that it is co-dominant with the allele it arose from, which would be the case under the promoter affinity model, eq. C1).

The fate of a mutant deriving from allele *x*_*u*_ (*u* ∈ {_♀_, _♂_}) is then captured by the recurrence,

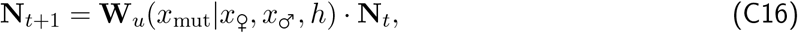

where **N**_*t*_ = {*N*_1,*t*_, *N*_2,*t*_} is a vector giving the number of mutant heterozygotes with genotypes *x*_♀_*/x*_mut_ (class 1, *N*_1,*t*_) and *x*_♂_*/x*_mut_ (class 2, *N*_2,*t*_) at time *t*, and **W**_*u*_(*x*_mut_|*x*_♀_, *x*_♂_, *h*) is a 2 *×* 2 matrix whose (*i, j*)-entry *w*_*u,ij*_(*x*_mut_|*x*_♀_, *x*_♂_, *h*) is the expected number of successful mutant heterozygotes of class *i* produced by mutant heterozygote of class *j*. Using the notations introduced in Appendix C.2.2, dropping the arguments of functions 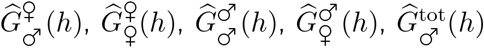 and 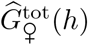 for brevity (eqs. C6 to C10), and following the same line of reasoning as in Appendix B.2.2, the fitness matrix is given by

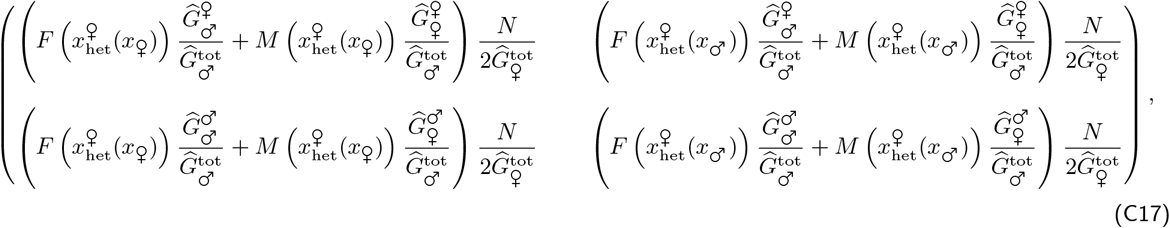

and

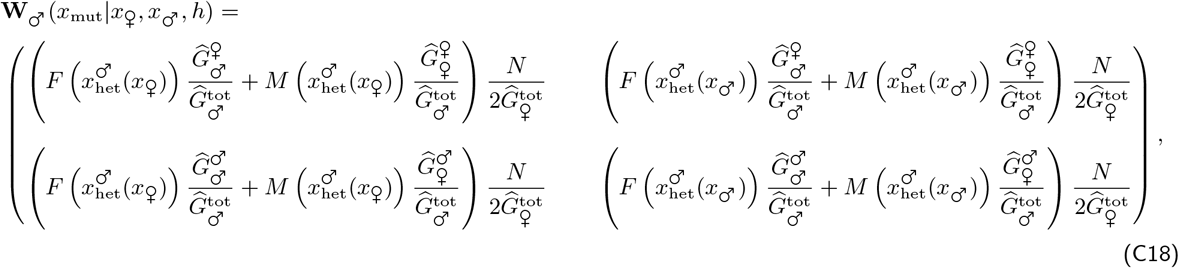

when the mutant derives from *x*_♀_ and *x*_♂_, respectively. The leading eigenvalues of **W**_♀_(*x*_mut_|*x*_♀_, *x*_♂_, *h*) and **W**_♂_(*x*_mut_|*x*_♀_, *x*_♂_, *h*), which we denote as *ρ*_♀_(*x*_mut_|*x*_♀_, *x*_♂_, *h*) and *ρ*_♂_(*x*_mut_|*x*_♀_, *x*_♂_, *h*), give the invasion fitness of mutants deriving from allele *x*_♀_ and *x*_♂_, respectively.

The selection gradients on alleles *x*_♀_ and *x*_♂_, which we write as *s*_♀_(*x*_♀_, *x*_♂_, *h*) and *s*_♂_(*x*_♀_, *x*_♂_, *h*), are obtained from these eigenvalues:

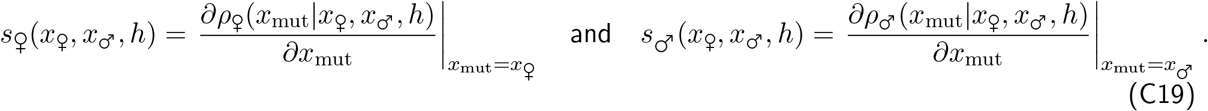

From these selection gradients, we follow the same approach as described in B.2.3 to determine how sex allocation alleles *x*_♀_ and *x*_♂_ evolve and where they converge to as a function of *h*. We denote their equilibrium as 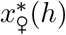 and 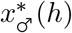, respectively. We find that when both gain curves are accelerating, the two alleles encode pure female and male strategies 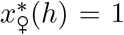and 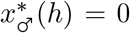, irrespective of *h*.

When one of the gain curves saturates, we find that changes in the dominance coefficient *h* have a small effect on the hermaphroditic strategy in every investigated case (Fig. S8). When the male gain curve saturates, a pure male allele 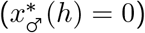 and an allele encoding a female-biased hermaphroditic strategy 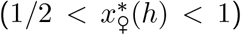 are maintained (androdioecy, Fig. S8A). When the female gain curve saturates, a pure female allele 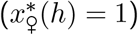 and an allele encoding a male-biased hermaphroditic strategy 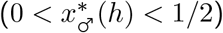 coexist for all *h* (gynodioecy, Fig. S8B).

#### C.2.4 Selection on dominance

We can now study selection on the dominance coefficient *h*, given that sex allocation alleles encode equilibrium strategies 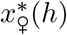 and 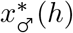.

The dynamics of a rare mutant *h*_mut_ at the dominance modifier can be modelled as

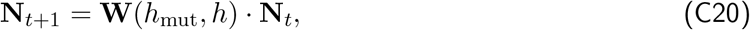

where the entries of **N**_*t*_ = (*N*_1,*t*_, *N*_2,*t*_, *N*_3,*t*_) give the number of mutant heterozygotes at the dominance modifier in each class at time *t*, i.e., the number of mutant heterozygotes *h*_mut_*/h* at the dominance modifier with genotype *x*_♀_*/x*_♀_ (*N*_1,*t*_), *x*_♀_*/x*_♂_ (*N*_2,*t*_) and *x*_♂_*/x*_♂_ (*N*_3,*t*_) at the sex allocation locus; and **W**(*h*_mut_, *h*) is a 3 *×* 3 matrix whose (*i, j*)-entry *w*_*ij*_(*h*_mut_, *h*) gives the expected number of successful mutant heterozygote offspring of class *i* produced by a mutant heterozygote of class *j* over one iteration of the life cycle.

Using eqs. (C6) to (C10), the three columns **w**_*i*_ (for *i* ∈ {1, 2, 3}) of the matrix **W**(*h*_mut_, *h*) are given by

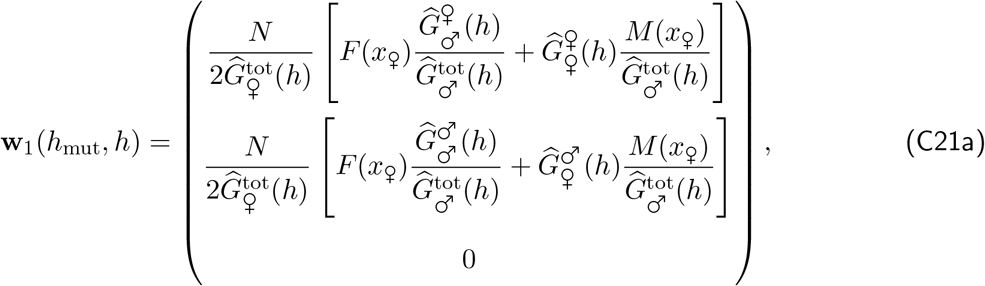

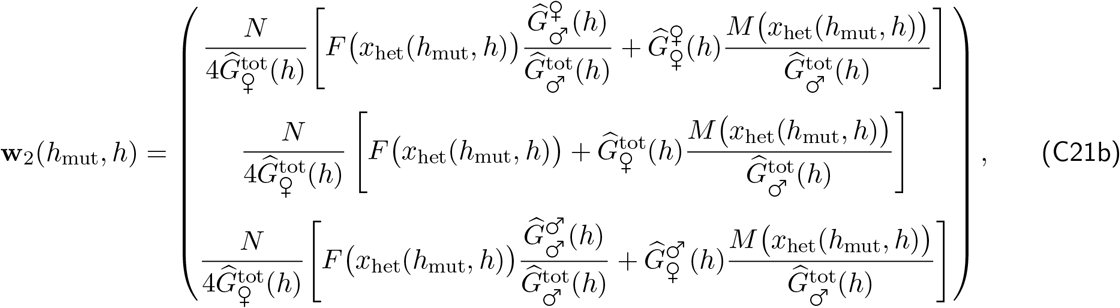

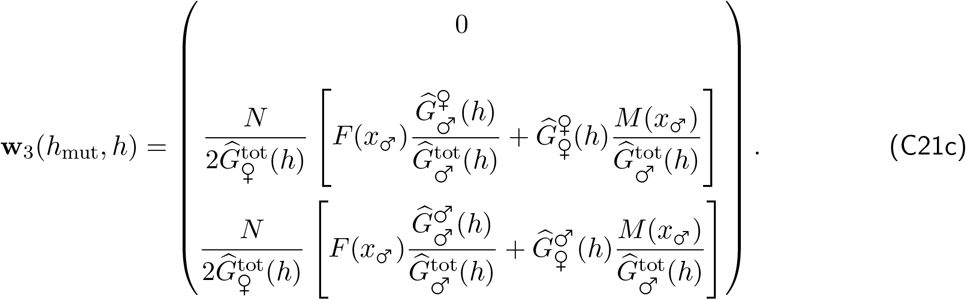

The entries of **W**(*h*_mut_, *h*) given in eq. (C21) were computed following a similar reasoning to that used to obtain eq. (B26) in Appendix B.

##### Mutant class-specific frequencies

To compute the selection gradient on dominance, let us first introduce **q**^*°*^(*h*), the right eigenvector of **W**^*°*^(*h*) = **W**(*h, h*) (the **W** matrix under neutrality), normalised so that

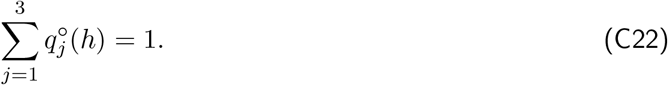

Its *j*^th^ element 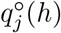 corresponds to the asymptotic frequency of mutants in class *j* under neutrality, which is equivalent to the frequency of genotypes class *j* in the resident population in this case. In other words, we have

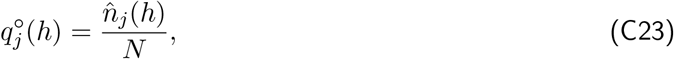

for all *j* ∈ {1, 2, 3}, where 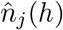 is given by eq. (C12).

##### Class-specific fitness effects

Second, we denote as **D**(*h*) the 3 *×* 3 matrix that contains the first order derivatives of the elements of **W** with respect to *h*_mut_, evaluated at *h*_mut_ = *h*, i.e. its (*i, j*)-entry is given by

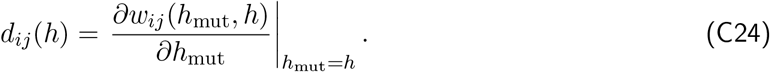

Each element *d*_*ij*_(*h*) gives the effect the mutant allele has on the expected number of class *i* offspring produced by a class *j* individual.

**Class reproductive values**

Finally, we let **v**^*°*^(*h*) be the left eigenvector of **W**^*°*^(*h*) normalised such that **v**^*°*^(*h*) *·* **q**^*°*^(*h*) = 1, which collects the reproductive values of individuals in each class in the resident population at equilibrium. The reproductive value 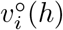 gives the asymptotic demographic contribution of an individual from class *i* relative to individuals from other classes, i.e. reproductive values capture the relative influence of individuals from each class on the long-term demography of the population (for more details, see p. 27 in Fisher, 1930; p. 37 (eq. 1.54a) in Charlesworth, 1980; section 4.6 starting on p. 92 in Caswell, 2001; p. 153 in Rousset, 2004).

Using the quantities defined above, the selection gradient on dominance, *s*_h_(*x*_♀_, *x*_♂_, *h*), is given by,

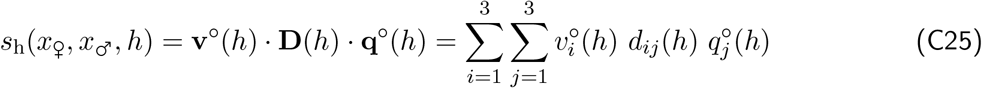

(eq. A2 in Taylor and Frank (1996), see also eq. 12.48a in Otto and Day (2011), Caswell, 2001; Taylor, 1990; Avila and Mullon, 2023). Eq. (C25) is most easily read right to left. The *j*^th^ element of 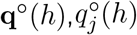, is the probability that a randomly sampled resident individual belongs to class *j*, which can be thought of as the probability that a mutation appears in a resident individual of that class. The element *d*_*ij*_(*h*), meanwhile, captures the effect of that mutation on the expected number of offspring of class *i* produced by that initial class *j* parent. Finally, each offspring is weighted by its reproductive value 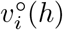, which gives its asymptotic contribution to the future of the population.

Plugging eq. (C21) into (C24) and substituting the result into eq. (C25), we obtain after some straight-forward re-arrangements that the selection gradient on dominance is proportional to

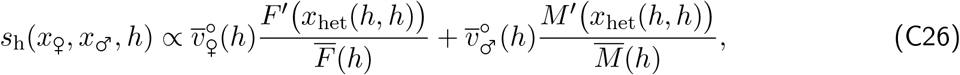

where *F* ^*′*^ (*x*_het_(*h, h*)) and *M* ^*′*^ (*x*_het_(*h, h*)) give the effect of a change in *h* on the fecundity of a *x*_♀_*/x*_♂_ heterozygote through female and male function, respectively;

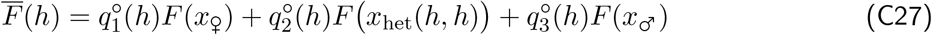

and,

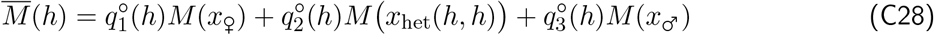

respectively are the average number of female and male gametes produced in the population, which measure the intensity of competition for reproduction through each sex; and finally

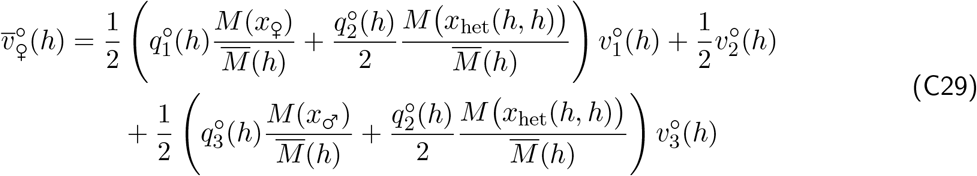

and,

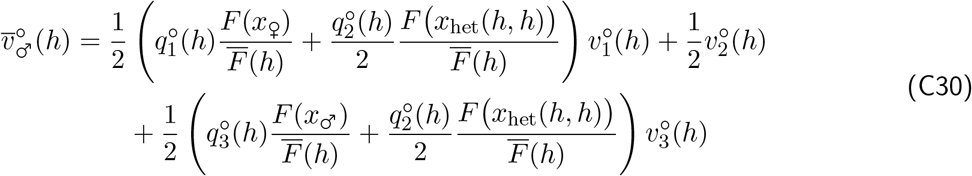

are the average reproductive values of descendants produced by a heterozygote through female and male function respectively. To see this, consider that 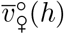 can be expanded into

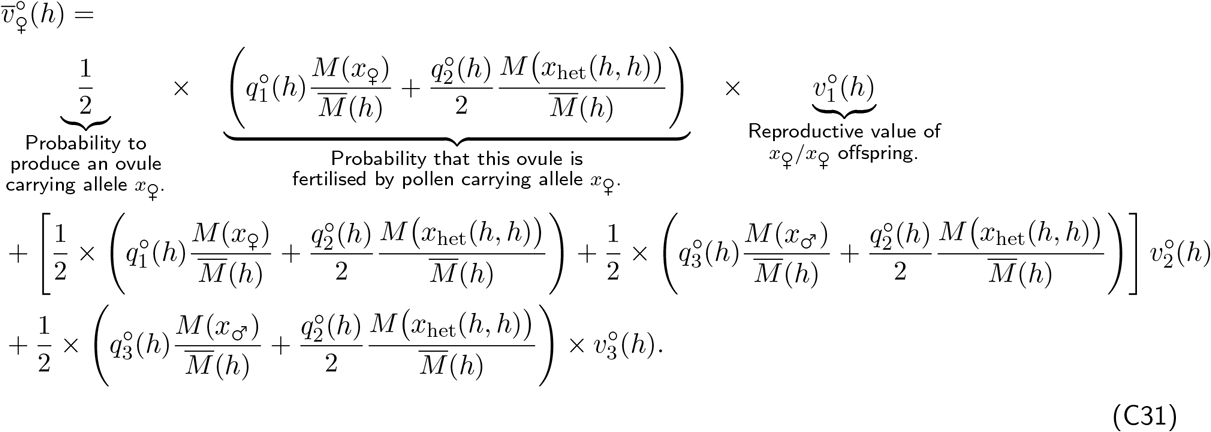

The underbraced term reveal how the first line can be read as the probability that a *x*_♀_*/x*_♂_ heterozygote produces a seed with genotype *x*_♀_*/x*_♀_, multiplied to the reproductive value (as defined in the **Class reproductive values** paragraph above) of such a seed. Likewise, the second and third lines of eq. (C31) correspond to the probability of producing a seed with genotypes *x*_♀_*/x*_♂_ and *x*_♂_*/x*_♂_, respectively, multiplied to the reproductive values of such seeds. Therefore, 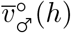 is the average reproductive value of descendants produced by a heterozygote *x*_♀_*/x*_♂_ through female function. Similarly, it can readily be shown that 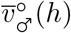 is the average reproductive value of descendants produced by a heterozygote *x*_♀_*/x*_♂_ through male function.

###### C.2.4.1 Competition between heterozygotes and with homozygotes determines selection on dominance

We first compute numerically the selection gradient on dominance (eq. C26) in a population where the resident allele at the dominance modifier locus codes for additivity (*h* = 1*/*2), i.e. we compute *s*_h_(*x*_♀_, *x*_♂_, 1*/*2) where 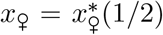 and 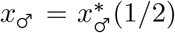. Results are shown in Fig. 4B of the main text. This shows that the sign of *s*_h_(*x*_♀_, *x*_♂_, 1*/*2) matches the outcome of our earlier simulations (detailed in Appendix C.1) nearly perfectly (compare Figs. 3D and 4B). Specifically, where *s*_h_(*x*_♀_, *x*_♂_, 1*/*2) > 0 so that selection favours an increase in dominance of the female allele when *h* = 1*/*2, simulated populations typically evolve a ZW system. Conversely, where *s*_h_(*x*_♀_, *x*_♂_, 1*/*2) < 0 so that selection favours an decrease in dominance of the female allele when *h* = 1*/*2, simulated populations typically evolve an XY system. To understand these results, we inspect the selection gradient on dominance eq. (C26) more closely below.

Eq. (C26) highlights that selection on dominance acts only in *x*_♀_*/x*_♂_ heterozygotes (the numerators in eq. C26), who compete with heterozygotes and homozygotes for reproduction (the denominators in eq. C26). The ratios

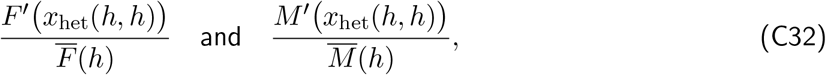

capture the effects of a small change in *h* in a *x*_♀_*/x*_♂_ heterozygote on its reproductive success through female and male function, respectively (i.e. the effect on the ‘quantity’ of offspring produced). These are weighted in eq. (C26) by the average reproductive value of offspring produced by a heterozygote through female and male function, 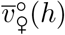 and 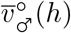 (as defined in eqs. (C29) and (C30), see also explanations and references given in paragraph **Class reproductive values** above). The two summands in eq. (C26) therefore quantify the fitness returns gained from a small change in *h* in a heterozygote through female and male function, respectively. How the shape of gain curves influence these fitness returns, however, is complicated because gain curves affect: (i) the fecundity gains achieved by heterozygotes through a small change in *h* (the numerators); (ii) the intensity of competition faced by heterozygotes (the denominators), both directly through the fecundities of resident homozygotes and heterozygotes (i.e. the resident *F* and *M*), and indirectly through the frequencies of these different genotypes in the population (i.e. the 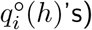; and (iii) the reproductive values 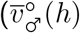 and 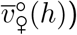.

To gain an intuitive understanding of selection on dominance, let us consider two scenarios for a *x*_♀_*/x*_♂_ heterozygote carrying a rare mutant dominance modifier. Consider first a scenario in which the resident population is composed exclusively of homozygotes at the sex allocation locus (so where 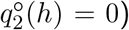. The *x*_♀_*/x*_♂_ heterozygote then faces intense competition from male (*x*_♂_*/x*_♂_) and female (*x*_♀_*/x*_♀_) homozygotes for reproduction through male and female function, respectively, as homozygotes always hold a competitive advantage over heterozygotes in their respective gamete pools. In this situation, selection favours a dominance modifier that makes the heterozygote allocate more to the sex in which competition is the weakest, i.e. dominance of the allele for the sex associated with the least increasing gain curve is favoured (Fig. S9A). Now consider the opposite scenario of a resident population composed exclusively of heterozygotes (so where 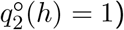). In this case, selection favours a dominance modifier that makes the heterozygote do better than resident heterozygotes, which is to allocate more to the sex where the fecundity benefits are the largest. Dominance of the allele for the sex associated with the most increasing gain curve is favoured (Fig. S9B).

The above argument illustrates how competition between heterozygotes and with homozygotes select for dominance to evolve in opposite directions. Whether selection favours XY (*h* → 0) or ZW sex determination (*h* → 1) for a given shape of gain curves (i.e. a given combination of *γ*_♀_ and *γ*_♂_) depends on the balance between these two opposing forces. When gain curves are strongly accelerating (*γ*_♀_ and *γ*_♂_ are large), homozygotes hold a substantial competitive edge over heterozygotes, so that heterozygotes contribute little to the overall competition for reproduction. As a result, competition imposed by homozygotes is the main factor driving the evolution of dominance here, favouring dominance of the allele for the sex with the least accelerating gain curve. As gain curves become more linear (*γ*_♀_ and *γ*_♂_ come closer to one), the competitive edge of homozygotes decreases, causing the contribution of heterozygotes to increase to the point where competition among heterozygotes becomes the main force determining the direction of selection on *h*, favouring dominance of the allele for the sex with the most accelerating gain curve.

###### C.2.4.2 A positive feedback leading to complete dominance

Once dominance has evolved away from additivity (i.e. *h≠* 1*/*2), heterozygotes become more similar to one homozygote than the other, reducing the competitive edge held by the homozygote they are more similar to and increasing the one held by the other homozygote. This asymmetry causes selection to intensify in the direction dominance has started to evolve towards, creating a positive feedback loop that eventually leads to complete dominance of one allele over the other (i.e. *h* = 0 or *h* = 1), as observed in all our simulations (see Fig. 3B-D in the main text).

To illustrate this, we can plot the selection gradient on dominance numerically for values of *h* ranging from 0 to 1 for a few representative cases and assuming power gain curves (eq. A4). We find that in every case, selection favours either *h* = 0 or *h* = 1 with a basin of attraction that varies depending on gain curve parameters (Fig. S10). In other words, there exists a threshold dominance coefficient *h*^*^ below which dominance of the male-biased allele is favoured (*h* → 0) and above which dominance of the female-biased allele is favoured (*h* → 1).

##### C.2.5 Effect of genetic drift

Equipped with the above results, we can now return to our simulations to explain why sometimes, the opposite outcome to the one favoured by selection evolves (Figs 3D and 3E in the main text). This is because selection on dominance, which is proportional to the divergence between sex allocation alleles, is weak when polymorphism first emerges at the sex allocation gene. Genetic drift can thus cause the population to cross the critical threshold *h*^*^ for selection on dominance (Fig. S10) before sex allocation alleles have sufficiently diverged, leading selection on dominance to switch direction. In scenarios where dioecy evolves (where *γ*_♀_ > 1 and *γ*_♂_ > 1), genetic drift can thus cause the population to acquire ZW instead of an XY sex determination (and vice-versa). Where the equilibrium population is gyno- or androdioecious (i.e., where only *γ*_♀_ > 1 or only *γ*_♂_ > 1), unisexual strategies can end up being encoded by a recessive rather than a dominant allele. In the gynodioecious case for instance, the pure female strategy can end up being encoded by a recessive allele, so that the population is composed of three genotypes instead of two at equilibrium: homozygous (XX) females, and heterozygous (XY) and homozygous (YY) hermaphrodites.

## Appendix D The effects of partial self-fertilisation and inbreeding depression

Here, we investigate the impact of partial selfing and inbreeding depression, and derive the results that are summarised in the main text section “Partial selfing and inbreeding depression favour XY sex determination”. First, we investigate how selfing influences the emergence of polymorphism in sex allocation in Appendix D.1. Second, we examine the effect of selfing on the gradual differentiation of alleles leading to dioecy or other sexual systems in Appendix D.2. And lastly in Appendix D.3, we look at how selfing affects the emergence of XY and ZW sex determination systems through dominance evolution.

### D.1 Evolutionary dynamics of sex allocation under partial selfing

We first study the evolutionary dynamics of sex allocation under partial selfing. Like in Appendix B, we assume that sex allocation *x* is genetically encoded by alleles with additive effects at a quantitative trait locus. We label alleles at this locus by their quantitative phenotypic effects, so that a carrier of alleles *x*_1_ ∈ [0, 1] and *x*_2_ ∈ [0, 1] expresses a sex allocation strategy *x* = (*x*_1_ + *x*_2_)*/*2. Mutations arise at a small and constant rate, and have weak, unbiased phenotypic effects.

#### D.1.1 Invasion fitness

We consider a rare mutant *x*_mut_ arising in a population fixed for a resident allele *x*. Due to partial selfing, the mutant is now able to mate with itself even when it is rare, so that it can exist both in heterozygous and homozygous form. In this case, the dynamics of the sub-population of mutants is captured by the matrix equation

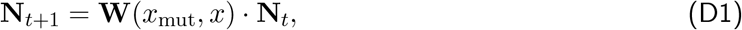

where **N**_*t*_ = (*n*_1,*t*_, *n*_2,*t*_) is a vector containing the number of heterozygous and homozygous mutants in the population at time *t* (i.e. the number of individuals that carry one and two copies of *x*_mut_, respectively) and **W**(*x*_mut_, *x*) is a 2 *×* 2 matrix whose (*i, j*)-entry *w*_*ij*_(*x*_mut_, *x*) corresponds to the number of successful mutant offspring of type *i* produced by a mutant of type *j* (where type 1 are *x*_mut_*/x* heterozygous and type 2 *x*_mut_*/x*_mut_ homozygous mutants). The elements of the **W**(*x*_mut_, *x*) matrix are given by

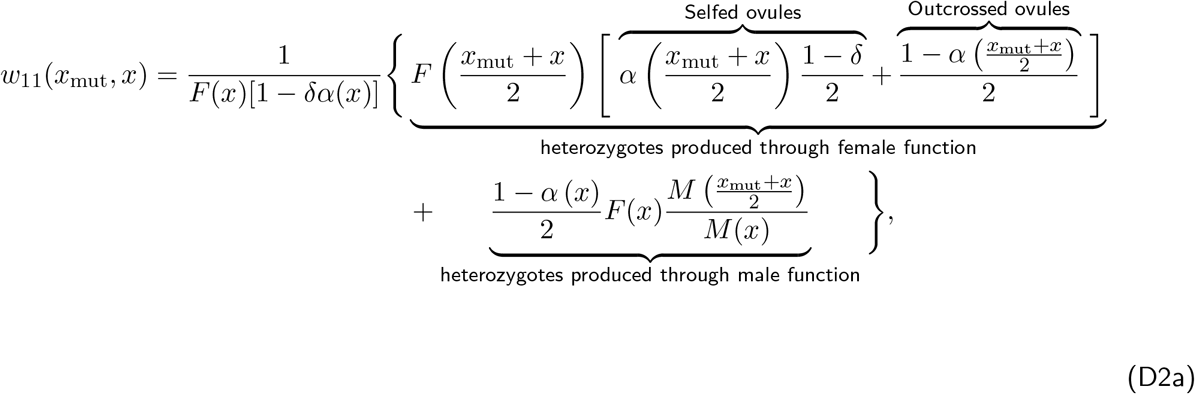

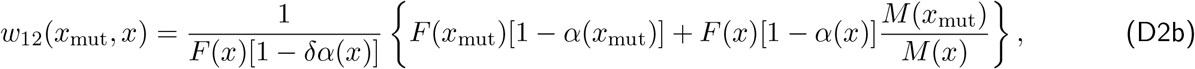

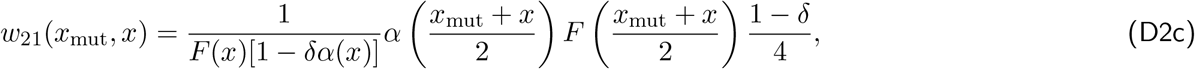

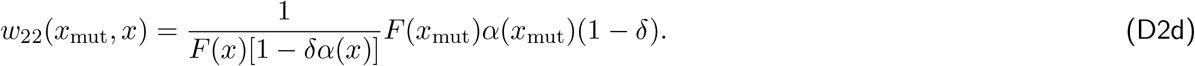

Let us consider *w*_11_(*x*_mut_, *x*) (eq. D2a) as an example of how these entries were derived. This is the number of heterozygous mutants produced by heterozygous mutants. Heterozygous mutant offspring can be produced: through female function via selfing and outcrossing, and through male function by siring the ovules of resident individuals (see braced labels in that equation). In each case, heterozygous mutants have a probability 1*/*2 of transmitting a single copy of the mutant allele *x*_mut_ to their offspring, resulting in the production of heterozygous mutants. Other entries were derived similarly.

The invasion fitness of the mutant allele *x*_mut_ is given by the leading eigenvalue *ρ*(*x*_mut_, *x*) of the matrix **W**(*x*_mut_, *x*). This eigenvalue can be obtained directly but is unsightly and thus of little help for interpretation. We thus refrain from giving its full expression here (but see the accompanying *Mathematica* notebook).

#### D.1.2 Directional selection on sex allocation under partial selfing

##### D.1.2.1 Selection gradient

We first examine how selfing influences directional selection on sex allocation through the selection gradient. The results we obtain in this subsection are largely already described in Charlesworth and Charlesworth (1981), which considers directional selection on sex allocation at the phenotypic level (or, equivalently, assumes individuals are haploid). Our analysis below shows that Charlesworth and Charlesworth (1981)’s results extend to diploidy and additive gene action. The main reason we go over these results here is that they are useful for understanding disruptive selection in section D.1.3, i.e. for understanding how selfing influences the emergence of polymorphism via evolutionary branching (which is not considered in Charlesworth and Charlesworth, 1981).

The selection gradient on sex allocation is obtained from the derivative with respect to *x*_mut_ of the leading eigenvalue *ρ*(*x*_mut_, *x*) of the matrix **W**(*x*_mut_, *x*) given by eq. (D2), which we find can be written as,

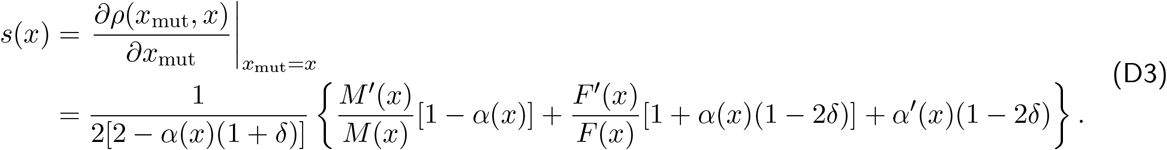

This expression is almost identical to the selection gradient obtained by Charlesworth and Charlesworth (1981) (eq. 2a on page 60 of their paper). The main difference is that Charlesworth and Charlesworth (1981) expressed female fecundity as a function of male fecundity, i.e. they wrote male and female fecundities as *b* and *f* (*b*) instead of *M* (*x*) and *F* (*x*), respectively, and computed the gradient on *b* instead of *x* (denoted *r* in their model).

To consider the effects of selfing and inbreeding depression, it is useful to rearrange the selection gradient (eq. D3) as,

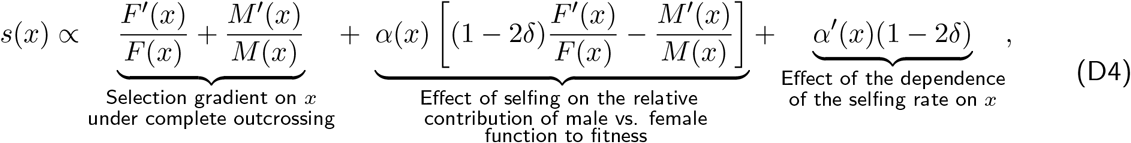

where the first term corresponds to the selection gradient on sex allocation under complete outcrossing (eq. B3), and the second and third terms capture the effects of selfing and inbreeding depression. Let us first consider the case where the selfing rate is independent of sex allocation (i.e. *β* = 0 in eq. A1) so that *α*(*x*) = *α*_0_, *α*^*′*^(*x*) = 0 and the third term in eq. (D4) vanishes. Solving for the singular strategy *x*^*^ using eq. (D4), we find that *x*^*^ must be such that

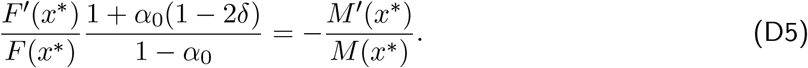

Since 1+*α*_0_(1−2*δ*) > 1−*α*_0_, comparing eq. (D5) with eq. (B5) indicates that when selfing is independent of sex allocation (i.e. *α*^*′*^(*x*) = 0), selfing always favours more female-biased sex allocation strategies relative to the complete outcrossing case (Charlesworth and Charlesworth, 1981). This is because male function only yields fitness returns via outcrossed ovules, whereas female function yields fitness returns through both selfed and outcrossed ovules, so that selfing increases the relative contribution of female function to fitness. Inbreeding depression *δ* decreases but never completely offsets this female bias (unless *δ* = 1).

When selfing decreases with allocation to female function (i.e. *β >* 0 in eq. A1 so that *α*^*′*^(*x*) < 0), the third term in eq. (D4) (which is now non-zero) reveals that there is now either added selection to increase allocation to female function when inbreeding depression is high (*δ >* 1*/*2), or opposite selection to increase allocation to male function when inbreeding depression is low (*δ <* 1*/*2; Charlesworth and Charlesworth, 1981). This is because there is selection to reduce selfing when inbreeding depression is high, and thus to invest into female function (owing to eq. A1). In contrast, when *δ <* 1*/*2, there is selection to increase selfing and thus to invest into male function as selfed offspring provide higher fitness returns than outcrossed offspring due to transmission advantage (Fisher, 1941). As a result, although partial selfing generally tends to favour allocating more resources to female function compared to the outcrossing case, it may be possible that when inbreeding depression *δ* is low, investing into male function is in fact selected for. This possibility, which is dismissed in Charlesworth and Charlesworth (1981), is explored in the next section.

##### D.1.2.2 Selfing can favour maleness

To investigate whether selfing can favour maleness, let us denote the singular sex allocation strategy favoured under complete outcrossing by 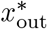 (i.e. satisfying eq. B5). If the selection gradient eq. (D4) is negative at 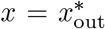, then selection favours a more male-biased strategy under partial selfing than under complete outcrossing. After substitution and simplification, we have

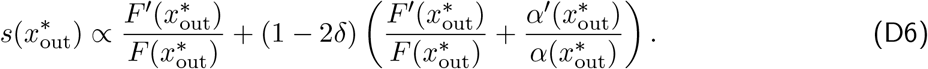

This reveals that where *δ <* 1*/*2, 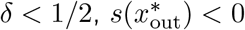only if

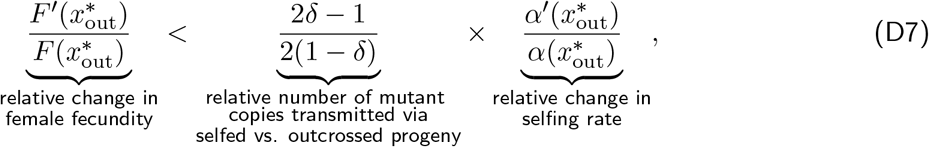

where the first factor in the right-hand side of eq. (D7) is the relative difference between the average number of mutant gene copies passed onto the next generation by a mutant parent via selfed and outcrossed seeds. To see this, let us denote by *m*_out_(*q*) and *m*_self_ (*q*), the average number of mutant copies carried by outcrossed (*m*_out_(*q*)) and selfed (*m*_self_ (*q*)) zygotes produced by a randomly sampled mutant parent, when the mutant sub-population is composed of heterozygotes *x/x*_mut_ and homozygotes *x*_mut_*/x*_mut_ in frequencies *q* and 1 − *q*. These are given by

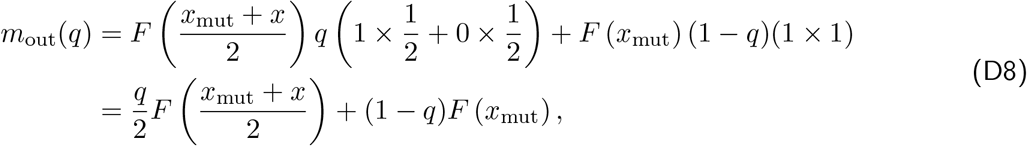

and,

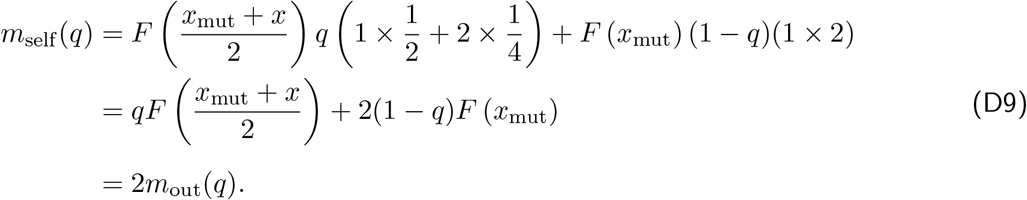

To understand these equations, consider the first line of eq. (D9) as an example. The first term gives the average number of mutant copies transmitted by a mutant heterozygote to selfed zygotes: it transmits one copy with probability 1*/*2, and two copies with probability 1*/*4 due to Mendelian segregation (and zero with probability 1*/*4). The second term gives the average number of mutant copies transmitted by a mutant homozygote via selfing, which is necessarily equal to two because homozygotes only carry mutant alleles. The twofold transmission advantage of selfing is made clear by the fact that *m*_self_ (*q*) = 2*m*_out_(*q*), i.e. the average number of copies transmitted to zygotes via selfing is always twice the average number of copies transmitted through outcrossing, irrespective of the frequencies of heterozygous and homozygous parents and of the phenotypic effect of the mutant (Fisher, 1941).

From eqs. (D8) and (D9), we can then obtain the average number of mutant copies contributed to the next generation (i.e. to the pool of viable seeds). For a randomly sampled mutant via outcrossing, this is 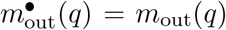, because all outcrossed zygotes survive to become viable seeds. To obtain the average number of mutant copies contributed to the next generation through selfing (denoted 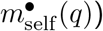, meanwhile, we must account for the fact that a self-fertilised zygote survives with probability 1 − *δ* to become a seed due to inbreeding depression. Thus, we have

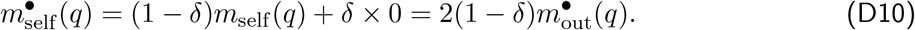

The relative difference between the average number of mutant gene copies passed onto the next generation by a mutant parent via selfed and outcrossed seed is thus given by

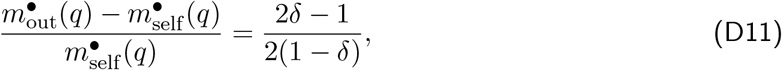

which corresponds to the factor on the right-hand side in eq. (D7).

The product on the right-hand side of eq. (D7) therefore quantifies the fitness cost of a decrease in selfing rate, whereas the left-hand side captures the fitness benefit of an increase in female fecundity. If becoming more female (increasing *x*) yields a fecundity benefit that is lower than the cost of producing lesser quality offspring (i.e. offspring carrying less mutant copies on average, that is more outcrossed offspring in our case because *δ <* 1*/*2), then selection will favour a more male-biased sex allocation under partial selfing than under complete outcrossing. The conditions for this to happen are restrictive, which may explain how they can go unnoticed in a numerical scan (as done in Charlesworth and Charlesworth, 1981). But these conditions may still be met. With power gain curves (eq. A4), for instance, eq. (D7) requires that *(i)* the male gain curve is saturating (*γ*_♂_ < 1*/*2), *(ii)* inbreeding depression is low (*δ <* (1 − 2*γ*_♂_)*/*2(1 − *γ*_♂_)); and *(iii)* the selfing rate decreases significantly with allocation to female function, i.e. such that

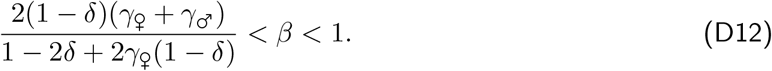

##### D.1.2.3 Singular strategy under power gain curves

Assuming power gain curves (eq. A4), the selection gradient on sex allocation becomes

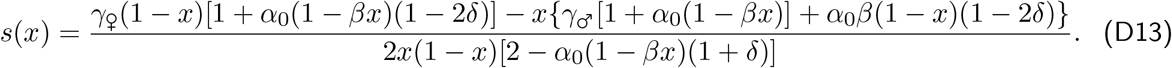

Solving for the singular sex allocation strategy *x*^*^ (as defined in eq. B4 and using eq. D13), we obtain

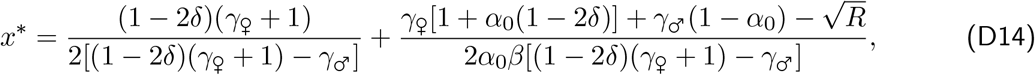

with

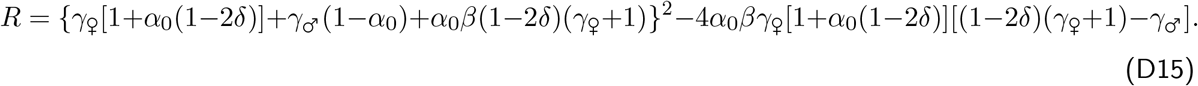

It can readily be shown that this singular strategy is always convergence stable.

#### D.1.3 Disruptive selection

Once the population expresses the singular strategy *x*^*^, it may either experience stabilising selection and remain monomorphic for *x*^*^, or disruptive selection and become polymorphic. Which of these two outcomes unfolds is determined by the sign of the disruptive selection coefficient,

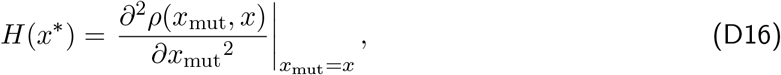

which we show after some re-arrangements to be proportional to

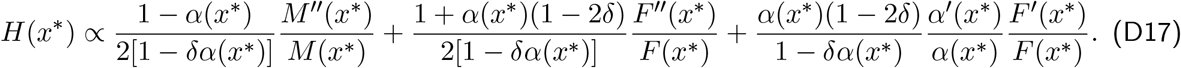

The first two terms capture the same effect of the shape of gain curves as under complete outcrossing (eq. B11): accelerating gain curves favour the emergence of polymorphism. However, the effects via male and female gain curves in eq. (D17) are each weighted by the relative contribution of each sexual function to fitness. To see these relative contributions, consider the number of mutant gene copies transmitted to the next generation by a randomly sampled mutant in a resident population monomorphic for *x*^*^. Using eqs. D8-D10, we have that under neutrality and at the singular strategy (i.e. where *x*_mut_ = *x* = *x*^*^), the average number of mutant copies transmitted by a mutant parent through selfed and outcrossed progeny are respectively given by

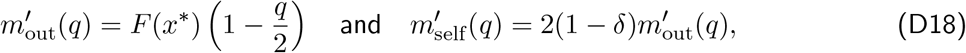

when the mutant sub-population is composed of heterozygotes and homozygotes in frequencies *q* and 1 − *q*. Thus, the average number of mutant copies transmitted through female function, 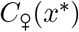, is

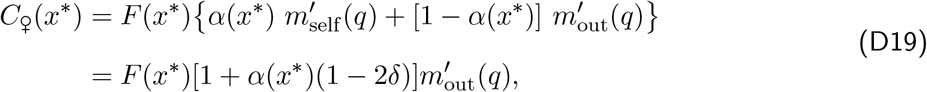

whereas the average number of copies transmitted through male function is

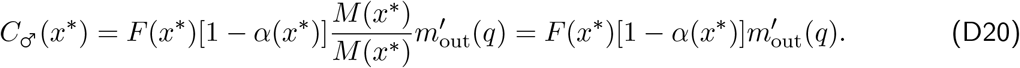

Therefore,

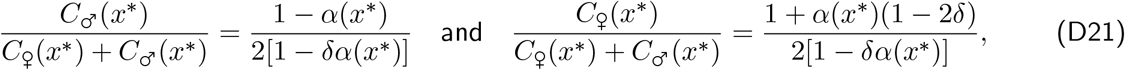

correspond to the relative contribution of male and female function to the fitness of a mutant individual. These tune the relative influence of the shape of the male and female gain curves on disruptive selection, as shown in the first two terms of eq. (D17)). The third term in eq. (D17) emerges due to the effect of sex allocation on the selfing rate (i.e. due to *α*^*′*^(*x*^*^) < 0). Since by definition *F* ^*′*^(*x*^*^) > 0, this term reveals that when inbreeding depression is high (*δ >* 1*/*2), polymorphism tends to be promoted, i.e. that individuals that become more female and others that become more male are both favoured by selection. This is because when inbreeding depression is high, one should avoid producing offspring through selfing and instead increase the rate of outcrossing, and there are two ways of achieving this: either increase allocation to female function to reduce one’s selfing rate (owing to eq. A1), or increase allocation to male function to increase outcrossing through pollination of others’ ovules. This pathway to dioecy can be seen as the classical inbreeding avoidance pathway (Charlesworth and Charlesworth, 1978a,b, 1981), here taken by gradual evolution, that is by small mutational steps.

### D.2 The emergence of dioecy, gyno- and androdioecy

We now examine how selfing influences the gradual divergence of sex allocation alleles under disruptive selection, in particular, whether selection eventually leads to alleles coding for dioecy, gyno- or androdioecy. We follow the same approach as in Appendix B.2.2 and thus consider a rare mutant *x*_mut_ that arises in a population where two alleles *x*_1_ and *x*_2_ already coexist.

#### D.2.1 Equilibrium of the resident population

We first characterise the demographic equilibrium of the resident population where alleles *x*_1_ and *x*_2_ coexist. The demographic state at generation *t* consists in the numbers of individuals carrying genotypes *x*_1_*/x*_1_, *x*_1_*/x*_2_ and *x*_2_*/x*_2_, which we denote as *n*_11,*t*_, *n*_12,*t*_ and *n*_22,*t*_, respectively. We write

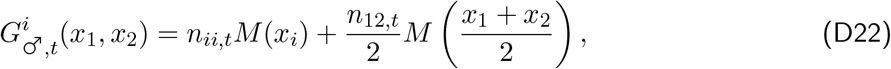

for the amount of pollen carrying allele *x*_*i*_ (*i* ∈ {1, 2}) produced in the population, and

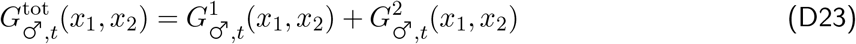

for the total amount of pollen produced in the population. Similarly, let

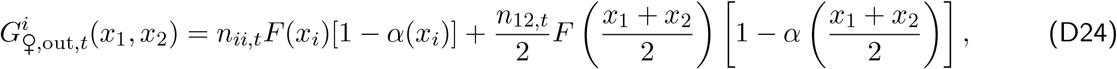

denote the number of ovules to be outcrossed carrying allele *x*_*i*_ in the population, and

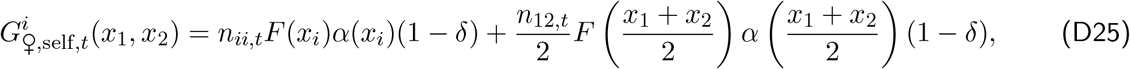

denote the number of ovules destined for selfing carrying allele *x*_*i*_, so that the total number of ovules produced by the whole population is given by

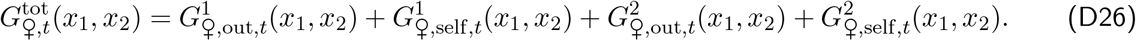

In what follows, we refer to the variables defined above in eqs. (D22)-(D26) as ‘*G*-variables’ for short.

Using these, the numbers of individuals of the three genotypes in generation *t* + 1 are given by

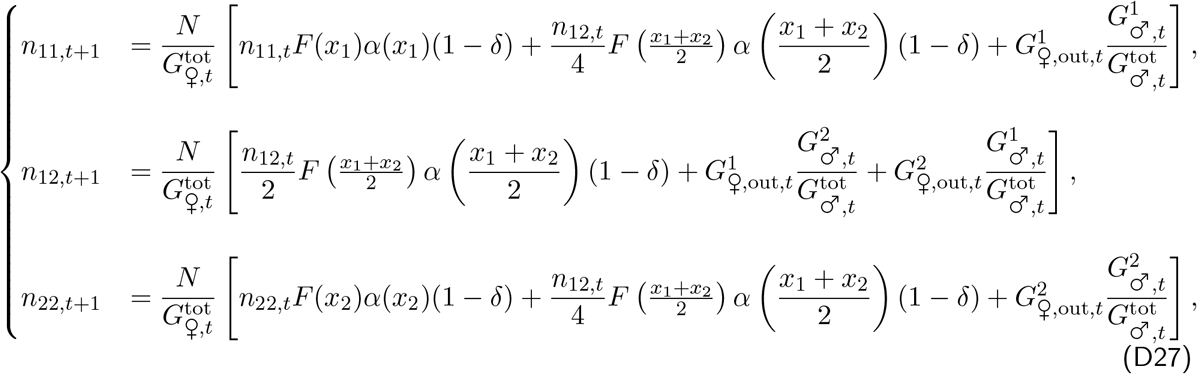

where the (*x*_1_, *x*_2_) arguments of *G*-variables were dropped for brevity. The number of individuals of each genotype at demographic equilibrium 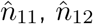 and 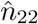, are then determined by

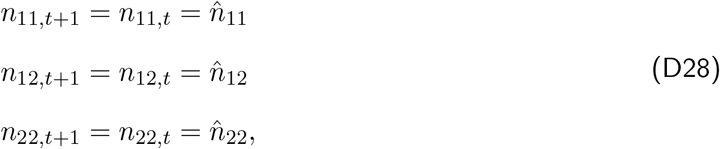

Where 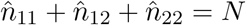. We do not solve for this equilibrium explicitly but use the above characterisation in upcoming numerical analyses.

#### D.2.2 Invasion analysis

Because individuals can self-fertilise, a rare mutant allele *x*_mut_ can be found in three different forms in the population (in contrast to Appendix B.2.2): two heterozygous forms (*x*_1_*/x*_mut_ and *x*_2_*/x*_mut_) and one homozygous form (*x*_mut_*/x*_mut_). These three forms are referred to as classes 1, 2 and 3 hereafter, respectively. The dynamics of the mutant population are modelled by the matrix equation

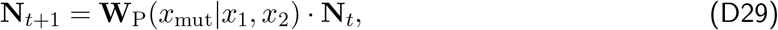

where **N**_*t*_ = {*N*_1,*t*_, *N*_2,*t*_, *N*_3,*t*_} is a vector containing the number of mutant individuals in each class at time *t*, and **W**_P_(*x*_mut_|*x*_1_, *x*_2_) is a 3 *×* 3 matrix whose (*i, j*)−entry 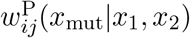 is the expected number of successful mutants of class *i* produced by a mutant of class *j*.

To specify the elements of **W**_P_(*x*_mut_|*x*_1_, *x*_2_), we introduce 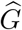-variables 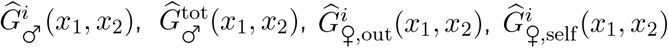 and 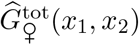 with *i* ∈ {1, 2}, which correspond to the *G*-variables defined above (eqs. D22-D26), expressed at the resident population’s demographic equilibrium (so where eq. D28 holds). Using these variables and dropping their arguments for brevity, we have

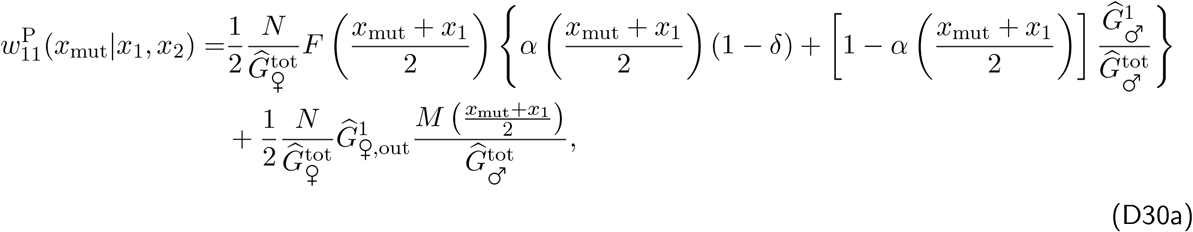

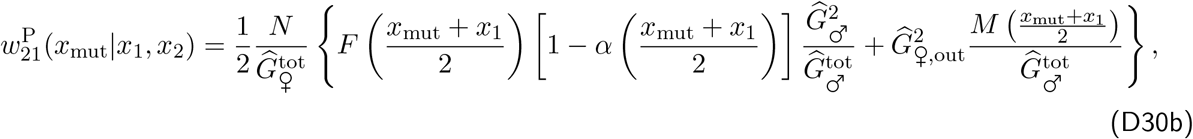

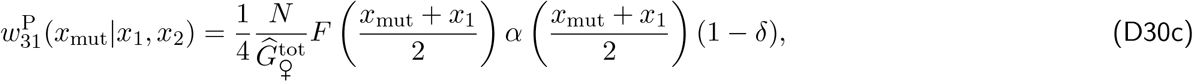

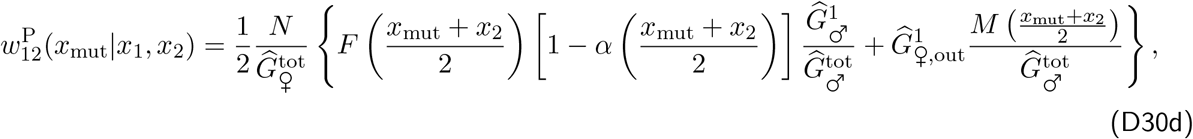

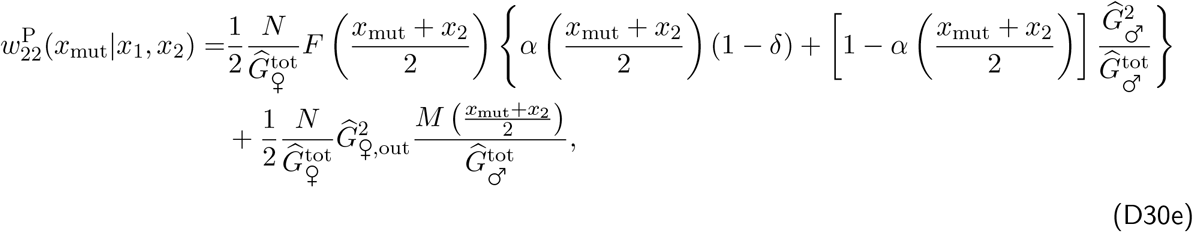

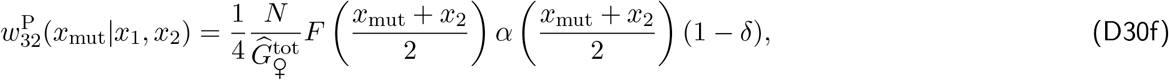

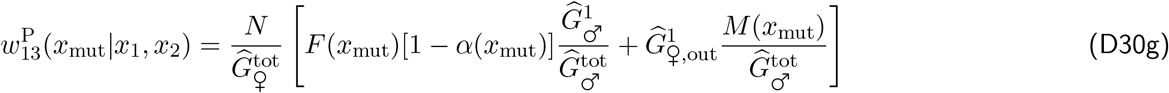

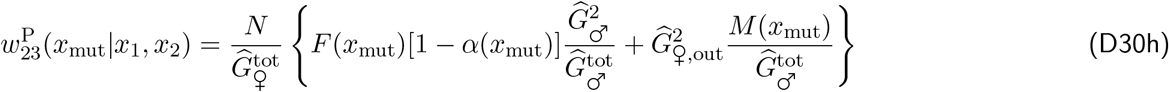

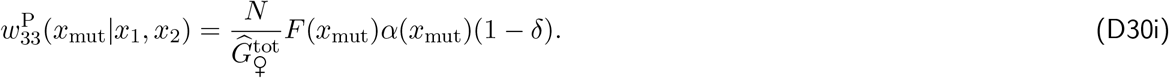

The invasion fitness of mutant allele *x*_mut_ is the leading eigenvalue *ρ*^P^(*x*_mut_|*x*_1_, *x*_2_) of the matrix **W**_P_(*x*_mut_|*x*_1_, *x*_2_). The selection gradient acting on alleles *x*_1_ and *x*_2_ are given by

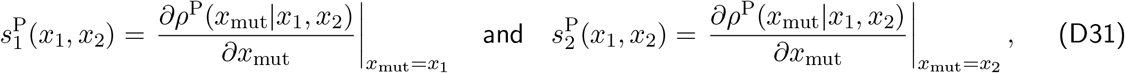

which can be used to infer on the evolutionary dynamics of the polymorphic population (as described in Appendix B.2.2 below eq. B27). We do so numerically in the next section.

#### D.2.3 Numerical analysis

We study the polymorphic equilibrium reached by the population numerically under low (*δ* = 0.25) and high (*δ* = 0.75) inbreeding depression and low (*α*_0_ = 0.25) and high (*α*_0_ = 0.75) baseline selfing rates, keeping *β* = 1 fixed (in eq. A1). Our approach largely follows the one described in Appendix B.2.3. For each combination of selfing rate *α*_0_ and levels of inbreeding depression *δ*, we compute the selection gradients 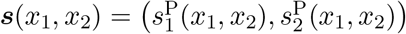 for many pairs (*x*_1_, *x*_2_) across phenotypic space to obtain a vector field that determines the evolutionary trajectories favoured by selection. Inspection of these vector fields reveals five possible evolutionary outcomes: cases (i)-(iv) are as those that are shown in Fig. S4, corresponding to monomorphic hermaphroditism (so that *x*_1_ = *x*_2_), gyno- and androdioecy and dioecy; case (v) corresponds to a dimorphic hermaphroditism whereby two differentiated alleles coexist, each coding for an intermediate sex allocation i.e. (0 *< x*_1_ *< x*_2_ < 1, e.g. Fig. S11).

Fig. S12 shows where cases (i)-(v) hold in the space of gain curves. As can be seen from this figure, dimorphic hermaphroditism tends to occur when both sexes have saturating gain curves and inbreeding depression is high (Fig. S12C-D, purple region). More generally, comparing Fig. S12 with Fig. S3 shows that in contrast to the outcrossing case, polymorphic sexual system can emerge even where both sexes have saturating gain curves, as long as selfing is sufficiently common and inbreeding depression is sufficiently high. Selection here is driven by inbreeding avoidance. But otherwise, selfing does not dramatically alter evolutionary outcomes. When inbreeding depression is low (*δ* = 0.25, Fig. S12A-B), polymorphism and in particular dioecy (red region) and androdioecy (yellow region) tend to be disfavoured by selfing. This is because of the diminishing fitness returns via female function generated by selfing in the absence of inbreeding depression. Conversely, because fitness returns via female function increase due to selfing when inbreeding depression is high (*δ* = 0.75, Fig. S12C-D), this tends to favour gynodioecy (green region).

### D.3 Evolution of XY and ZW sex determination under partial selfing

Finally, we compute the selection gradient on dominance in a population that is polymorphic at the sex allocation locus to investigate the impact of partial selfing on the evolution of XY and ZW sex determination. We extend the model of dominance evolution described in Appendix C.2 to include partial selfing and inbreeding depression. As a reminder, we assume that two sex allocation alleles *x*_♀_ and *x*_♂_ (*x*_♀_ *> x*_♂_) segregate in the population. In heterozygotes *x*_♀_*/x*_♂_, allele *x*_♀_ expresses in proportion to a dominance coefficient *h*. We assume *h* to be a quantitative trait influenced by an unlinked modifier locus at which alleles are additive. We label alleles at the modifier by their quantitative effect on dominance, so that an individual carrying alleles *h*_*i*_ and *h*_*j*_ ∈ [0, 1] at the modifier expresses a dominant coefficient

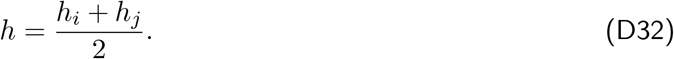

We denote by *x*_het_(*h*_*i*_, *h*_*j*_) the sex allocation strategy expressed by a heterozygote carrying these alleles at the modifier. This strategy is given by

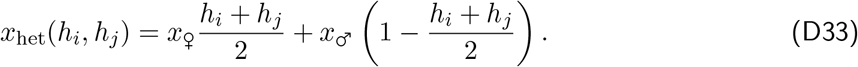

We study the evolution of *h* by considering the fate of a rare mutant allele *h*_mut_ in a resident population otherwise fixed for *h*.

#### D.3.1 Invasion analysis

We begin by characterising the equilibrium state reached by a resident population fixed with *h* at the dominance modifier, and where alleles *x*_♀_ and *x*_♂_ coexist at the sex allocation locus. This is equivalent to the analysis described in Append D.2.1, except that here dominance in the resident population is arbitrary (rather than *h* = 1*/*2). The state of the population at a given generation *t* is characterised by the number of individuals with genotypes *x*_♀_*/x*_♀_, *x*_♀_*/x*_♂_ and *x*_♂_*/x*_♂_ in the population, which we denote as 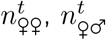 and 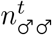, respectively. To characterise the change in the number of individuals of each genotype between two generations, we denote as

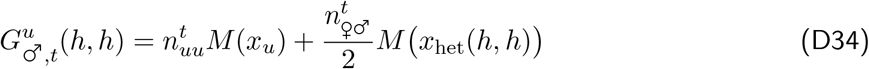

the number of male gametes carrying allele *x*_*u*_ (*u* ∈ {_♀_, _♂_}) produced by the resident population, and

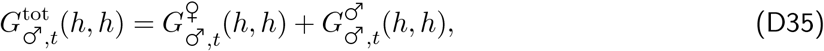

the total number male gametes produced by the resident population. Furthermore, we denote as

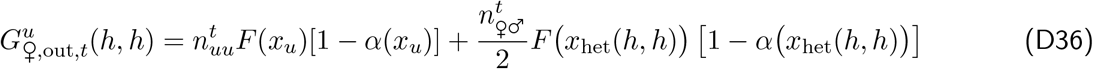

the number of outcrossed seeds carrying allele *x*_*u*_ (*u* ∈ {_♀_, _♂_}) produced by the resident population, and

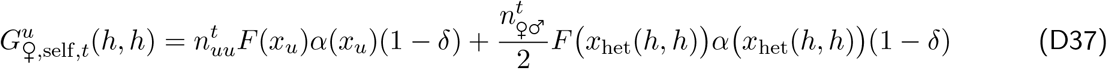

the number of selfed seeds carrying allele *x*_*u*_ (*u* ∈ {_♀_, _♂_}) produced by the resident population, so that the total number of seeds carrying allele *x*_*u*_ is given by

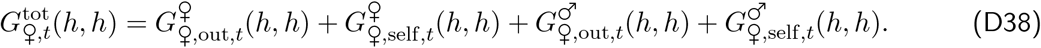

The variables defined above are collectively referred to as *G*-variables hereafter. Using these variables, we have

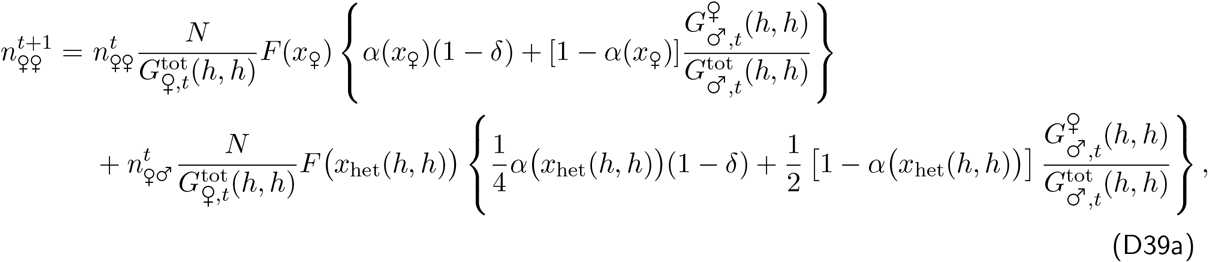

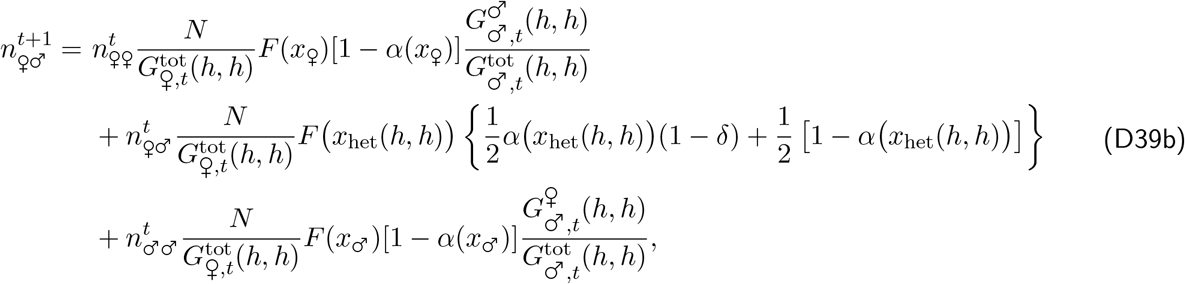

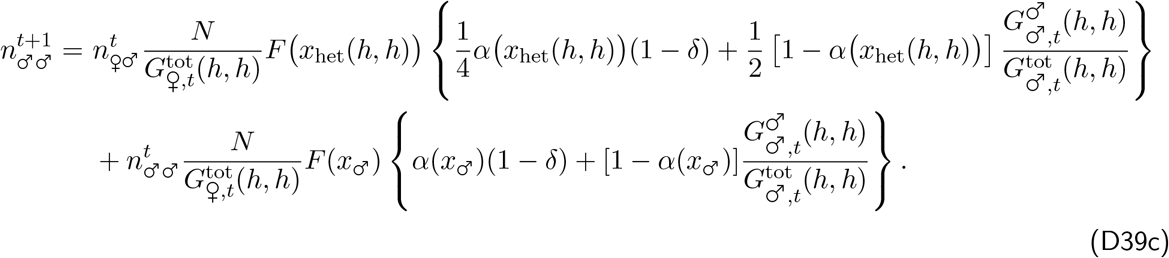

These recursions can then be used to obtain equilibrium number of individuals of each genotype,

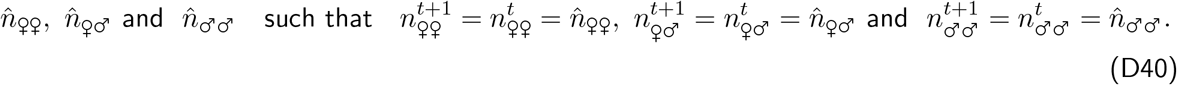

Next, we introduce a mutant *h*_mut_ allele at the dominance modifier locus, when the resident population is otherwise fixed for *h* at this locus but polymorphic at the sex allocation locus (with genotypes *x*_♀_*/x*_♀_, *x*_♀_*/x*_♂_ and *x*_♂_*/x*_♂_ present). Because of partial selfing, mutants at the dominance modifier can be either heterozygous (*h*_mut_*/h*, as before) or homozygous (*h*_mut_*/h*_mut_). The mutant population can therefore be divided among six classes: 2 genotypes at the dominance modifier *×* 3 genotypes at the sex allocation locus. We label these from 1 to 6: heterozygotes *h*_mut_*/h* at the dominance modifier with genotype *x*_♀_*/x*_♀_, *x*_♀_*/x*_♂_ and *x*_♂_*/x*_♂_ at the sex allocation locus are labeled as 1, 2 and 3, respectively, and homozygotes *h*_mut_*/h*_mut_ with genotype *x*_♀_*/x*_♀_, *x*_♀_*/x*_♂_ and *x*_♂_*/x*_♂_ are labeled as 4, 5 and 6, respectively. The dynamics of the mutant population is modelled by a matrix equation

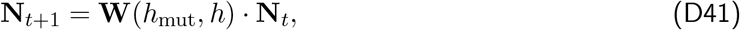

where **N**_*t*_ = (*n*_*i,t*_)_1⩽*i*⩽6_ is a vector containing the number of individuals in each class, and **W**(*h*_mut_, *h*) is a 6 *×* 6 matrix whose (*i, j*)-entry *w*_*ij*_(*h*_mut_, *h*) gives the number of successful mutants of type *i* produced by a focal mutant of type *j*. The **W**(*h*_mut_, *h*) matrix is very large and is thus not shown here (see accompanying *Mathematica* notebook).

#### D.3.2 Effect of selfing and inbreeding depression on selection on dominance

From standard theory on selection in class-structured population (eq. A2 in Taylor and Frank (1996), see also Caswell, 2001; Taylor, 1990; Avila and Mullon, 2023), the selection gradient on dominance *s*_h_(*x*_♀_, *x*_♂_, *h*) can then be computed

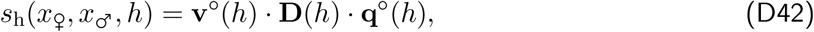

where **q**^*°*^(*h*) is the vector of asymptotic class frequencies under neutrality, which is given by the right eigenvector of **W**^*°*^(*h*) = **W**(*h, h*), normalised such that

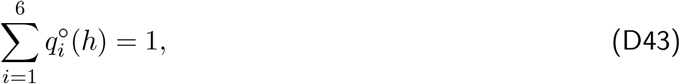

**D**(*h*) is the matrix of first derivatives of **W**(*h*_mut_, *h*), whose (*i, j*)-entry *d*_*ij*_(*h*) is given by

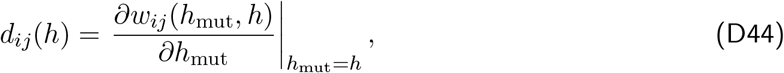

and **v**^*°*^(*h*) is the vector of class-specific reproductive values under neutrality, which is given by the left eigenvector of **W**^*°*^(*h*) normalised such that

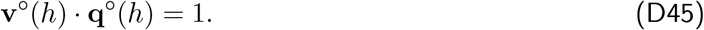

Recall from section C.2.4 that reproductive values capture the relative influence of individuals from each class on the long-term demography of the population (e.g. Fisher, 1930; Charlesworth, 1980; Caswell, 2001; Rousset, 2004). These reproductive values are difficult to characterise analytically but they can be straightforwardly computed numerically. One useful property here is that because homozygous carry twice as many copies of the mutant allele but are identical to the heterozygotes in every other aspect under neutrality (so when *h*_mut_ = *h*), the reproductive value of homozygous offspring is exactly twice that of the heterozygous i.e.,

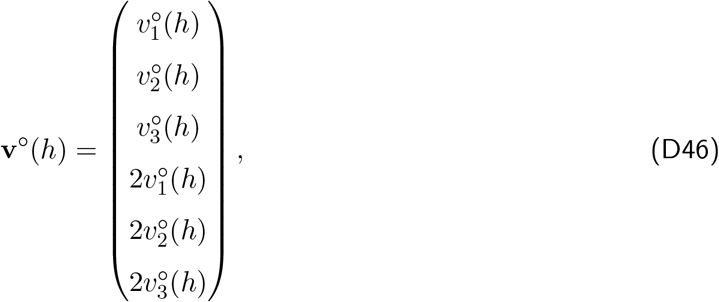

Where 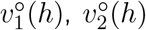 and 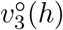 are the reproductive values of individuals that are heterozygote for the mutant allele (*h*_mut_*/h*). Similarly, although we could not characterise analytically the asymptotic class frequencies (the **q**^*°*^(*h*) vector), these frequencies are, by definition, related to one another as,

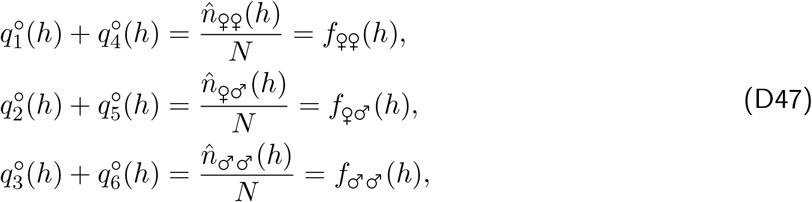

where *f*_♀♀_(*h*), *f*_♀♂_(*h*) and *f*_♂♂_(*h*) denote the frequencies of genotypes *x*_♀_*/x*_♀_, *x*_♀_*/x*_♂_ and *x*_♂_*/x*_♂_ in the resident population (with *f*_♀♀_(*h*) + *f*_♀♂_(*h*) + *f*_♂♂_(*h*) =1).

Plugging eqs. (D46) to (D47) together with the **W**(*h*_mut_, *h*) matrix given in the accompanying *Mathematica* notebook into eq. (D42), we find that the selection gradient on dominance is proportional to

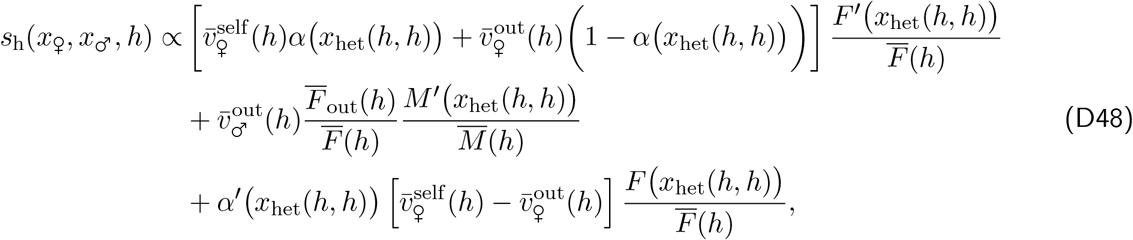

where

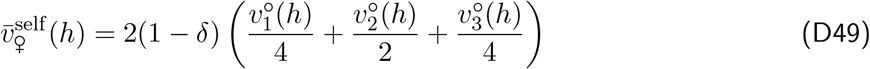

is the mean reproductive value of offspring produced by a *x*_♀_*/x*_♂_ heterozygote via self-fertilisation;

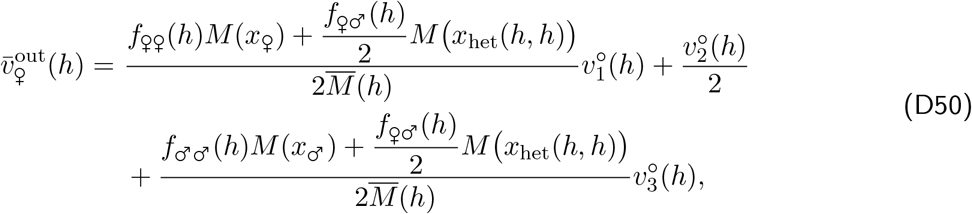

and,

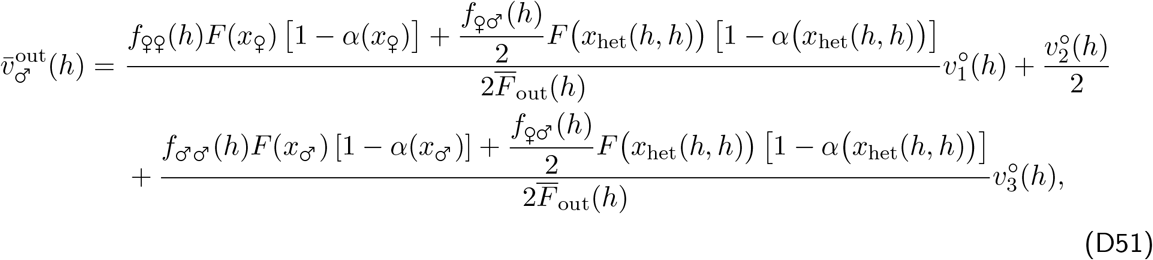

are the mean reproductive value of offspring produced by heterozygotes via outcrossing through female and male function, respectively;

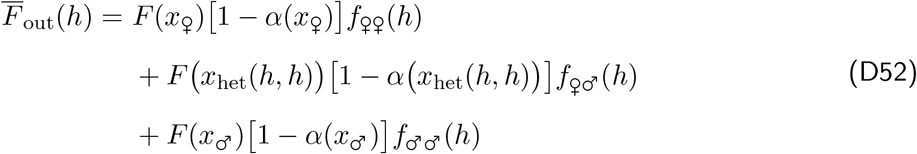

is the average number of ovules available for outcrossing produced by a resident individual; and

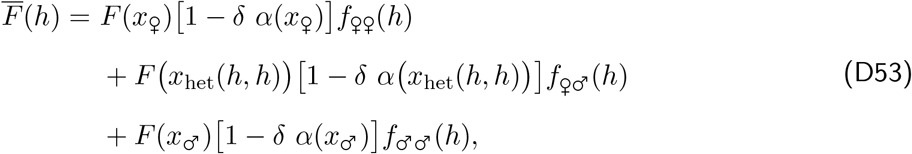

and,

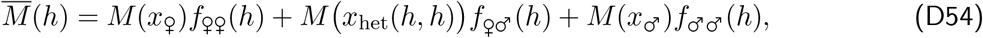

are average number of viable seeds and pollen grains produced by a resident individual.

The first two lines of eq. (D48) correspond to the fitness effect of a change in dominance in a *x*_♀_*/x*_♂_ heterozygote through female and male function, respectively. Comparing these two lines with the complete outcrossing case (eq. C26) highlights how selfing influences fitness gained through female and male function functions differently depending on the level of inbreeding depression. The third line eq. (D48) corresponds to the fitness effect of a change in dominance in a *x*_♀_*/x*_♂_ heterozygote through its effect on the selfing rate.

To understand better how selfing and inbreeding depression influence the emergence of sex determining systems, we computed the selection gradient on dominance (eq. D48) in a population where the resident allele at the dominance modifier locus codes for additivity (*h* = 1*/*2), i.e. we computed *s*_h_(*x*_♀_, *x*_♂_, 1*/*2) where *x*_♀_ and *x*_♂_ are the equilibria of evolutionary dynamics when *h* = 1*/*2 (see Appendix D.2.3). As in the outcrossing case, we expect that ZW-systems are favoured when *s*_h_(*x*_♀_, *x*_♂_, 1*/*2) > 0, and XY-systems are favoured when *s*_h_(*x*_♀_, *x*_♂_, 1*/*2) < 0. Results of this analysis are presented in Fig. S13.

Comparing Fig. S13 with the outcrossing case shown in Fig. 4B reveals that selfing typically favours the evolution of XY systems, especially when inbreeding depression is high (Fig. S13D). This is due to two effects of selfing, for which the decomposition eq. (D48) is useful to understand. First, selfing increases the frequency of *x*_♀_*/x*_♀_ relative to *x*_♂_*/x*_♂_ homozygotes. This increases competition through female function (i.e. makes 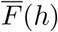 larger) but reduces competition through male function (i.e. makes 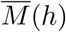 smaller). A *x*_♀_*/x*_♂_ heterozygote thus has an advantage to becoming more male, favouring the evolution of XY sex determination. The second effect that selfing has when inbreeding depression *δ* is high, is to decrease the mean reproductive value of offspring produced through female function (i.e. the term in square brackets on the first line of eq. D48 becomes small). This is because selfed offspring have a lower reproductive value than outcrossed offspring when *δ* is large (i.e.,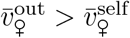). This reduces any potential benefits that a *x*_♀_*/x*_♂_ would have by increasing female function, thus facilitating the evolution of XY.

## Appendix E A multilocus simulation model

Here, we detail the simulation model used to generate Fig. 5 of the main text, where sex allocation is initially a polygenic trait whose basis evolves. Our model is built on those of van Doorn and Dieckmann (2006) and Kopp and Hermisson (2006) and is illustrated in Fig. S14. The basic structure of the simulation follows the one described in Appendix C.1 (we assume complete outcrossing here for simplicity), except that the sex allocation strategy expressed by an individual is now determined by *L* unlinked loci. Each locus consists of a promoter sequence and a gene affecting sex allocation. The promoter sequence controls dominance relationships between alleles segregating at the gene, as before (Appendix C.1), so that the sex allocation strategy encoded by the *k*^th^ locus in individual *i, x*_*i,k*_, is given by

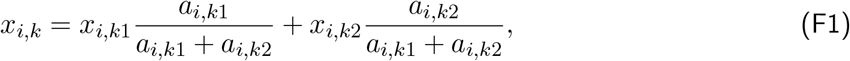

where *a*_*i,k*1_ and *a*_*i,k*2_ ∈ (0, +∞) are the promoter affinities of alleles at the *k*^th^ locus and *x*_*i,k*1_ and *x*_*i,k*2_ ∈ [0, 1] are the sex allocation strategies encoded by alleles at the *k*^th^ locus.

We additionally consider evolution at an unlinked modifier locus that determines the contribution made by each sex allocation locus to the phenotype (in light blue in Fig. S14). The effect of an allele at the modifier locus is given by a vector of size *L*, such that we write the *n*^th^ allele carried by individual *i* at the modifier as

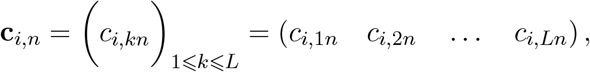

such that *c*_*i,kn*_ ∈ (0, +∞) modulates the relative contribution of locus *k*. Alleles at the modifier are assumed to be additive, so that the relative contribution *C*_*i,k*_ of the *k*^th^ locus to the phenotype of the *i*^th^ individual is given by the sum of its contribution values at the modifier divided by the sum of the contributions of all loci, i.e.

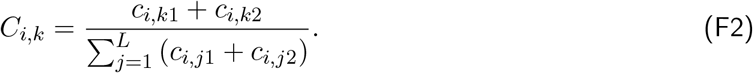

Overall, the sex allocation strategy *x*_*i*_ expressed by individual *i* is given by the sum of the strategies encoded by the *L* loci, weighted by their relative contributions, i.e.,

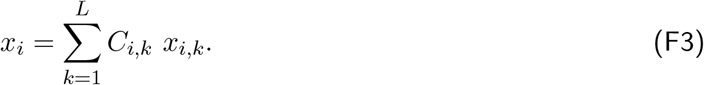

Each time an offspring is produced, each of its alleles mutates independently with probability *µ*. When a promoter (e.g. *a*_*i,k*1_) or gene (e.g. *x*_*i,k*1_) allele mutates, its new value is sampled in a Gaussian distribution centered on the parental value, with standard deviation *σ*, ensuring their effect is kept within bounds (i.e. greater than zero for promoter affinities, and between 0 and 1 for sex allocation strategies). When an allele (e.g. **c**_*i,n*_) mutates at the modifier, all the *L* contributions it encodes independently change to new values sampled in a Gaussian distribution centered on their respective parental values, with standard deviation *σ* (and truncated to be kept within bounds if necessary i.e. above zero).

The population is initially fixed at all sex allocation loci for some arbitrary strategy *x*_0_ ∈ (0, 1) and affinity *a*_0_ = 1 at the promoter, and fixed for equal contributions at the modifier (i.e. all contributions are fixed to *c*_0_ = 1). Figure 5 in the main text was obtained using this simulation program. It illustrates that disruptive selection leads to the concentration of the genetic architecture of sex allocation into a single sex-determining locus, via the silencing of all others (i.e. all the contributions *c*_*i,kn*_ except one go to zero).

## Appendix F Fruit dispersal and the shape of the female gain curve

Our results have revealed how the shape of the male and female gain curves can affect the evolution of sexual systems. Little is known about the factors that affect gain curve shape, and especially about those that may lead to accelerating gain curves, but a few hypotheses have been suggested (Janzen, 1971; Lloyd, 1982; Givnish, 1982; Charlesworth, 1999). In plants with fleshy fruits, in particular, individuals allocating more resources to their female function may have more efficient seed dispersal if seed dispersers are more attracted to plants producing larger crops. This in turn may lead to reduced kin competition among their offspring, yielding an accelerating female gain curve (Givnish, 1982). This argument has been analysed theoretically, and the coupling of seed dispersal ability and seed production has indeed been shown to promote dioecy (Vamosi et al., 2007; Biernaskie, 2010). In this appendix, we give a simple mathematical formalisation of this argument using the notations and methods of our model.

### F.1 The model

We consider a population of *N* individuals, where each occupies one of *N* homogeneous breeding spots, with the following life cycle. *(i) Sexual development:* First, individuals allocate resources to their female and male functions in proportions *x* and 1 − *x*, respectively, resulting in the female and male fecundities *F* (*x*) = *F*_0_*x* and *M* (*x*) = *M*_0_(1 − *x*). Fecundity is assumed to increase linearly with resource allocation to highlight how relevant non-linearity can emerge from other ecological factors (this is equivalent to setting *γ*_♀_ = *γ*_♂_ = 1 in eq. A4). *(ii) Mating:* Individuals export all their pollen, so that no self-fertilisation occurs, distributing it equally among the *N* − 1 other individuals in the population. The pollen grains received by an individual compete to fertilise its ovules, so that eventually all ovules are fertilised (there is no pollen limitation). The diploid zygotes formed through syngamy are assumed to immediately undergo division to give rise to two haploid seeds, so that resulting individuals are haploid. We make this assumption because it simplifies mathematical analysis. *(iii) Dispersal:* Individuals disperse a fraction *d*(*x*) = *x* of their seeds to other patches in the population, where the dispersal rate *d*(*x*) depends on their sex allocation strategy, and a fraction 1 − *d*(*x*) remain in their natal patch. Here, the seed dispersal rate is assumed to increase with allocation to female function, i.e. seed dispersal is coupled with seed production. This assumption captures the idea that individuals producing larger crops of fruits can attract more dispersers, and thus enjoy more efficient dispersal of their seeds, as previously proposed (Givnish, 1982). Dispersed seeds are equally likely to fall on each of the *N* − 1 non-natal patches in the population (as in Wright’s island model, Wright, 1931). Seed dispersal is assumed to increase linearly with seed production for simplicity. *(iv) Density-regulation:* All adults die, and the seeds present on a patch compete to occupy it and grow into an adult.

### F.2 Invasion analysis

The sex allocation strategy *x* is encoded by a quantitative trait locus undergoing recurrent small effect mutations (‘continuum of alleles’ model). We study the evolution of *x* by considering the fate of a rare mutant *x*_mut_ arising in a resident population monomorphic for *x*. The invasion fitness of the mutant *W* (*x*_mut_, *x*) can be decomposed into its male and female components, *w*_♂_(*x*_mut_, *x*) and *w*_♀_(*x*_mut_, *x*).

#### Male fitness component

The male component corresponds to the fraction of seeds sired by the mutant that inherit the mutant allele, which simplifies to

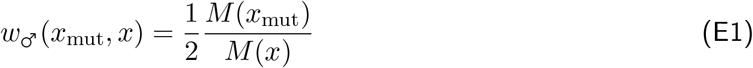

after accounting for density-dependence and assuming *N* is large. The male component of fitness depends linearly on sex allocation *x* because male fecundity *M* (*x*) is assumed to be linear in *x* (i.e. the male gain curve is linear).

#### Female fitness component

The female component corresponds to the sum of mutant seeds competing for recruitment locally and globally,

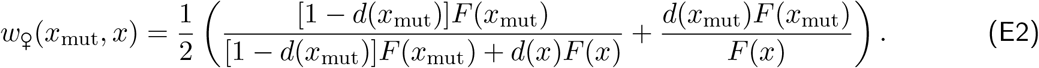

Contrary to the male component of fitness, the female component depends non-linearly on sex allocation despite female fecundity *F* (*x*) being a linear function of *x*, as a result of limited seed dispersal and the coupling of seed dispersal and seed production. Limited seed dispersal causes related seeds to compete with one another for recruitment, which generates kin competition. This leads to diminishing fitness returns in the female function (i.e. a saturating female gain curve), because kin competition intensifies as the number of competing seeds increases. The coupling of seed dispersal and seed production allows individuals to partially avert the effect of kin competition on their progeny as they increase seed production, which generates increasing fitness returns in the female function (i.e. an accelerating female gain curve). Thus, the non-linearity of the female gain curve results from ecological interactions in this model.

#### Invasion fitness

Using eqs. (E1) and (E2), the invasion fitness of the mutant is given by

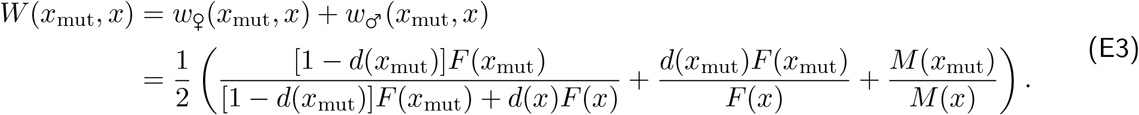

##### F.2.1 Directional selection

As before, the selection gradient is

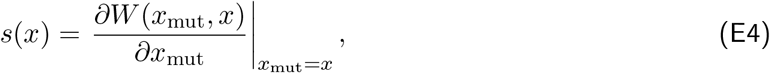

which using eq. (E3) yields

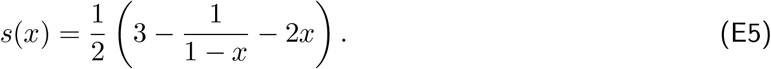

Solving *s*(*x*^*^) = 0 for *x*^*^ gives the singular strategy

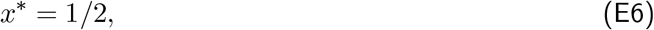

which satisfies

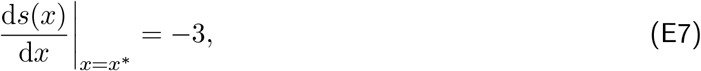

and is therefore convergence stable. Thus, directional selection leads an initially monomorphic population to express sex allocation strategy *x*^*^ = 1*/*2, where all individuals allocate equally to the male and female functions.

### F.2.2 Disruptive selection

Once the population expresses this strategy, it may either experience stabilising selection and remain monomorphic, thereby maintaining hermaphroditism, or disruptive selection and become polymorphic. Which of these two outcomes unfolds depends on

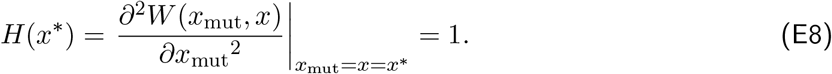

The fact that *H*(*x*^*^) is positive indicates that once the population has converged to *x*^*^ = 1*/*2, it experiences disruptive selection and becomes polymorphic. Taking the second derivative of the male and female components of the mutant’s fitness (eqs. E1 and E2) with respect to *x*_mut_, and evaluating them at *x*_mut_ = *x* = *x*^*^, we have

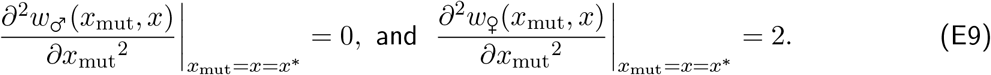

This shows that male fitness varies linearly with *x* around the singular strategy, whereas female fitness is accelerating around *x*^*^. Thus, the coupling of seed dispersal with seed production generates an accelerating female gain curve, which favours the emergence of polymorphism. Further, since the male curve is least accelerating, we expect the evolution of an XY system if sex allocation was expressed at the diploid stage.

---

**Figure S1:**
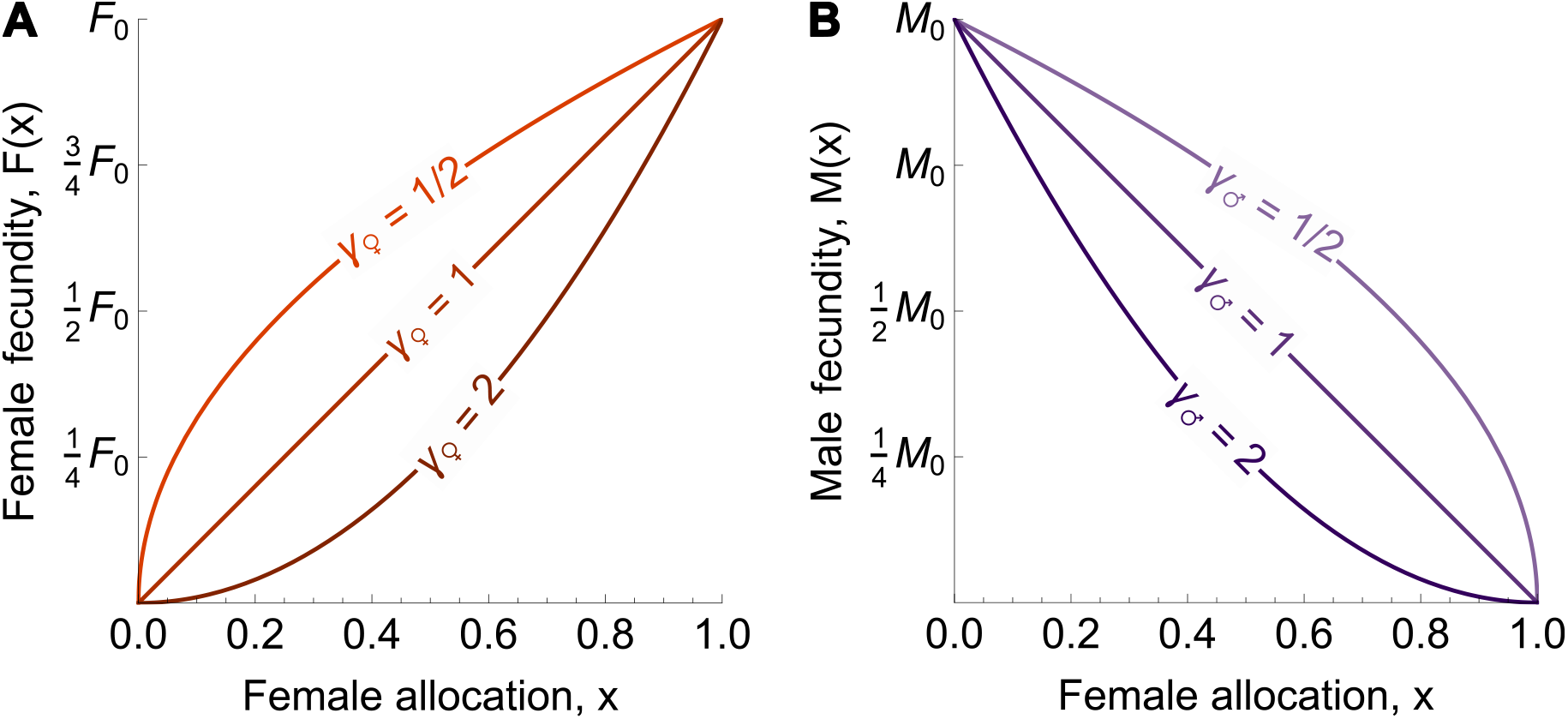
Male and female gain curves. **A** Female gain curve *F* (*x*) (eq. A4) as a function of female allocation *x* when it is saturating (*γ*_♀_ < 1), linear (*γ*_♀_ = 1) and accelerating (*γ*_♀_ > 1). **B** Shape of the male gain curve *M* (*x*) (eq. A4) as a function of female allocation *x* when it is saturating (*γ*_♂_ < 1), linear (*γ*_♂_ = 1) and accelerating (*γ*_♂_ > 1) with respect to male allocation (1 − *x*).

**Figure S2:**
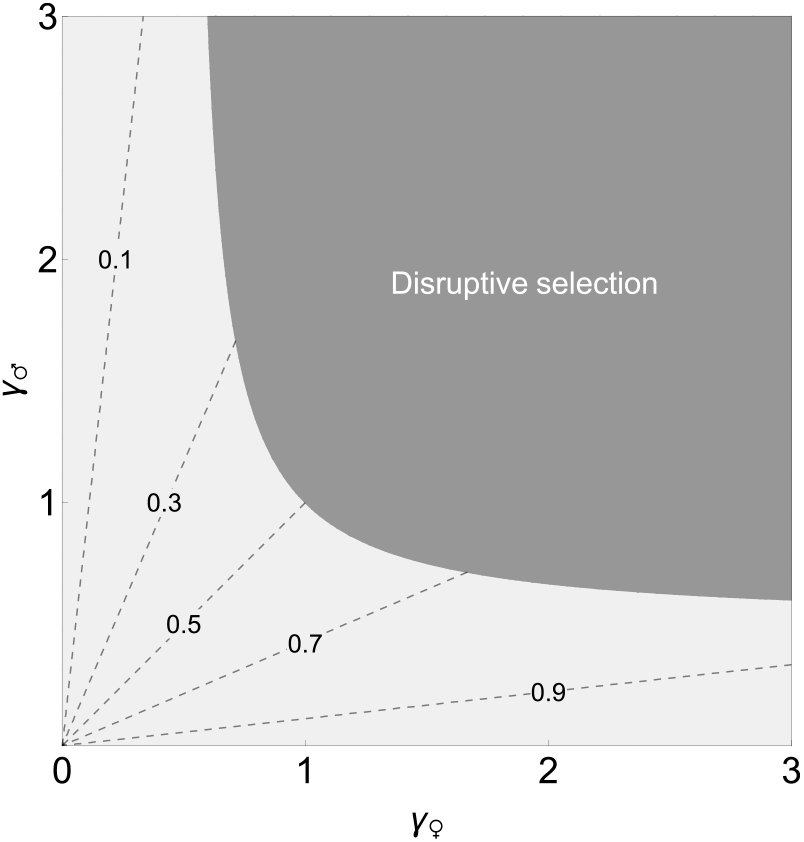
Condition for the emergence of polymorphism as a function of the male and female gain curve exponents. Condition (B13) is satisfied in the dark grey area, which results in disruptive selection and the emergence of polymorphism. The light grey area corresponds to the parameter space in which condition (B13) is not satisfied, so that selection is stabilising at *x*^*^ and the population remains monomorphic. The dashed lines in this region indicate the evolutionary stable sex allocation strategy *x*^*^ reached by the population (eq. B6).

**Figure S3:**
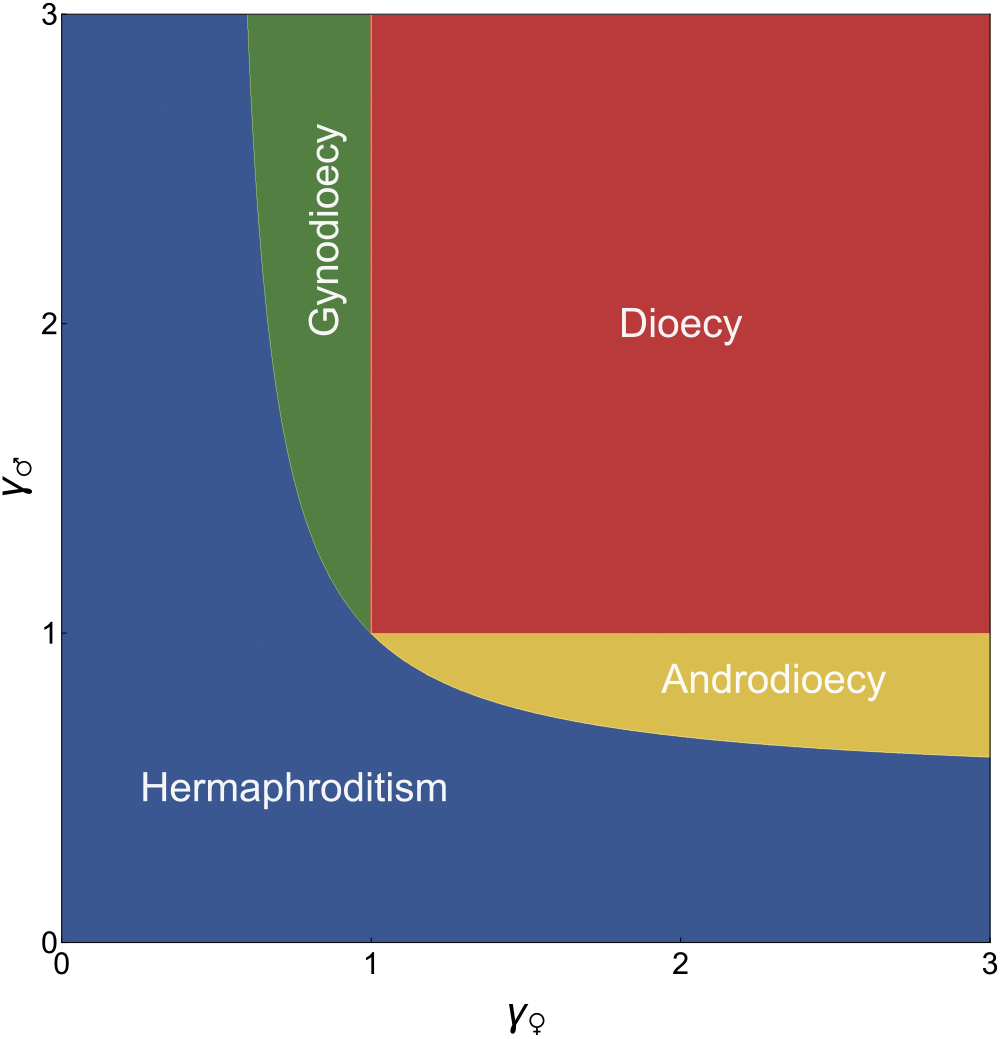
Parameter space corresponding to the four possible outcomes of gradual evolution: hermaphroditism (blue), androdioecy (yellow), gynodioecy (green) and dioecy (red).

**Figure S4:**
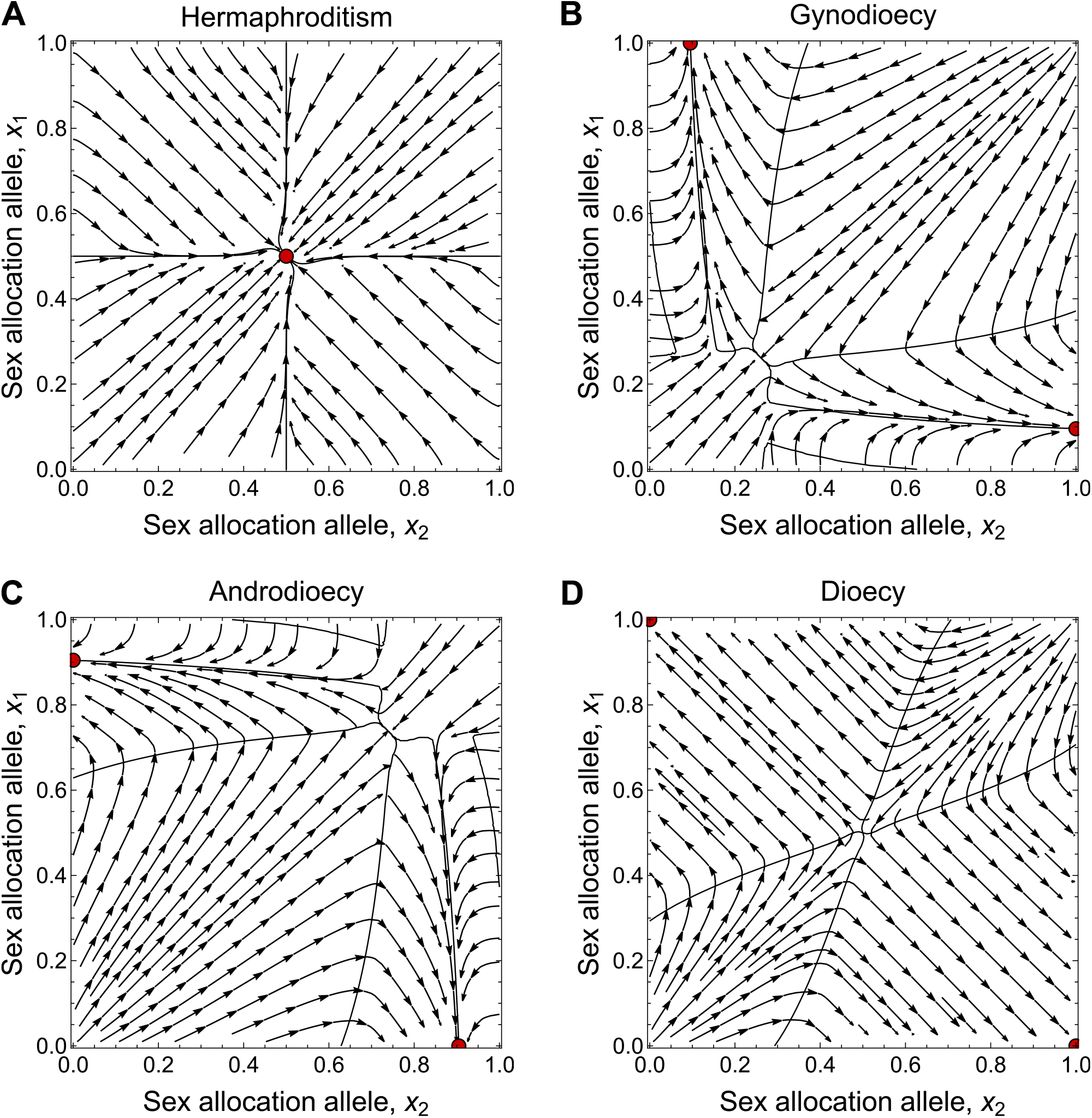
Streamplots illustrating the evolutionary dynamics of the population in the four regions presented in Fig. S3. Arrows indicate the direction in which the population evolves at a given point, and red dots indicate stable states. Solid black lines are isoclines (i.e., lines along which 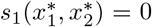 and 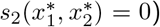). **A** Hermaphroditism 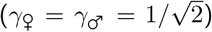: selection pushes the population to be monomorphic for the singular sex allocation strategy *x*^*^, i.e. *x*_1_ = *x*_2_ = *x*^*^ (*x*^*^ = 1*/*2 in this example, eq. B6). **B** Gynodioecy 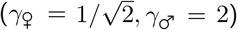: selection favours the evolution and maintenance of a pure female allele (*x* = 1) and a malebiased hermaphroditic allele (*x* = 0.1) with two symmetrical equilibria at 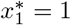 and 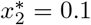, or 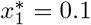 and 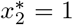. **C** Androdioecy 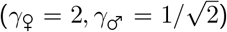: selection favours the evolution and maintenance of a pure male allele (*x* = 0) and a female-biased hermaphroditic allele (*x* = 0.9) with two symmetrical equilibria, similar to gynodioecy. **D** Dioecy (*γ*_♀_ = 2, *γ*_♂_ = 2): selection favours the evolution and maintenance of a pure female allele (*x* = 1) and a pure male allele (*x* = 0). All plots were obtained by computing the selection gradient on strategies *x*_1_ and *x*_2_ at many points in the phenotype space (eq. B27) and interpolating them. The interpolated selection gradients were then used to obtain the isoclines by solving *s*_1_(*x*_1_, *x*_2_) = 0 and *s*_2_(*x*_1_, *x*_2_) = 0 numerically.

**Figure S5:**
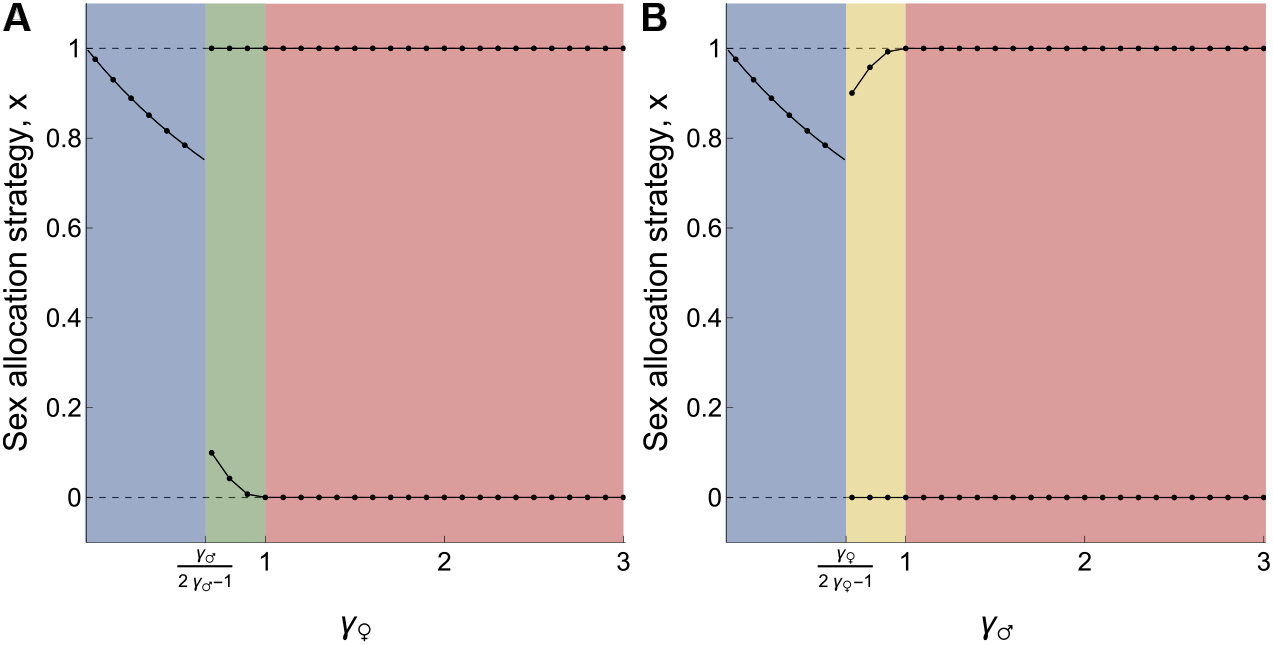
Equilibrium sex allocation strategies encoded by alleles, computed by applying the method described above to eq. (B27). **A** *γ*_♂_ = 2 and *γ*_♀_ varies from 0 to 3. **B** *γ*_♀_ = 2 and *γ*_♂_ varies from 0 to 3. Background colours corre-spond to regions in Figure S3.

**Figure S6:**
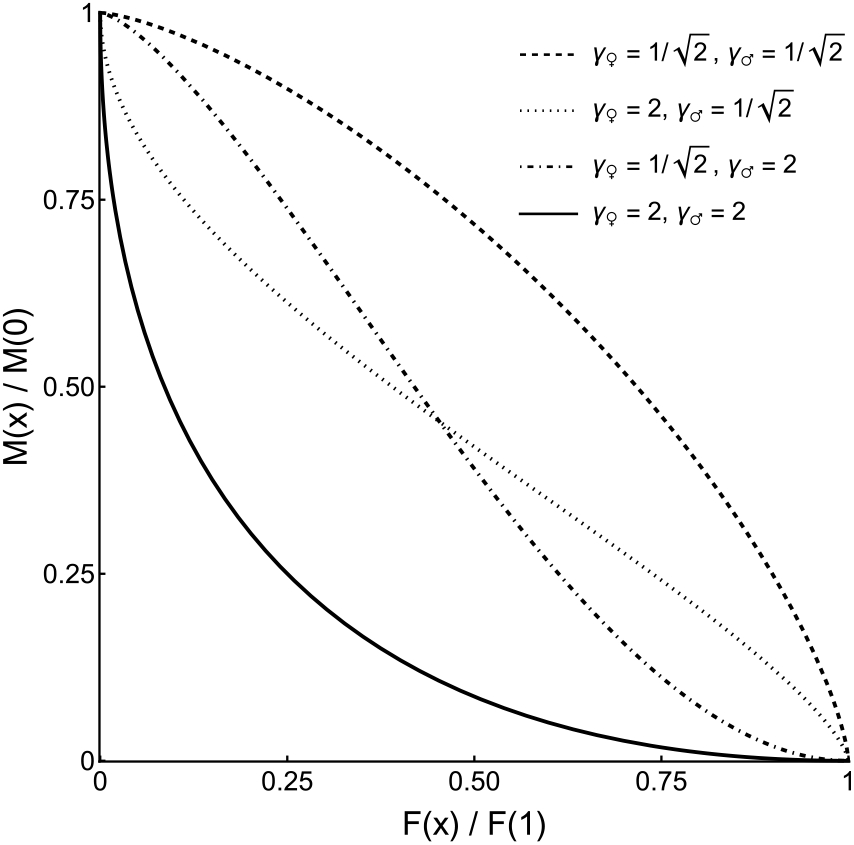
Fitness set *φ* for different shapes of gain curves. The dashed line shows a convex fitness set which favours the maintenance hermaphroditism. The dotted and dot-dashed lines show convex-concave and concave-convex fitness sets, which favour andro- and gynodioecy, respectively. The solid line shows a concave fitness set, which favours the evolution of dioecy.

**Figure S7:**
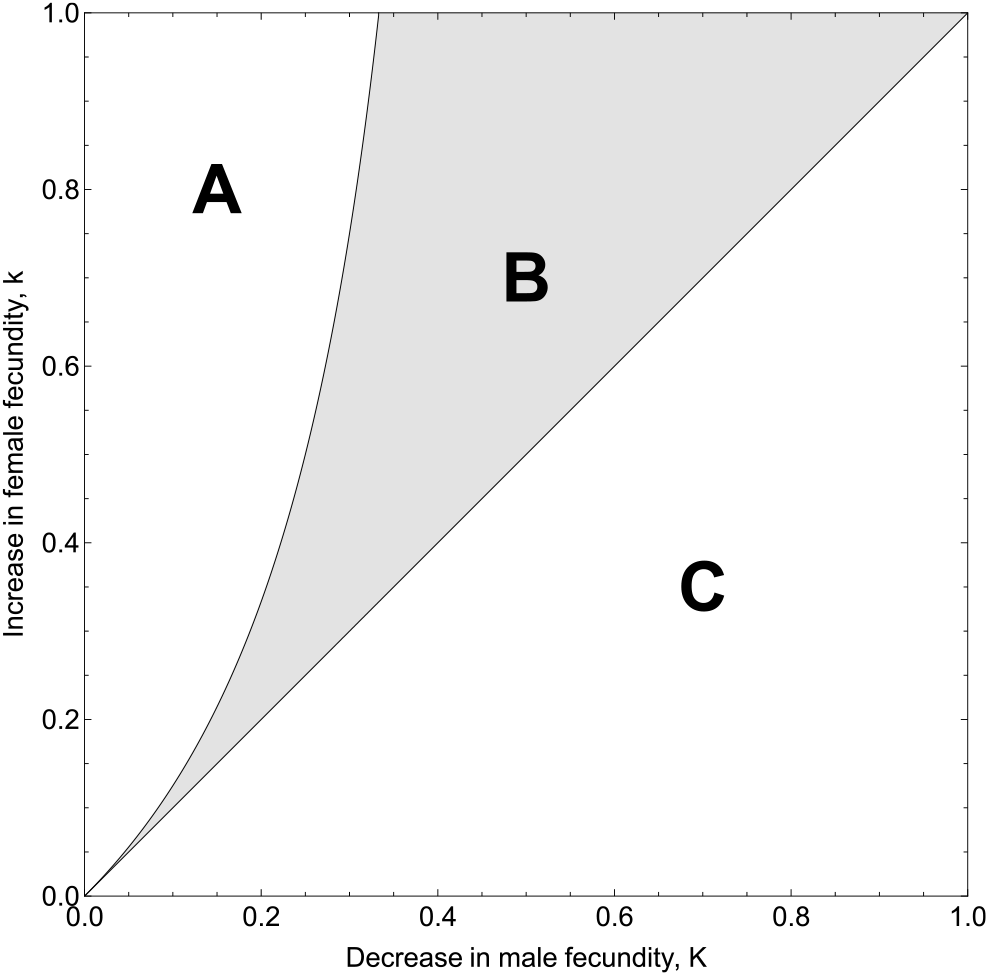
Polymorphism conditions in Charlesworth and Charlesworth (1978b) under complete outcrossing, as a function of the increase in female fecundity *k* and the decrease in male fecundity *K* (from eq. B36). **A** Fixation of the female-biased mutant (*W*_f_ > 1 and *W*_0_ < 1); **B** Genetic polymorphism (*W*_f_ > 1 and *W*_0_ > 1); **C** Exclusion of the female-biased mutant (*W*_f_ < 1 and *W*_0_ > 1).

**Figure S8:**
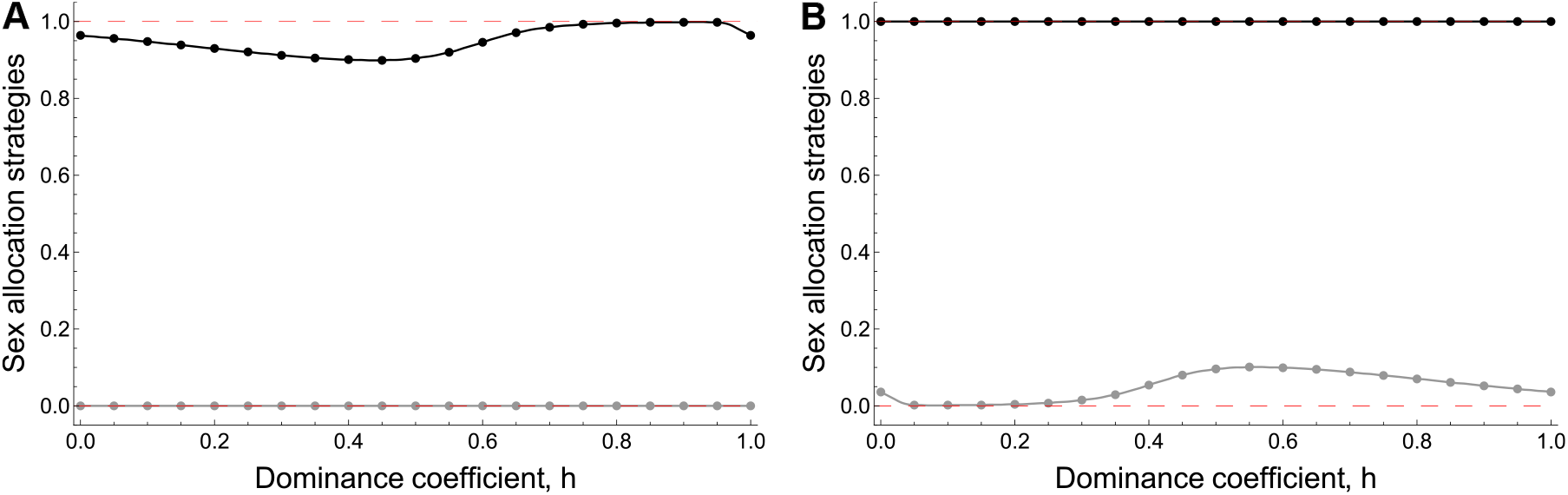
Equilibrium sex allocation strategies 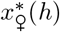 (black) and 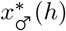 (grey), encoded by allele *x*_♀_ and *x*_♂_ as a function of the dominance coefficient *h*. Dashed red lines indicate 0 and 1. **A** Androdioecy, *γ*_♀_ = 2 and 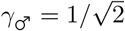. **B** Gynodioecy, 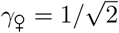 and *γ*_♂_ = 2. See section C.2.3 for analysis.

**Figure S9:**
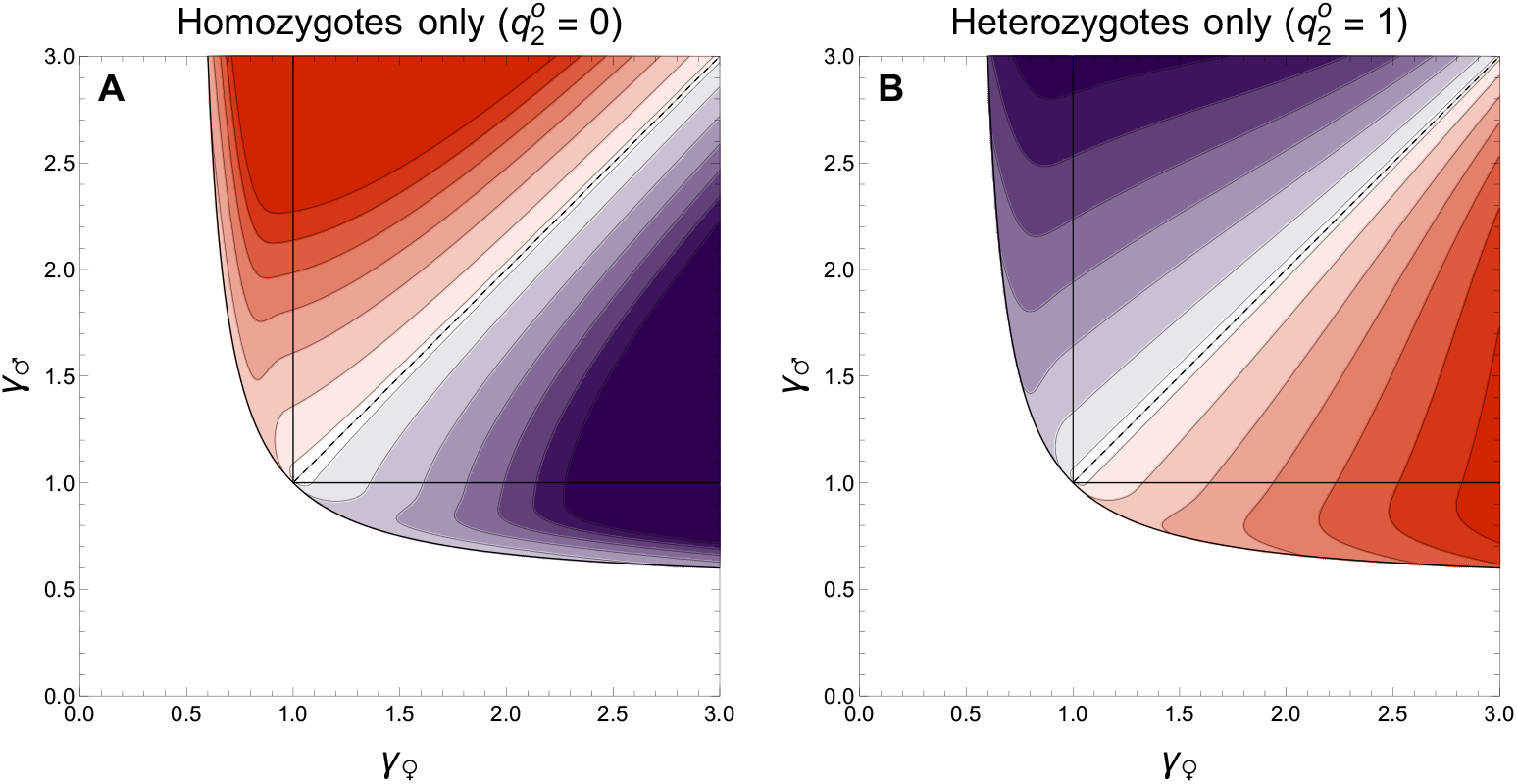
A Selection gradient on dominance at additivity (*h* = 1*/*2) assuming the resident population comprises only homozygotes. **B** Selection gradient on dominance at additivity (*h* = 1*/*2) assuming the resident population comprises only heterozygotes. Orange indicates a positive selection gradient favouring ZW sex determination (*h* →1) and purple indicates a negative selection gradient favouring XY sex determination (*h* → 0). The darker the colour, the more intense selection is.

**Figure S10:**
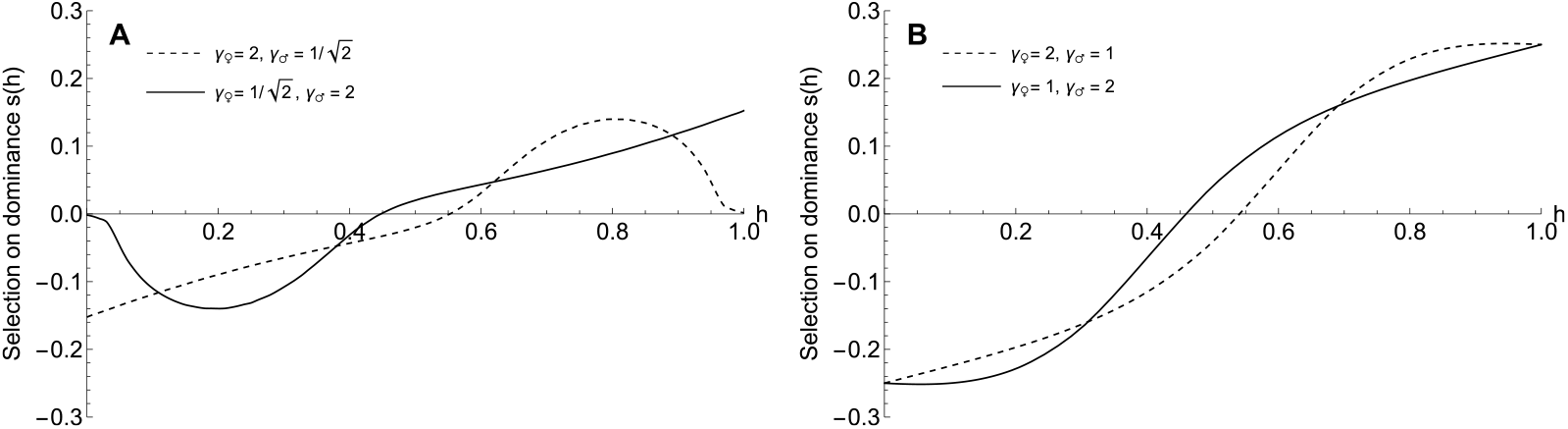
Selection gradient on dominance in four cases, **A** Androdioecy (dashed line) and Gynodioecy (solid line), **B** Dioecy with *γ*_♀_ *> γ*_♂_ (dashed line) and *γ*_♀_ *< γ*_♂_ (solid line).

**Figure S11:**
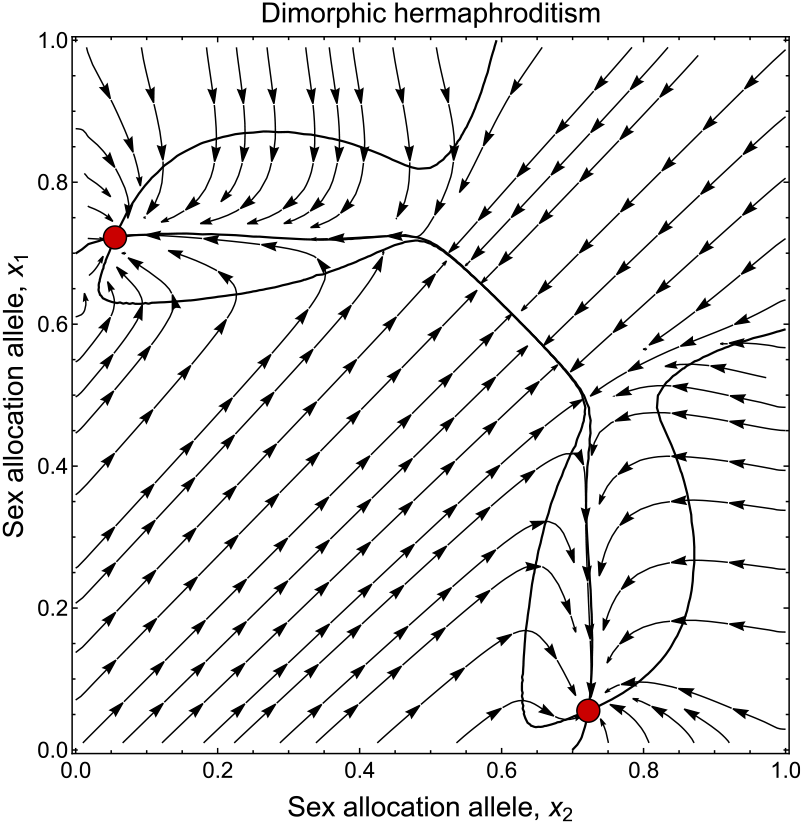
Evolutionary dynamics of sex allocation under high inbreeding depression (*δ* = 0.75) with saturating gain curves (*γ*_♀_ = *γ*_♂_ = 0.83) leading to dimorphic hermaphroditism. Other parameters: *α*_0_ = 0.75 and *β* = 1. Approach is explained in section D.2.3.

**Figure S12:**
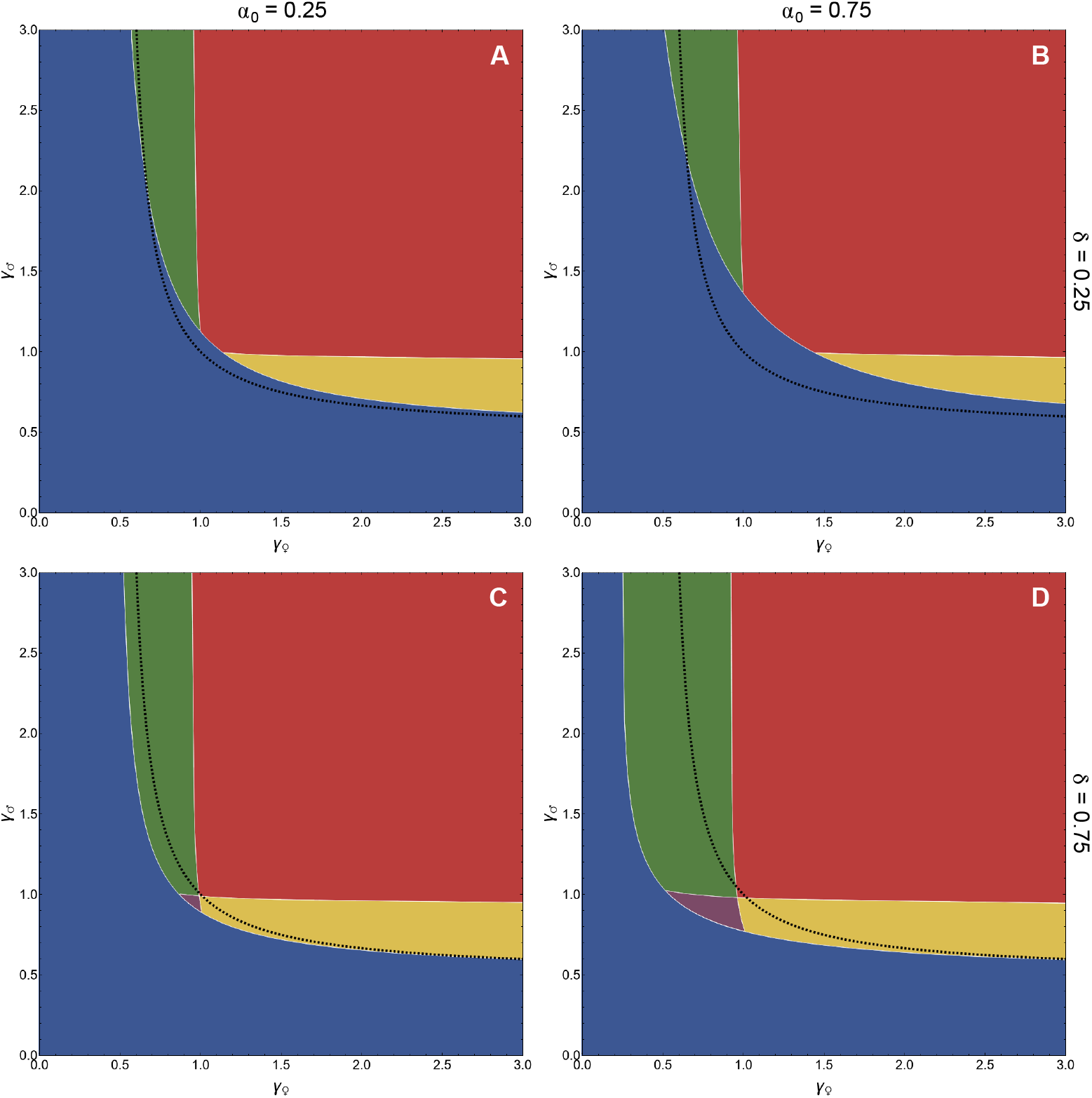
Parameter space corresponding to the possible outcomes of gradual evolution under low and high inbreeding depression (*δ*) and selfing rates (*α*_0_). The region in which monomorphic hermaphroditism is maintained is indicated in blue, and the region in which selection favours dimorphic hermaphroditism is shown in purple. The regions in which androdioecy, gynodioecy and dioecy are favoured are shown in yellow, green and red, respectively. The black dashed line indicates the limit above which polymorphism is favoured in the strictly outcrossing case (so when *α*_0_ = 0). Other fixed parameters: *β* = 1. Approach is explained in section D.2.3.

**Figure S13:**
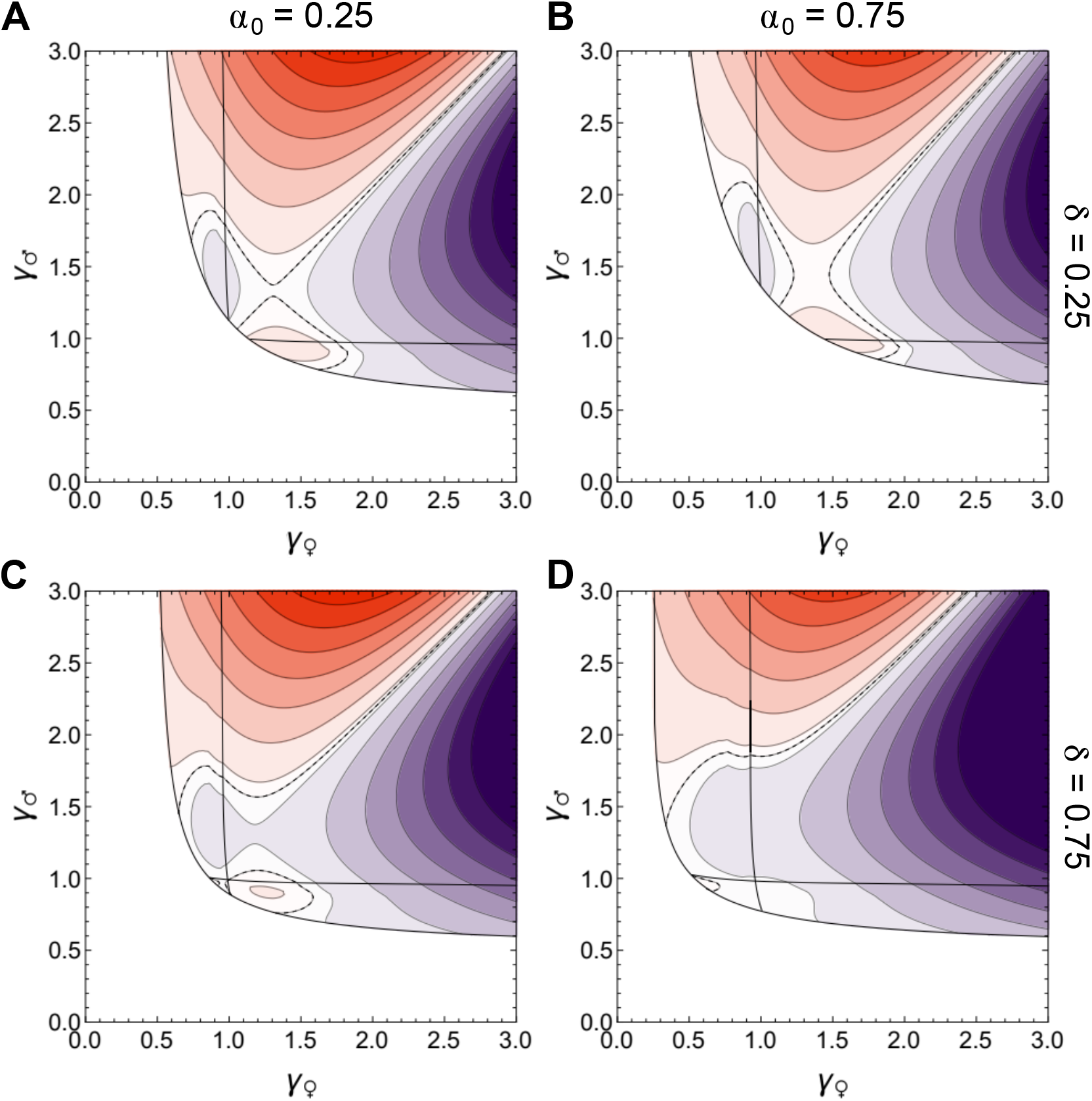
Selection gradient on dominance at additivity (eq. D48 with *h* = 1*/*2) for four representative combinations of selfing rate (*α*_0_) and inbreeding depression (*δ*). Orange shades indicate a positive selection gradient, favouring ZW sex determination, and dark purple shades indicate a negative gradient, favouring XY. The darker the colours, the more intense selection.

**Figure S14:**
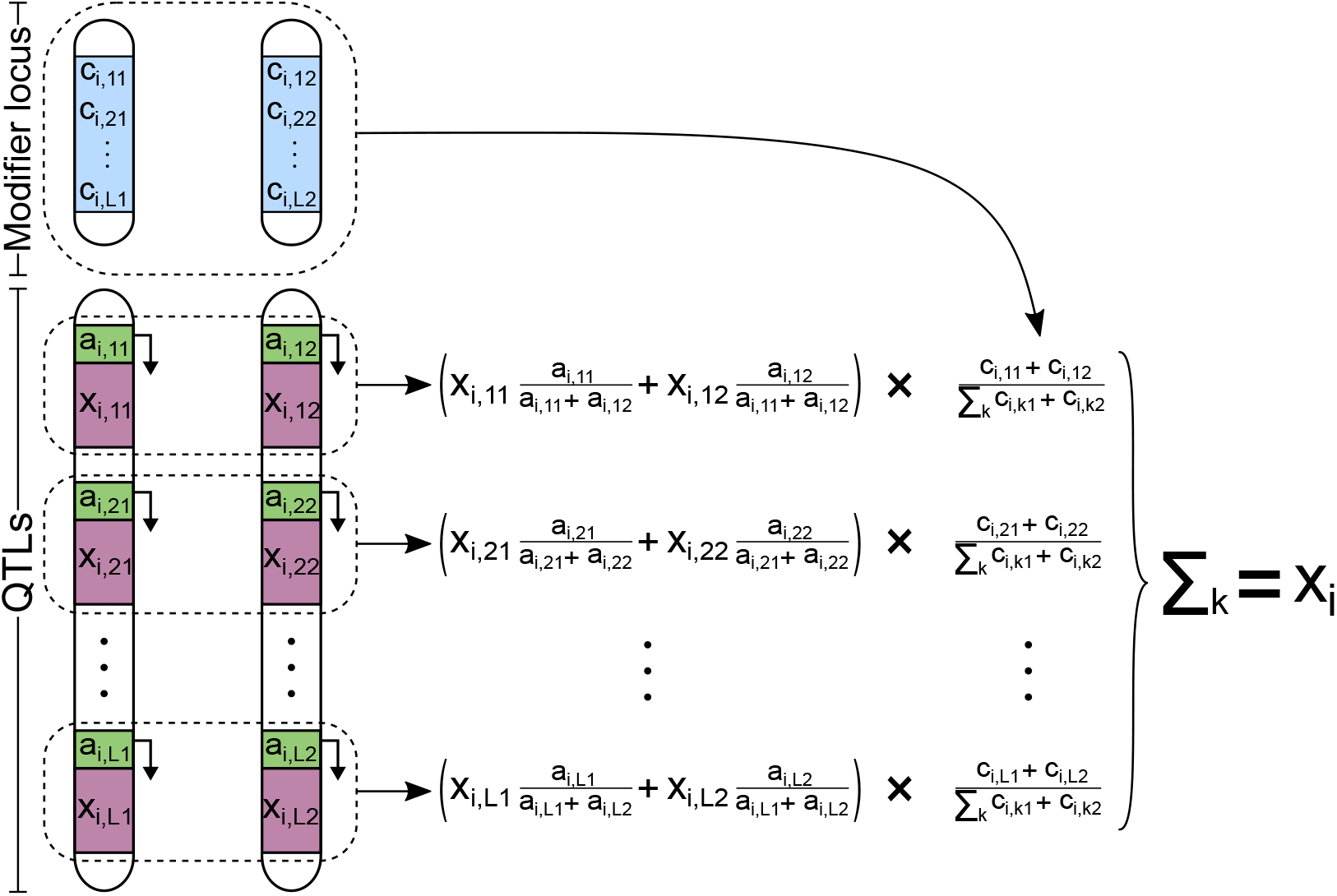
Genetic architecture assumed in the simulations. The *L* sex allocation loci are shown in green (promoter) and pink (gene), and the modifier is shown in blue. Each sex allocation locus encodes a sex allocation strategy (in brackets), and the strategy *x*_*i*_ expressed by the individual is given by the sum of the strategies encoded by the *L* loci, weighted by their relative contributions given by the modifier.

